# Comprehensive glycoprofiling of oral tumours associates *N*-glycosylation with lymph node metastasis and patient survival

**DOI:** 10.1101/2022.11.21.517331

**Authors:** Carolina Moretto Carnielli, Thayná Melo de Lima Morais, Fábio Malta de Sá Patroni, Ana Carolina Prado Ribeiro, Thaís Bianca Brandão, Evandro Sobroza, Leandro Luongo Matos, Luiz Paulo Kowalski, Adriana Franco Paes Leme, Rebeca Kawahara, Morten Thaysen-Andersen

**Author notes:** Joint corresponding authors. These authors contributed equally to this work (Rebeca Kawahara, Morten Thaysen-Andersen).

## Abstract

While altered protein glycosylation is regarded a trait of oral squamous cell carcinoma (OSCC), its heterogeneous glycoproteome and dynamics with disease progression remain unmapped. To this end, we here employ an integrated multi-omics approach comprising unbiased and quantitative glycomics and glycoproteomics applied to a valuable cohort of resected tumour tissues from OSCC patients with (n = 19) and without (n = 12) lymph node metastasis. While all tumour tissues displayed uniform *N*-glycome profiles suggesting relatively stable global *N*-glycosylation during lymph node metastasis, glycoproteomics and advanced correlation analysis notably uncovered altered site-specific *N*-glycosylation and previously unknown associations with several key clinicopathological features. Importantly, focused analyses of the multi-omics data unveiled two *N*-glycans and three *N*-glycopeptides that were closely associated with patient survival. This study provides novel insight into the complex OSCC tissue *N*-glycoproteome forming an important resource to further explore the underpinning disease mechanisms and uncover new prognostic glyco-markers for OSCC.

**Teaser:** Deep survey of the dynamic landscape of complex sugars in oral tumours paves a way for new prognostic disease markers.

## Introduction

Oral squamous cell carcinoma (OSCC) is the most common type of head and neck cancer (*1*). With more than 377,000 new cases and 170,000 deaths annually worldwide, OSCC is an aggressive disease with a dishearteningly low five-year survival rate of 50% (*2, 3*), mainly due to lymph node metastasis and loco-regional failures. OSCC prognostication is currently based on the clinical staging system of tumour-lymph node-metastasis (TNM system) of the disease. However, this system has several flaws, as patients assigned with the same TNM stage may present with different clinical features and experience different outcomes (*4, 5*). Thus, identification of molecular signatures that may assist in a more precise staging and prognosis of patients with OSCC is urgent needed.

Histopathological analysis of formalin-fixed paraffin-embedded (FFPE) tissues remains the principal method for diagnosis and prognosis of OSCC patients (*6, 7*). FFPE tissue slides preserve the cellular morphology and molecular features of the tumour tissues, and have therefore been explored as a source for biomarker investigations (*8, 9*). We recently used sensitive mass spectrometry (MS)-based proteomics to map different areas of FFPE tissues from OSCC patients, which revealed key proteins with potential prognostic value as demonstrated by their associations with various clinicopathological parameters; the prognostic value of these proteins was also demonstrated by targeted proteomics of saliva from OSCC patients with and without lymph node metastasis (*10*). Our findings also revealed that specific glycoproteins i.e. ITGAV and COL6A1 from the tumour stroma, and COL1A2 from tumour cells associate with lymph node metastasis and type of disease treatment, which collectively point to the involvement of protein glycosylation in OSCC and suggest that glycoprofiling may augment OSCC prognostication.

Despite the growing list of candidate biomarkers reported for oral cancer, none has to date been implemented to aid the clinical decision-making, but further evaluation studies are underway and may eventually provide a path towards clinically robust biomarkers for improved disease management (*11*).

Evidence is emerging that not only the protein abundance, but also the glycosylation patterns and levels of the underlying glycosylation enzymes, are altered in OSCC (*12, 13*). Moreover, glycosylation changes have previously been associated with altered adhesion behaviour, migration and metastasis of oral cancer cells as well as OSCC disease progression (*13–15*). Aberrant *N*-glycosylation of E-cadherin, a key marker of epithelial mesenchymal transition in cancer development (*16*), and reduced adhesion of human salivary epidermoid carcinoma cells were found to result from the overexpression of dolichol-P-dependent *N*-acetylglucosamine-1-phosphate transferase (DPAGT1), an enzyme that initiates the synthesis of the lipid-linked oligosaccharide precursor for protein *N*-glycosylation in the endoplasmic reticulum (*17*). Overexpression of DPAGT1 detected in specimens from resected OSCC tumours in the oral cavity was also found to be associated with an aberrant activation of the canonical Wnt signalling pathway that contributes to the development and progression of many human cancers (*14, 18, 19*). Moreover, site-specific glycoprofiling of a gingival carcinoma cell line uncovered elevated sialylation and reduced fucosylation relative to a non-cancerous oral epithelial cell line from gingiva (*20*). The study also revealed that B7-H3 (also known as B7 homolog 3 or CD276 isoform 1) knockdown suppresses tumour cell proliferation, and that B7-H3 restoration enhances tumour growth. Finally, elevated sialidase activity was observed in saliva of oral cancer patients as compared to healthy individuals and patients with precancerous conditions (*21*). While these studies collectively indicate that aberrant protein glycosylation is linked to the onset and progression of oral cancer, altered glycosylation has not previously been associated with patient prognosis including lymph node metastasis status and survival outcome.

Herein, we employ an integrative glyco-centric multi-omics approach to quantitatively profile the heterogenous *N*-glycoproteome in tumour tissues from OSCC patients with (N+) or without (N0) lymph node metastasis to establish the glycan fine structures, their site-specific microheterogeneity, and their disease dynamics. Our study is the first to unpick the OSCC *N*-glycoproteome; we report on novel associations between altered *N*-glycosylation and key clinical features including lymph node metastasis and patient survival, and provide a comprehensive resource to further explore the underpinning disease mechanisms and uncover new disease markers for OSCC.

## Results

### Comprehensive *N*-glycoprofiling of OSCC tumours from patients with and without lymph node metastasis

This study applied an integrative MS-driven multi-omics approach comprising both quantitative glycomics and glycoproteomics to comprehensively profile the protein *N*-glycosylation in FFPE tumour tissues surgically removed from 31 patients operated with curative intent due to OSCC, including patients with (N+, n = 19) and without (N0, n = 12) lymph node metastasis, Figure 1 and Supplementary Figure S1. See Supplementary Table S1-S2 for patient metadata and overview of the generated LC-MS/MS datasets, respectively.

**Figure 1.**
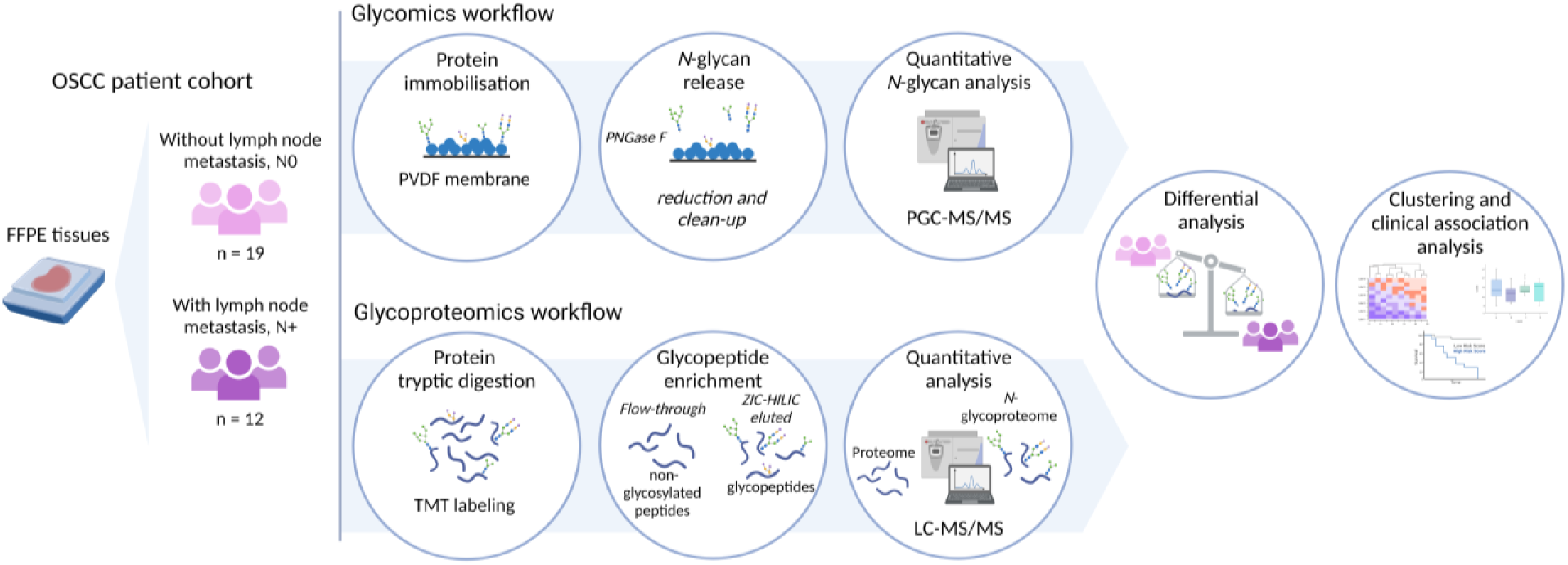
Study overview. Protein *N*-glycosylation was investigated using quantitative glycomics (top) and glycoproteomics (bottom) from resected tumour tissues from 31 OSCC patients with (N+, n = 19) or without (N0, n = 12) lymph node metastasis. The resulting *N*-glycome and *N*-glycoproteome data were quantitatively compared between patient groups using advanced statistical tests including clustering analysis and associations to a range of clinicopathological features were explored using patient metadata.

### Comparative *N*-glycome profiling of OSCC tissues from N+ and N0 patients

State-of-the-art quantitative *N*-glycomics of proteins extracted from the OSCC tumour tissues was used to profile, with glycan fine structural resolution, a total of 83 *N*-glycan structures spanning 52 *N*-glycan compositions (denoted Glycan 1-52 with isomers denoted a, b, c…). See Supplementary Figure S2 for a map of the identified *N*-glycan structures and Supplementary Table S3 and Supplementary File S1 for tabulated glycomics data and spectral evidence, respectively.

Mainly complex (38.7-61.5%), oligomannosidic (18.3-36.5%) and paucimannosidic (9.8-31.9%) *N*-glycans were identified across the OSCC tumour tissues, Figure 2A and Supplementary Figure S3. Despite minor fold-change differences between N+ and N0 patients (−0.82 to 0.94), no prominent differences were observed in the *N*-glycan type distribution nor in the common *N*-glycan structural features such as the degree of branching and levels of fucosylation and sialylation (all *p* ≥ 0.05), Figure 2B. Uniform global *N*-glycosylation across the N0 and N+ tissues was further demonstrated using pair-wise *N*-glycome correlation analysis, PCA and hierarchical clustering analysis, Supplementary Figure S4 and Supplementary Table S4.

**Figure 2.**
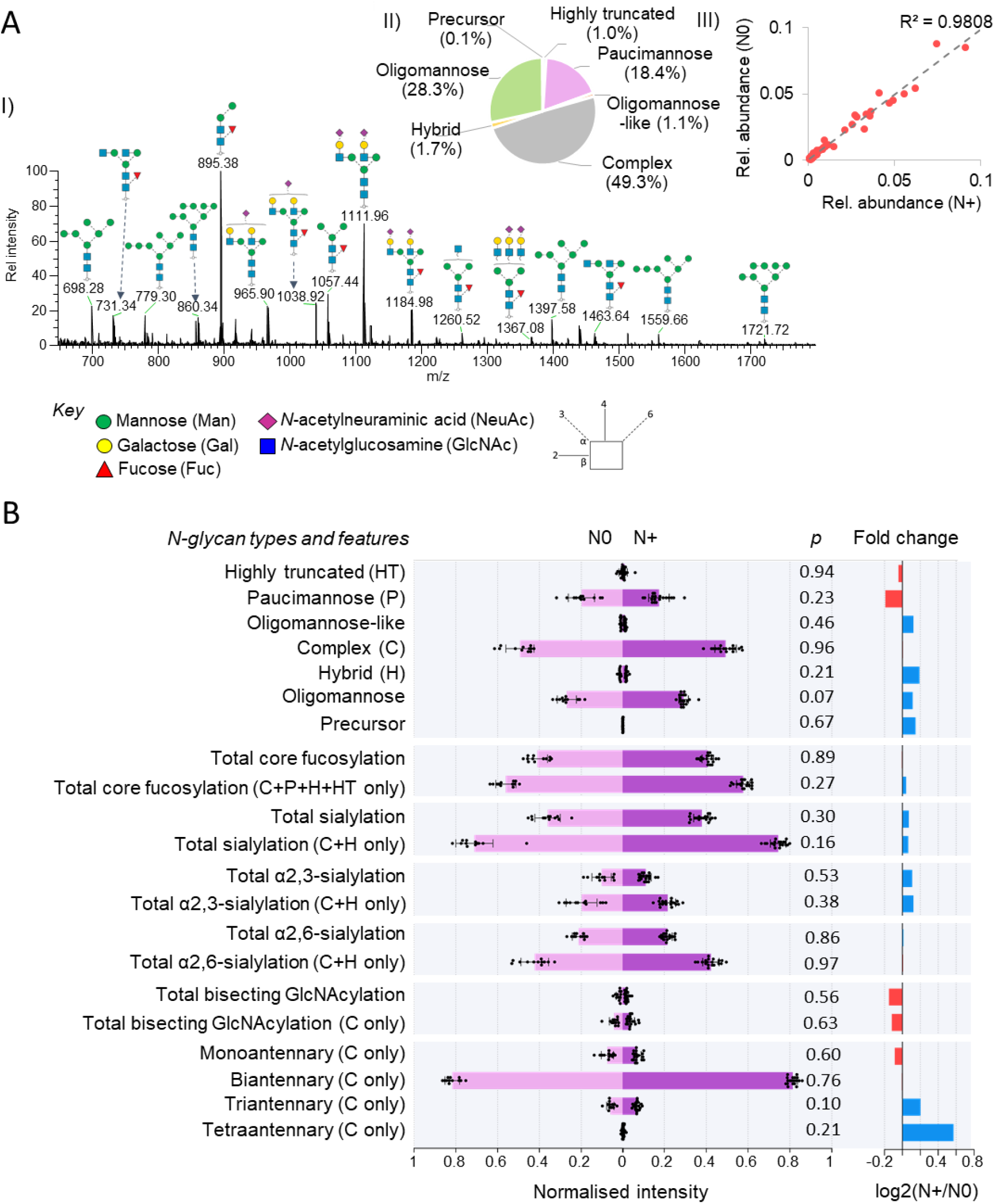
Stable *N*-glycome in OSCC tumour tissues. A) I- The OSCC tumour tissue *N*-glycome predominantly comprises complex, oligomannosidic and paucimannosidic *N*-glycans as shown by an exemplar summed MS1 spectrum. II- Overall distribution of *N*-glycan classes. III- Similar *N*-glycan distribution within N+ and N0 tissues as demonstrated by a high correlation coefficient (R^2^). B) Quantitative comparison of *N*-glycan types and other key structural features between N0 and N+ including total core fucosylation levels (calculated both out of the entire *N*-glycome and out of only structures able to carry core fucosylation) and total sialylation levels including α2,3- or α2,6-sialylation, bisecting GlcNAcylation and degree of branching (mono-, bi- or triantennary *N*- glycosylation). While minor fold-change differences were observed (right), no significant differences were consistently found between N0 and N+. HT: highly truncated, P: paucimannose, H: hybrid, C: complex, see Supplementary Figure S2 for structures and classification. Data are plotted as means and error bars represent their standard deviation (n = 19, N+ and n = 12, N0). Statistical test: two-sided Student’s t test.

Despite notable *N*-glycome similarities, differential expression was observed for six individual *N*-glycan structures (*p* < 0.05) including one down-regulated *N*-glycan (Glycan 34) and five up-regulated *N*-glycans (Glycan 20a, Glycan 40a, Glycan 45b, Glycan 46a, Glycan 49b) in N+ compared to N0 tissues, Supplementary Table S3. Interestingly, these differentially expressed *N*-glycans were able to accurately stratify the N0 and N+ patients with confidence using a logistic regression model (AUC values ranging from 67.5-85.5%), and, in part, also by a random forest model (AUC values from 49.1% to 70.2%), Supplementary Figure S5.

### Comparative *N*-glycoproteome profiling of OSCC tumour tissues from N+ and N0 patients

Guided by the acquired *N*-glycomics data, we then explored the glycoproteome complexity of the OSCC tumour tissues using our recently developed glycomics-informed glycoproteomics method (*22*). A total of 3,117 unique *N*-glycopeptides (i.e. glycopeptides carrying a discrete glycan composition at a discrete site) displaying 55 different *N*-glycan compositions from 419 different source *N*-glycoproteins spanning a wide dynamic range were profiled across the OSSC tissue cohort, Figure 3A and Supplementary Table S5-S6.

**Figure 3.**
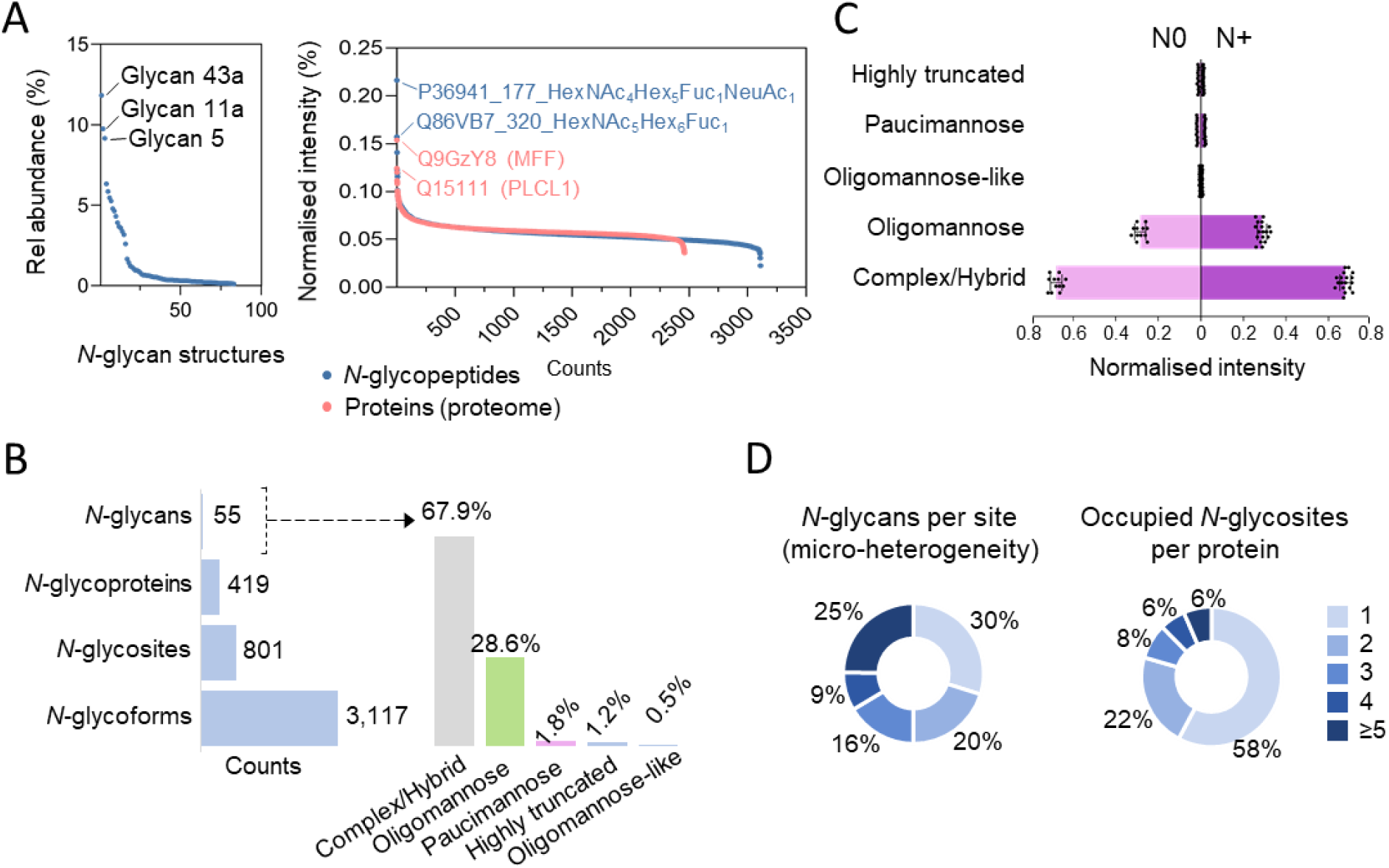
Overview of the *N*-glycoproteome profile of OSCC tumour tissues. A) Abundance range of identified *N*-glycan compositions (left) and identified *N*-glycopeptides (both from the acquired *N*-glycoproteome data) and proteins (from the proteome data). The most abundant *N*-glycans, *N*-glycopeptides and proteins are labelled in each graph. B) Overview of the *N*-glycoproteome of the investigated OSCC tumor tissues including the *N*-glycan compositions (and their distribution across the *N*-glycan classes, right), source *N*-glycoproteins, *N*-glycosites, and unique *N*-glycoforms (unique protein + unique site + unique glycan) identified in the ZIC-HILIC-enriched fractions. C) Comparison between the distribution of *N*-glycan classes in the N0 and N+ tumour tissues. D) Site-specific *N*-glycan micro-heterogeneity (top) and occupied sites per protein (bottom) in the OSCC tumour tissues. HT: highly truncated, P: paucimannose, O: oligomannose, O-L: oligomannose-like, C/H: complex and hybrid (grouped since these cannot reliably be distinguished through glycoproteomics data).

In line with the *N*-glycome data, most *N*-glycopeptides carried complex/hybrid (67.9%) or oligomannosidic (28.6%) *N*-glycans, Figure 3B. The under-representation of paucimannosidic *N*-glycopeptides in the glycoproteomics data (1.8%) relative to the considerable levels found in the *N*-glycome (18.4%) can likely be attributed to inefficient HILIC-SPE enrichment of these less hydrophilic *N*-glycopeptides (unpublished observation). Recapitulating findings from the *N*-glycome data, the glycoproteomics data did not reveal any consistent differences in the *N*-glycan type distribution, Figure 3C, and the global *N*-glycoproteome, Supplementary Figure S6, between the N0 and N+ patients. Providing further insights into the OSCC tumour tissue *N*-glycoproteome complexity, the glycoproteomics data also revealed that most *N*-glycosites (70%) carried more than one discrete *N*-glycan composition (25% of sites were decorated with ≥5 glycan compositions) while most *N*-glycoproteins (58%) were identified with only one occupied *N*-glycosylation site, Figure 3D.

Despite the relatively uniform *N*-glycoproteome across the N0 and N+ patients, a total of 79 *N*-glycopeptides from 56 source *N*-glycoproteins were found to be quantitatively altered between the two patient groups (*p* < 0.05) including 57 *N*-glycopeptides from 43 source *N*-glycoproteins that were elevated and 22 *N*-glycopeptides from 14 source *N*-glycoproteins of lower abundance in N+ relative to N0 tissues, Supplementary Table S5. Excitingly, all 79 *N*-glycopeptides demonstrated a notable potential to stratify N0 and N+ OSCC patients (AUC-ROC >60% by logistic regression model), Supplementary Table S7. In support, a high proportion of these *N*-glycopeptides (50 *N*-glycopeptides, 63.3%) were also found to stratify the patient groups with an AUC-ROC >60% using a random forest model.

### *N*-glycosylation-guided clustering of OSCC patients

To further investigate the relatively subtle yet consistent differences in *N*-glycosylation found within the investigated OSCC tumour tissue cohort, the obtained *N*-glycome and *N*-glycoproteome data were then interrogated using advanced data analysis and visualisation methods.

Unsupervised hierarchical clustering of the *N*-glycome data revealed two major tumour clusters (T-C1 and T-C2) and two major *N*-glycan clusters (NG-C1 and NG-C2), Figure 4A. Within the NG-C1 cluster, altered distribution of all *N*-glycan classes were found between T-C1 and T-C2, while in NG-C2 differences in the distribution of paucimannosidic and complex *N*-glycans, but not oligomannosidic *N*-glycans, were observed between the two tumour clusters (*p* < 0.05) (note that the other glycan classes were not observed in NG-C2), Figure 4B.

**Figure 4.**
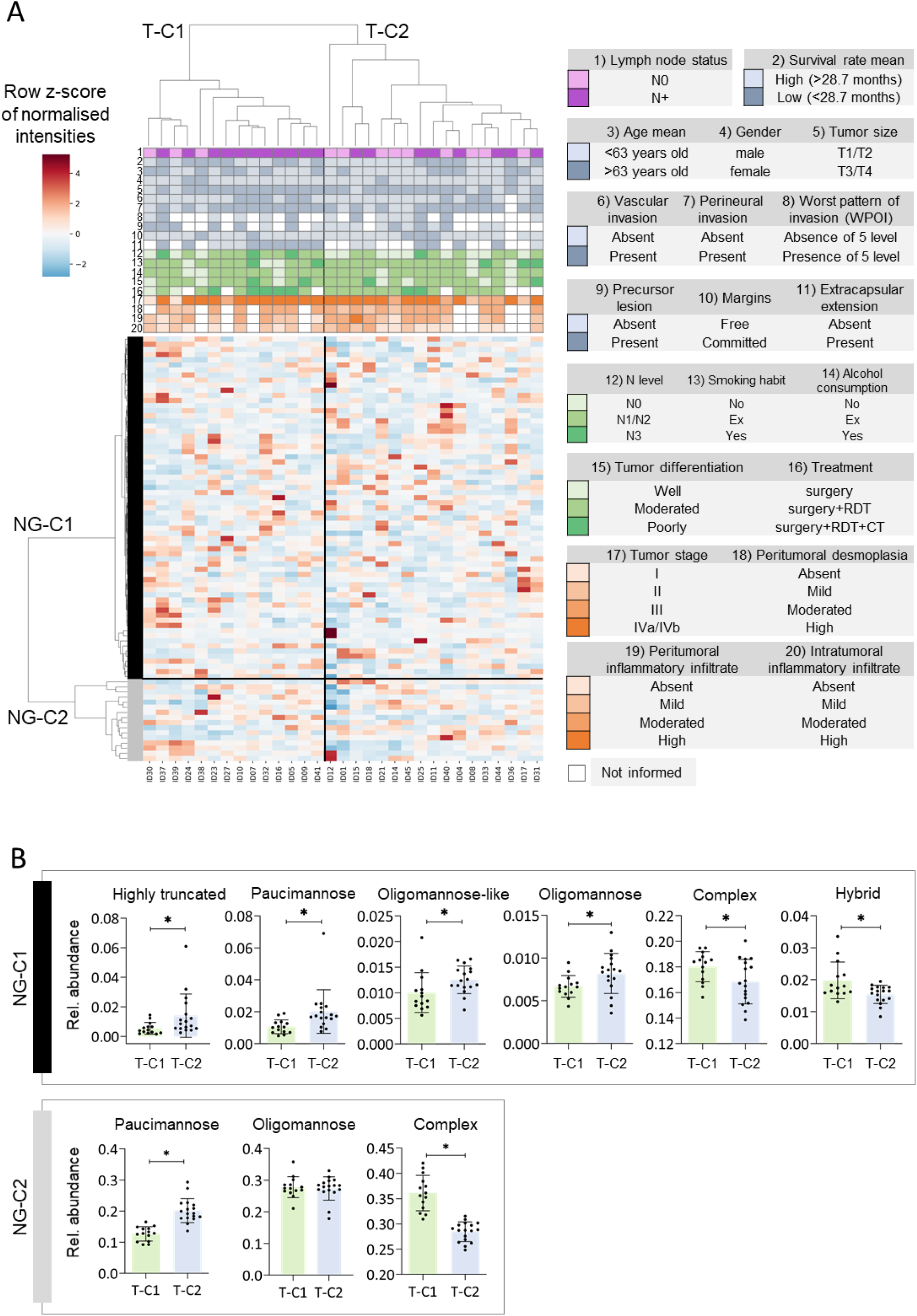
*N*-glycome-driven clustering of OSCC tumour tissues. A) Unsupervised hierarchical clustering analysis of the OSCC tumour tissue *N*-glycome data from N0 and N+ patients, performed with the ‘cluster map’ function in the Saborne package under Python using Euclidean distance and Ward linkage. Two major tumour clusters (T-C1 and T-C2) and two major *N*-glycan clusters (NG-C1 and NG-C2) were observed. Clinical and patient data are presented on top of the heat map and expanded to the right. B) Relative abundance of the *N*-glycan types between T-C1 (green bars) and T-C2 (blue bars) in NG-C1 (top) and NG-C2 (bottom), respectively. Student’s t test, two-sided, α = 0.05, \**p* < 0.05, \*\**p* < 0.01, \*\*\**p* < 0.001.

Similarly, unsupervised hierarchical clustering of the *N*-glycoproteome data revealed two main tumour clusters (T-C1 and T-C2) and two *N*-glycopeptide clusters (IG-C1 and IG-C2), Figure 5A. Differences in expression of highly truncated, oligomannosidic and complex/hybrid *N*-glycans were identified between T-C1 and T-C2 in IG-C1, while only the highly truncated *N*-glycans exhibited differences between the two tumour clusters in IG-C2, Figure 5B.

**Figure 5.**
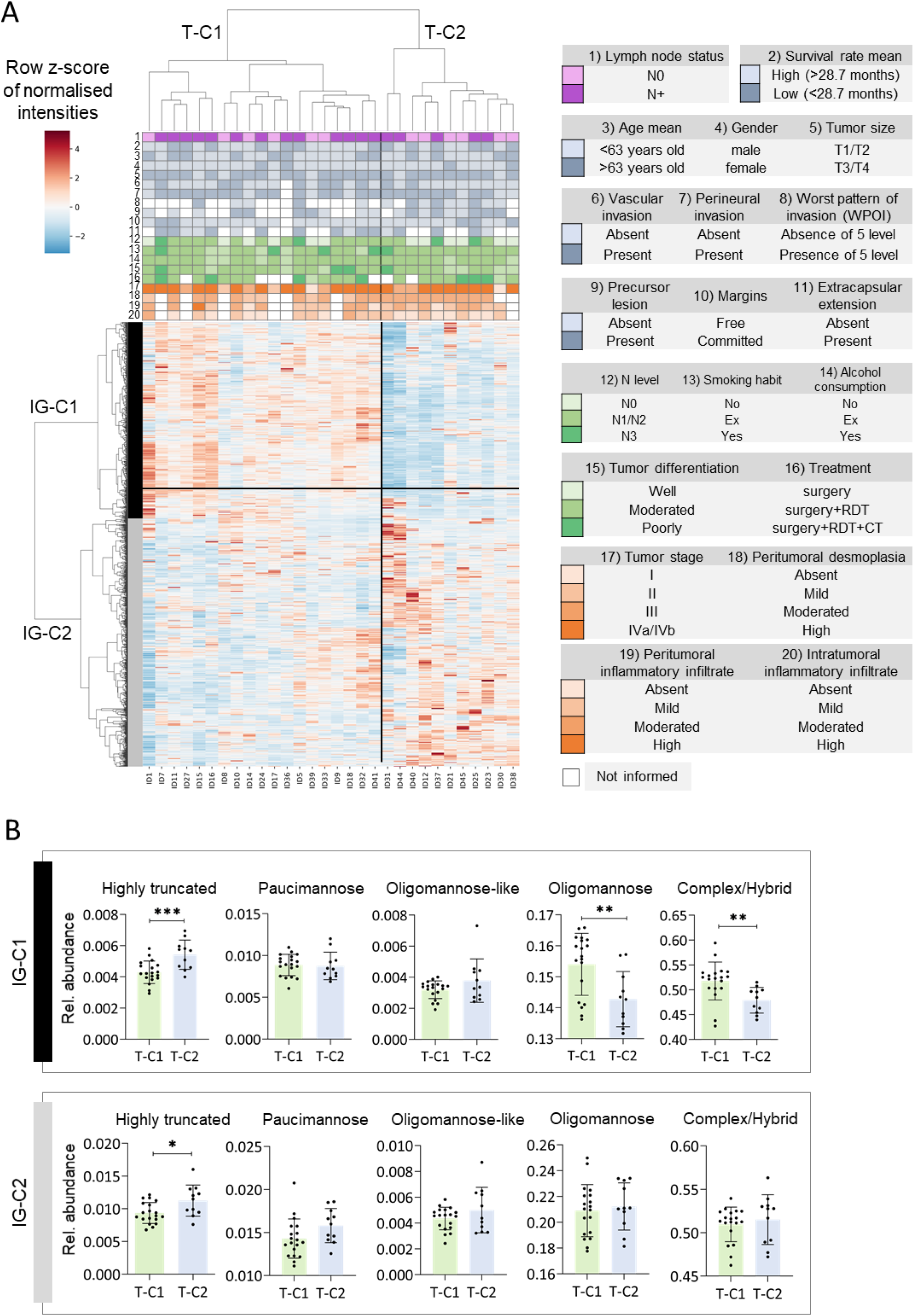
*N*-glycoproteome-driven clustering of OSCC tumour tissues. A) Unsupervised hierarchical clustering analysis of *N*-glycopeptides identified in the tumor tissues from N0 and N+ patients, performed with the ‘cluster map’ function in the Saborne package under Python using Canberra distance and Ward linkage. Two major tumor clusters i.e. T-C1 and T-C2 and two major *N*-glycopeptide clusters i.e. IG-C1 and IG-C2 were observed. Clinical and patient data are presented on top of the heat map, and expanded to the right. B) Relative abundance of *N*-glycan classes between tumor clusters (T-C1 in green, and T-C2 indicated in blue) in IG-C1 (top) and IG-C2 (bottom). Student’s t test, two-sided, α = 0.05, \**p* < 0.05, \*\**p* < 0.01, \*\*\**p* < 0.001.

To explore potential biological roles of the source *N*-glycoproteins identified within each *N*-glycopeptide cluster, pathway enrichment analyses were performed. Extracellular matrix organisation was the most enriched biological process in both *N*-glycopeptide clusters (FDR adjusted *p* = 9.4 x 10^-36^ in IG-C1 and *p* = 3.6 x 10^-45^ in IG-C2), followed by platelet degranulation in IG-C1 (FDR adjusted *p* = 1.1 x 10^-20^) and neutrophil degranulation in IG-C2 (FDR adjusted *p* = 2.0 x 10^-21^) when comparing T-C1 and T-C2, Supplementary Figure S7A. We then tested for differential *N*-glycan class distribution within these highly enriched pathways, which demonstrated altered expression of highly truncated *N*-glycans decorating proteins involved with platelet degranulation within IG-C1, and showed altered pauci- and oligomannosidic *N*-glycosylation on proteins involved in neutrophil degranulation within IG-C2, Supplementary Figure S7B.

### Associations between *N*-glycoproteome components, clinical features and biological processes

The six *N*-glycans and 79 *N*-glycopeptides found to be differentially expressed in N0 and N+ patients were then evaluated for associations with a range of clinical features.

Unsupervised hierarchical clustering of the expression data of the six *N*-glycan structures revealed two new distinct tumour groups that displayed differences in vascular invasion (*p* = 0.049, Fisher’s Exact Test) and lymph node (N) status (*p* = 0.039, Pearson Chi-Square test), Figure 6A-B. In contrast, the two tumour groups formed by the 79 *N*-glycopeptides differed in terms of N status (*p* = 0.0078), tumour size (*p* = 0.0169, both Fisher’s Exact Test) and type of treatment received by the OSCC patient (*p* = 0.0354, Pearson Chi-Square), Figure 6C-D.

**Figure 6.**
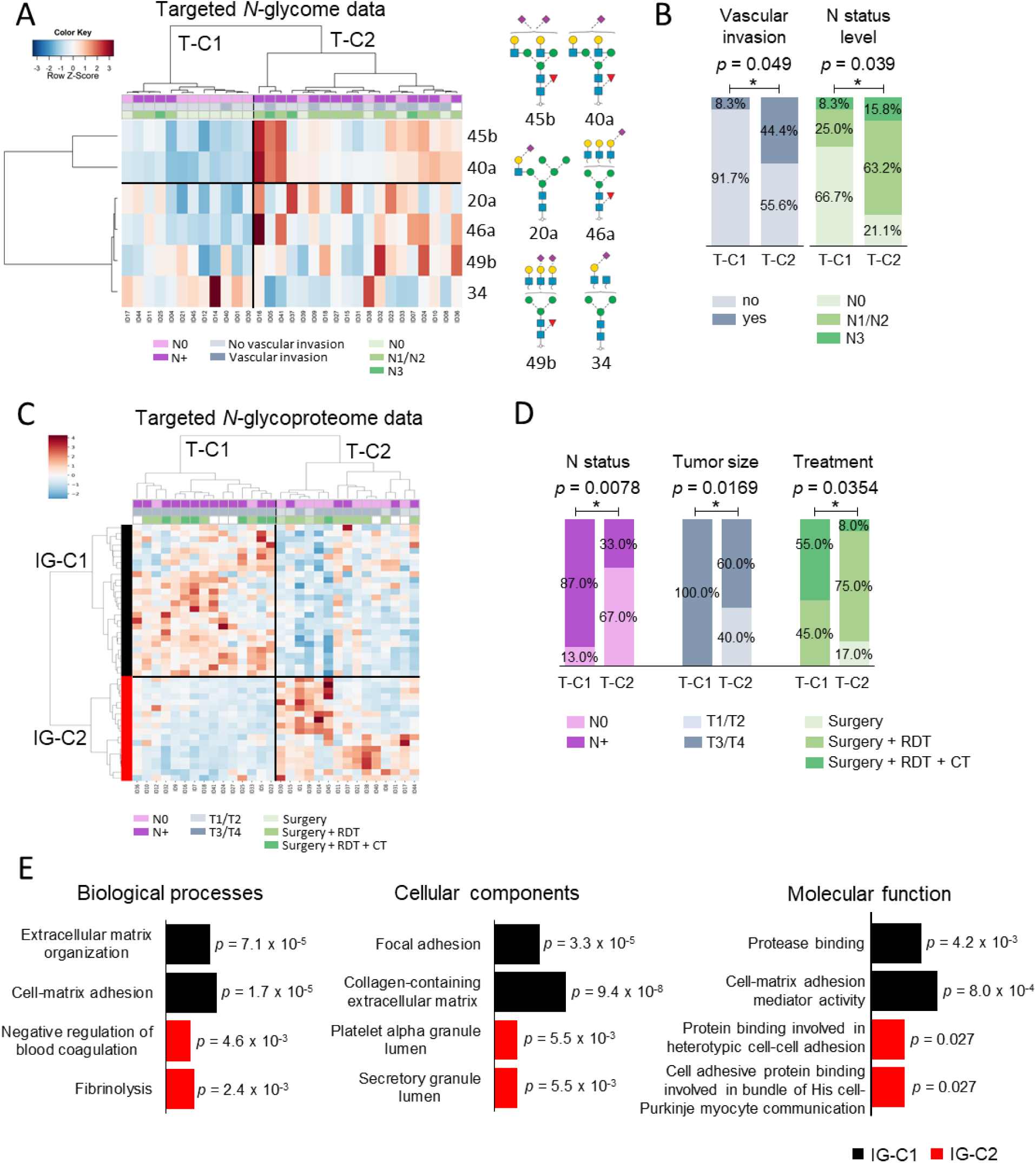
*N*-glycan and *N*-glycopeptide-guided tumour clusters associate with clinical features. A) Unsupervised hierarchical clustering analysis of six *N*-glycans found to be differentially abundant in OSCC tumour tissues from N0 and N+ patients. Plots are made with the ‘heatmap.3’ function under the R environment using Euclidean distance and Ward linkage. B) Cluster association analysis with clinical features (*p* < 0.05, Fisher’s Exact Test for two group comparisons; or Pearson Chi-Square test for more than two group comparisons) performed using IBM SPSS Statistics. C) Unsupervised hierarchical clustering analysis of 79 *N*-glycopeptides found to be differentially expressed between the N0 and N+ tumour tissues. Plots were performed with the ‘cluster map’ function in the Saborne package under Python using Ward and correlation for clustering. D) Cluster association analysis with clinical features. E) The two most enriched Gene Ontology biological processes, cellular components and molecular function in the IG-C1 (black) and IG-C2 (red) glycopeptide clusters. An adjusted *p* value is indicated for each term in the enrichment analysis performed using the Enrichr tool. Fisher’s Exact Test or Pearson Chi-Square test, \**p* < 0.05. RDT: radiotherapy; CT: chemotherapy.

To investigate whether *N*-glycosylation was associated to specific biological processes, cellular compartments and molecular function, we searched for enriched gene ontology (GO) terms within the IG-C1 and IG-C2 *N*-glycopeptide clusters. This analysis revealed significant enrichment of extracellular matrix organisation and cell-matrix adhesion in IG-C1, while ‘negative regulation of blood coagulation’ and ‘fibrinolysis’ were enriched in IG-C2, Figure 6E. Proteins from IG-C1 were enriched in ‘focal adhesion’ and ‘collagen-containing extracellular matrix’ compartments, and were involved in ‘protease binding’ and ‘cell-matrix adhesion mediator activity’, whereas proteins from IG-C2 showed trends of being of platelet and secretory granule origins, and involved in protein binding.

### Specific OSCC *N*-glycans and *N*-glycopeptides associate with key clinical outcomes

In total 25 of the 79 *N*-glycopeptides that displayed differential abundance between N0 and N+ patients were found to correlate with distinct clinicopathological features, Supplementary Figure S8. Amongst them were five *N*-glycopeptides all carrying Glycan 40a (but arising from five different source glycoproteins), which associated with lymph node status and surgical margin involvement (*23*), Figure 7A. Surprisingly, three of the source *N*-glycoproteins (BTN3A1, SIGLEC1 and PLTP) were not identified in the global proteomics data, Supplementary Table S6. Moreover, while the *N*-glycopeptide from COL6A3 was less abundant in tumours from N+ patients, the source glycoprotein was found in higher levels in these individuals. Finally, ITGB6, whose glycopeptide was more abundant in N+ patients, displayed reduced protein abundance in this condition, observations that collectively indicate separate protein and glycan regulation in OSCC, Figure 7B.

**Fig. 7.**
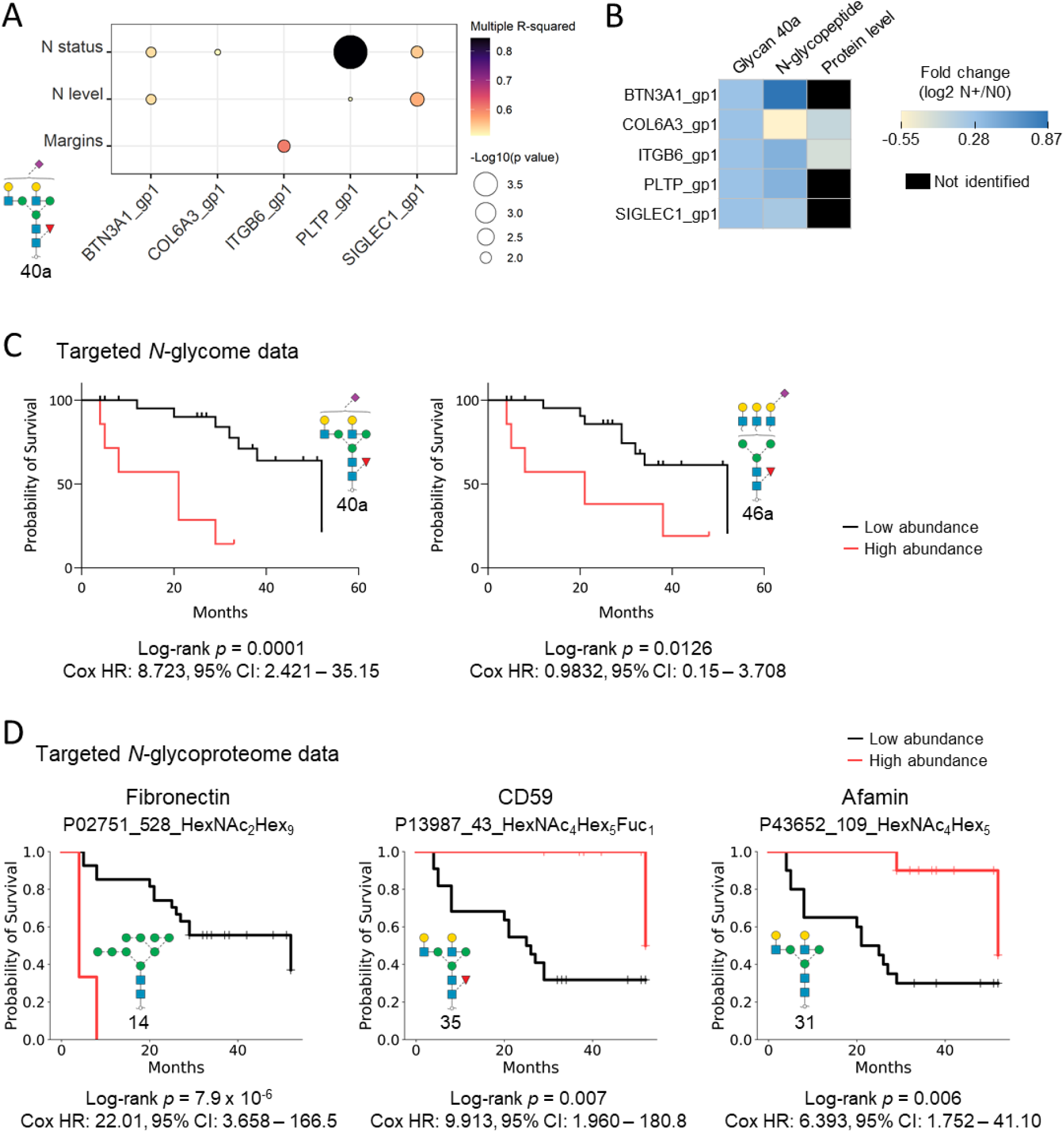
*N*-glycan and *N*-glycopeptide levels are associated with clinical outcomes in OSCC. A) Five out of 25 *N*-glycopeptides that associated with clinicopathological features were found to carry Glycan 40a. B) Distribution of fold change values (log2 N+/N0) of Glycan 40a, glycopeptides carrying Glycan 40a and their source glycoproteins in the OSCC patient cohort. C) Glycan-guided Kaplan-Meier survival analysis. Patients with relative high levels of Glycan 40a and Glycan 46b presented worse overall survival (*p* < 0.05, log-rank test). D) Glycopeptide-guided Kaplan-Meier survival analysis. Left: Relative high abundance of a HexNAc_2_Hex_9_ glycopeptide from fibronectin was associated with relatively low patient survival (elevated glycopeptide levels found in N+ samples). Middle-right: Relative low abundance of *N*-glycopeptides from CD59 and afamin, respectively, were associated with a relatively low patient survival (reduced levels in N+ samples).

Notably, Kaplan-Meier plots showed that relatively high levels of Glycan 40a and Glycan 46a were associated with a relatively poor patient survival, Figure 7C. The potential prognostic value of Glycan 40a and 46a was supported by their ability to stratify N0 and N+ patients as demonstrated by high ROC-AUC values (AUC 78.1 for Glycan 40a and AUC 85.5 for Glycan 46a, both by logistic regression model), Supplementary Figure S5.

Similarly, three *N*-glycopeptides that displayed abundance differences between N0 and N+ showed an association with patient survival, Figure 7D. Specifically, higher abundance of a fibronectin peptide (P02751) carrying an M9 oligomannosidic *N*-glycan (Glycan 14) was found to associate with lower patient survival, while reduced levels of a CD59 peptide (P13987) carrying a fucosylated complex *N*-glycan (Glycan 35) and an afamin peptide (P43652) carrying a related non-fucosylated *N*-glycan variant (Glycan 31) were associated with lower patient survival. These three *N*-glycopeptides also demonstrated an ability to stratify N0 and N+ patients as demonstrated by ROC analysis, Supplementary Figure S9.

Taken together, unbiased and global as well as targeted interrogations of our quantitative *N*-glycome and *N*-glycoproteome data pointed to previously unknown associations between OSCC tumour *N*-glycosylation and key clinical outcomes including, most importantly, lymph node metastasis and patient survival.

## Discussion

Previous studies have investigated the glycosylation patterns in oral cancer within diverse biological specimens including in cell lines (*12, 20*), tumour tissues (*24*), saliva (*13, 21, 25*), blood (*13*), plasma (*26*) and serum (*27, 28*) in attempts to uncover changes related to cell transformation (*12*) and with disease progression (*13*). Most of those studies evaluated the total levels of specific glycans or glycan features (*13, 21, 24, 26*) using relatively simple analytical techniques such as lectin-based strategies (*24, 26, 27*) and MALDI-MS profiling (*12, 20, 27, 28*) not able to survey the glycoproteome in an unbiased and quantitative manner with glycan fine structure and site-specific information (*29*). Our study is the first to use integrated glycomics and glycoproteomics to provide detailed insights into the heterogenous *N*-glycoproteome of OSCC tumour tissues obtained from a valuable cohort of patients with and without lymph node metastasis, a principal prognostic factor in OSCC (*3, 30, 31*).

To allow for a comprehensive examination of the global *N*-glycosylation of OSCC tumour tissues using our glycomics-informed glycoproteomics method (*22*), we firstly obtained quantitative *N*-glycomics data which served to establish a detailed *N*-glycome profile and inform a customised *N*-glycan database to aid the challenging downstream glycoproteomics data analysis. Intuitively, tailoring the glycan database reduces the search space resulting in lower glycopeptide FDRs and reduced search times (*32*) and ultimately more accurate quantitative information of the protein carriers, site occupancy (macroheterogeneity), and site distribution (microheterogeneity) of *N*-glycosylation, Figure 1 and Supplementary Figure S1.

We found surprisingly few global differences in the *N*-glycome between the N+ and N0 patient groups including in the *N*-glycan class distribution and *N*-glycosylation features (core fucosylation, sialylation and branching) suggesting a relative stable glycosylation machinery during lymph node metastasis, Figure 2B. Related studies have reported more dramatic glycan differences including altered sialylation, fucosylation and branching when comparing OSCC patients with healthy donors (*24, 28*) suggesting that disease onset rather than disease progression (involving lymph node metastasis) has a stronger influence on glycan remodelling.

Six sialic acid-capped *N*-glycans (displayed in Figure 6A) showed different expression between N+ and N0 tumour tissues. These six *N*-glycans appeared structurally (and thus biosynthetically) unrelated, suggesting that alteration in the abundance and/or activity of multiple glycosylation enzymes from a diversity of cellular/tissue origins may contribute to the subtle glycan remodelling observed upon lymph node metastasis. While their altered expression patterns remain mechanistically unexplained, the six differentially expressed *N*-glycans exhibited an interesting capacity to accurately stratify the N+ and N0 patient groups with confidence (AUC values > 60%), Supplementary Figure S5.

Bisecting GlcNAcylated *N*-glycans were consistently observed in the OSCC tumour tissue *N*-glycome (albeit at relatively low abundance) as confirmed by their early PGC-LC elution and prominent D- and D-GlcNAc fragment ions (*33*). Aberrant expression of bisecting *N*-glycans has repeatedly been reported as a feature of human cancers (*34*) including in colorectal cancer (*35*), prostate cancer (*22*), breast cancer (*36*) and, importantly, in oral cancer (*37*). Contrasting these reports, our *N*-glycome and *N*-glycoproteome data did not reveal any differences in bisecting GlcNAcylation across the N0 and N+ tumour tissues indicating that bisecting GlcNAc is not involved directly in the risk of lymph node metastasis in OSCC.

With a total of 3,117 identified *N*-glycopeptides from 419 source glycoproteins, the OSCC tumour tissue *N*-glycoproteome displayed an extreme complexity and dynamic range, Figure 3. Recapitulating the stable *N*-glycome, no striking global *N*-glycoproteome differences were observed between the N0 and N+ tumour tissues. Only 79 *N*-glycopeptides (∼2.5%) were found to be altered between the N0 and N+ patient groups and, similarly to the regulated *N*-glycans, these appeared biosynthetically unrelated spanning all *N*-glycan classes and belonging to different source proteins supporting that lymph node metastasis is a complex process that is not limited to the dysregulation of only a single glycosylation enzyme from a single cellular origin. Importantly, the 79 *N*-glycopeptides demonstrated a notable potential to accurately stratify N+ from N0 patients, Supplementary Table S7.

Seeking to mine the information-rich *N*-glycome and *N*-glycoproteome data further, unsupervised clustering revealed distinct tumor clusters that displayed interesting differences in the *N*-glycan class distribution including pauci- and oligomannosidic *N*-glycans, Figure 4A-B. While less explored relative to the more widely recognised cancer-associated changes in the complex/hybrid-type *N*-glycans and related features such as fucosylation and sialylation (*15, 38, 39*), we and others have recently shown that both pauci- and oligomannosylation are important *N*-glycan features altered in a wide range of human cancers (*40, 41*). The altered expression of these mannose-terminating *N*-glycans across the tumour clusters implies regulation, and possibly functional involvement, of the family of α-mannosidases (*40, 42*) and β-hexosaminidases (*43, 44*) in OSCC, speculations which require further exploration.

Through enrichment analysis, we investigated biological functions of the glycoproteins localising to each glycopeptide cluster, and observed extracellular matrix (ECM) organisation, and platelet and neutrophil degranulation being enriched processes, Supplementary Figure S7. ECM remodelling has repeatedly been linked to cancer development and progression, including changes in the abundance and composition of ECM components, post-translational modifications (e.g. glycosylation) (*38, 45*), and proteolytic activity, with several studies pointing to tumour-driven changes in the ECM supporting tumour growth, tumour cell migration and metastatic processes (*46, 47*). Further, platelet degranulation has been associated with cancer metastasis, possibly through the release of platelet factors that can enhance vascular permeability, and consequently promote tumour cell extravasation (*48, 49*). Finally, elevated neutrophil degranulation observed in a subset of the OSCC tumour tissues not only aligns with robust literature pointing to the involvement of neutrophils in tumour progression (*50*), but also provide a possible mechanistic explanation for the *N*-glycan differences observed between the tumour clusters including altered expression of paucimannosidic and highly truncated (chitobiose core) *N*-glycans. Neutrophils are namely known to store and upon activation secrete bioactive and highly unusual *N*-glycoproteins carrying paucimannosidic and chitobiose core type *N*-glycans as we have demonstrated (*51–54*). Infiltration of neutrophils in tumours (tumour-associated neutrophils) has been described in many cancers (*55*), including oral cancer, in which high neutrophil counts have been associated with poor clinical outcomes (*56*). In an early study, Wang *et al.* (*57*) identified that tongue squamous cell carcinomas exhibiting high neutrophil infiltration displayed increased lymph node metastasis, more advanced clinical stage and increased risk of tumour recurrence.

Interesting associations between key clinical features and two well-separated *N*-glycan- and *N*-glycopeptide-guided tumour clusters were identified through advanced clustering analysis. In addition to the association with the lymph node status, the *N*-glycan-guided tumour clusters informed by Glycan 20a, Glycan 34, Glycan 40a, Glycan 45b, Glycan 46a, and Glycan 49b were associated with vascular invasion, Figure 6A-B. Likewise, the two *N*-glycopeptide-informed tumour clusters contained patients that exhibited different tumour size and that received different treatments, Figure 6C-D. Moreover, correlation analysis of the 79 differentially expressed *N*-glycopeptides revealed associations with 1) the presence of perineural invasion, an important indicator of poor prognosis in oral cancer (*58*), 2) extracapsular extension, which is associated with poor prognosis and increased risk of recurrence and survival in head and neck cancers (*1, 59*), 3) the WPOI (worst pattern of invasion) score used for histological risk assessment in oral cancer (*60*), amongst other key clinical patient features, Supplementary Table S8 and Supplementary Figure S8.

Amongst the 25 *N*-glycopeptides associated with clinicopathological features, five were identified to carry Glycan 40a (a mono-α2,6-sialylated core fucosylated complex-type *N*-glycan), Figure 7A. One of the five Glycan 40a-carrying glycopeptides belongs to BTN3A1 (butyrophilin subfamily 3 member A1 or CD277) and was identified to associate with lymph node status. BTN3A1 plays a well-known role in T-cell activation and in the adaptive immune response, and altered BTN3A1 expression has been reported in different human cancers including in head and neck squamous cell carcinoma and in breast cancer (*61*). Remarkably, BTN3A1 was not detected in our OSCC proteome profile, Figure 7B, but only identified in HILIC-enriched glycoproteome analysis, highlighting the importance of employing multi-omics approaches to obtain a comprehensive coverage and holistic insights in the disease complexity and dynamics.

Interestingly, focused analysis of the *N*-glycome and *N*-glycoproteome data demonstrated an association between two *N*-glycans (Glycan 40a and 46a, both mono-α2,6-sialylated core fucosylated complex-type *N*-glycans) and three seemingly unrelated neutral *N*-glycopeptides and low patient survival rate, Figure 7C-D. Powered by our glycomics-assisted glycoproteomics method, the identification of several survival-associated glycans and glycopeptides expands on similar findings made in colorectal cancer (*62*) and gastric cancer (*63*), and represents an extremely exciting discovery as it opens for previously unexplored avenues to better prognosticate, monitor and manage patients diagnosed with OSCC.

In summary, this is the first study to report on the complexity and dynamics of the OSCC tumour tissue *N*-glycoproteome in patients with and without lymph node metastasis. Our comprehensive and quantitative *N*-glycomics and *N*-glycoproteomics data have unveiled a range of previously unknown associations between protein *N*-glycosylation in OSCC tumour tissues and key patient clinical features and survival outcome, and provide an important publicly-available resource to explore the underpinning disease mechanisms and to uncover potential prognostic glyco-markers for OSCC.

## Materials and Methods

### OSCC tumour tissues

The study was approved by the Ethics Review Board of the Cancer Institute of São Paulo (ICESP), Octavio Frias de Oliveira, ICESP, São Paulo, SP, Brazil, and Plataforma Brasil (protocol CAAE 61402116.8.0000.0065). Informed consent was obtained from all patients included in the study. The methods and experimental protocols were performed in accordance with the approved guidelines and regulations. Surgically removed oral squamous cell carcinoma (OSCC) tissues provided as formalin-fixed paraffin-embedded (FFPE) tissues from a 31-patient cohort collected over a two-year period (2017-18) were investigated. In total, 19 patients presented lymph node metastasis (N+) and 12 patients presented without lymph node metastasis (N0) as determined by histopathology. See Supplementary Table S1 for key clinicopathological data.

### Tissue sample preparation

Eight histological sections (each 10 μm thick) were prepared for each of the 31 OSCC FFPE tissues. Paraffin blocks were cut using a microtome and the tumour area was assigned by a trained pathologist for manual excision. A total tumour area of 7-8 cm^2^ were excised from 3-8 histological sections from each tumour tissue sample and transferred to a microtube for deparaffinization and protein extraction using a published protocol with modifications (*64*). For paraffin removal, 1 mL xylene was added to each sample, mixed for 1 min and centrifuged at 12,000 x g for 15 min at room temperature. The pellet was washed with 1 mL ethanol, mixed for 30 s, centrifuged as above and dried in a vacuum concentrator. Samples were resuspended in 300 µL lysis buffer containing 0.1 M *N*-(2-hydroxyethyl)piperazine-*N*′-(3-propanesulfonic acid) (EPPS), 0.1 M dithiothreitol (DTT), 1 x protease inhibitor cocktail (cOmplete ULTRA Tablets, Mini, EDTA-free, Roche; 1:10), pH 8.5. SDS was added to a final concentration of 4% (w/v) and the samples incubated for 60 min at 99°C with agitation (400 rpm). Samples were then centrifuged at 15,000 x g for 30 min at 4°C and submitted to ice-cold acetone precipitation with three-fold (v/v) excess acetone for 16 h at −20°C. Following centrifugation at 15,000 x g for 30 min at 4°C, protein pellets were resuspended in 30 µL buffer containing 8 M urea, 50 mM EPPS, protease inhibitor cocktail, 1 mM sodium fluoride, 1 mM sodium orthovanadate, 1 mM phenylmethylsulfonyl fluoride (PMSF), 1 mM EDTA, pH 8.5. Protein concentrations were determined by a bicinchoninic acid (BCA) protein assay kit (Thermo Fisher Scientific, Waltham, MA) and stored at −20°C until further handling.

### *N*-glycan release

Glycans were prepared as described (*65*). Briefly, the protein extracts (15 μg/sample) were blotted on a primed 0.45 μm polyvinylidene fluoride (PVDF) membrane (Merck-Millipore). Protein spots were stained using Direct Blue (Sigma-Aldrich, Australia), excised and transferred to a flat bottom polypropylene 96-well plate (Corning Life Sciences, Australia), blocked with 1% (w/v) polyvinylpyrrolidone in 50% (v/v) methanol and washed with MilliQ water. The *N*-glycans were then released using 10 U recombinant *Elizabethkingia miricola N*-glycosidase F (Promega, V4831, 10 U/μL) for 16 h at 37°C. The detached *N*-glycans were reduced with 1 M sodium borohydride in 50 mM aqueous potassium hydroxide for 3 h at 50°C. The reaction was stopped using glacial acetic acid and the *N*-glycans desalted sequentially using strong cation exchange (AG 50W X8, Bio-Rad), C18 and porous graphitised carbon (PGC, Thermo Fisher Scientific) solid phase extraction. The desalted *N*-glycan samples were stored at −20°C until analysis.

### *N*-glycome profiling

*N*-glycomics data were acquired using an established PGC-LC-MS/MS method (*65, 66*), see Supplementary Table S2 for experimental overview and details of all generated ‘omics datasets. The *N*-glycans were separated on an UltiMate 3000 HPLC system (Dionex, Sunnyvale, CA, USA) interfaced with a linear ion trap quadrupole (LTQ) Velos Pro (Thermo Scientific, San Jose, CA, USA). The glycan samples (3 µL injection volume) were loaded on a PGC HPLC capillary column (Hypercarb KAPPA, 5 μm particle size, 200 Å pore size, 180 μm inner diameter x 100 mm length, Thermo Scientific) operated at 50°C with a constant flow rate (4 μL/min) supplemented with a post-column make-up flow supplying pure acetonitrile (ACN) delivered by the HPLC system. Aqueous ammonium bicarbonate (10 mM), pH 8.0 (solvent A) and 10 mM ammonium bicarbonate in 70% ACN (solvent B) were used as mobile phases with the following 86 min-gradient: 8 min at 2.6% B, 2.6-13.5% B over 2 min, 13.5-37.3% B over 55 min, 37-64% B over 10 min, 64-98% B over 1 min, 5 min at 98% B, 98-2.6% B over 1 min and 4 min at 2.6% B. The ESI source was operated in negative ion polarity mode with source potential of 3.6 kV. Full MS1 scans were acquired in the range *m/z* 570-2,000 using 1 microscan, *m/z* 0.25 full width half maximum (FWHM) resolution, 5 × 10^4^ automatic gain control (AGC) and 50 ms maximum accumulation time. MS/MS data were acquired using *m/z* 0.35 FWHM resolution; 2 × 10^4^ AGC, 300 ms maximum accumulation time, and 2 *m/z* precursor ion isolation window. The five most abundant precursors in each MS1 full scan were selected for collision-induced dissociation (CID)-based MS/MS using a normalised collision energy (NCE) of 33% with an activation Q of 0.250 and 10 ms activation time. To prevent instrument bias, the injection order of the glycan samples from the OSCC patient cohort was randomised (*67*), using the *psych* package (*68*) under R (v3.6.0).

### *N*-glycan data analysis

Xcalibur v2.2 (Thermo Scientific) was used to browse and interrogate the raw LC-MS/MS data. Putative glycan precursor ions were extracted using RawMeat v2.1 (Vast Scientific) (*22, 69*). Common contaminants not matching known *N*-glycan compositions and redundant precursors were manually removed. The monoisotopic precursors were searched against GlycoMod (*70*) with a mass tolerance of 0.5 Da to identify putative monosaccharide compositions. Only compositions containing Hex, HexNAc, dHex, NeuAc, and NeuGc were considered and *N*-glycans already reported in the UniCarbKB database were prioritised. The *N*-glycan fine structures were manually elucidated using monoisotopic mass, absolute and relative PGC-LC retention time and MS/MS fragmentation patterns as described (*33, 71*). EIC-based relative quantification of the identified *N*-glycans was performed using Skyline v.20.1.0.31 as described (*22, 72*). GlycoWorkBench v2.1 (*73*) was used to aid the manual annotation of the glycan fragment spectra and to generate glycan cartoons. Poorly expressed *N*-glycans with a relative abundance below 0.1% were excluded from the relative quantitation due to poor spectral signal-to-noise ratios.

### Protein digestion and peptide desalting

Protein samples (35 µg/sample) were reduced with 5 mM DTT (final concentration) for 25 min at 56°C and alkylated using 14 mM iodoacetamide (IAA) for 30 min at room temperature in the dark. Urea was added to a final concentration of 1.6 M in 50 mM ammonium bicarbonate and 1 mM calcium chloride prior to digestion with 1 µg sequencing grade porcine trypsin (Promega) for 16 h at 37°C (*74*). The reaction was quenched with a final concentration of 0.4% (v/v) trifluoroacetic acid (TFA). The resulting peptides were desalted using Oasis HLB 1 cc (30 mg) SPE cartridges (Waters) using an established protocol with modifications (*74*). Columns were activated with 1 mL methanol, washed twice with 1.5 mL 1% formic acid (FA) in 70% ACN (both v/v) and equilibrated with 1.5 mL 1% (v/v) aqueous FA prior to sample loading. After washing twice with 1.5 mL 1% (v/v) aqueous FA, peptides were eluted twice with 250 µL 1% FA in 70 % ACN (both v/v) and dried.

### TMT labelling of peptides

A reference sample was prepared to allow for quantitative comparisons across multiple TMT experiments by pooling 3 µg digested protein from each sample. In total, 25 µg from this pool and 25 µg protein extract from each sample were used for TMT labelling (*22*). The peptide samples were randomised before being labelled using three separate TMT-11plex sets. The reference sample was consistently labelled with the 131 Da (131C) reporter ion channel in all three sets. In total, 30 peptide samples were labelled with TMT for quantitative proteomics including 12 N0 and 18 N+ samples (one random N+ sample from the sample cohort was excluded in TMT experimental design due to limited channels available), Supplementary Table S2. For the TMT labelling, 100 μL 100 mM triethylammonium bicarbonate buffer (final concentration) was added to each peptide sample that was labelled individually with 0.4 mg TMT-11plex mass tags (Thermo) in 41 μL anhydrous ACN over a 1 h incubation period at room temperature. The labelling reactions were quenched using 8 μL 5% (v/v) hydroxylamine for 15 min at room temperature. After TMT labelling, all peptide samples were mixed 1:1:1 (w/w/w), desalted using HLB SPE cartridges (Waters) and dried. Aliquots of the TMT-labelled peptide mixtures from the three TMT-11plex experiments were analysed without any pre-fractionation by LC-MS/MS, Supplementary Table S2.

### Glycopeptide enrichment

TMT-labelled peptides (from 275 μg protein extract from 10 tumour tissue samples + one pool) was reconstituted in 50 μL loading/washing solvent containing 1% TFA in 80% ACN (both v/v). Five microliters (∼18 µg protein digest) were allocated to the proteome analysis. The remaining peptides were loaded onto primed custom-made HILIC SPE micro-columns packed with ZIC-HILIC resin (10 μm particle size, 200 Å pore size, kindly provided by Sequant/Merck, Umea, Sweden) onto supporting C8 disks (Empore) in p10 pipette tips (*75*). The flow-through fractions were collected. The HILIC-SPE micro-columns were then washed with 50 μL loading/washing solvent and the wash fraction combined with the flow-through fraction for separate downstream analysis, as this fraction contained the non-glycosylated peptides. The retained *N*-glycopeptides were eluted in three sequential steps, firstly with 50 μL 0.1% (v/v) aqueous TFA, followed by 50 μL 25 mM aqueous ammonium bicarbonate and then 50 μL 50% (v/v) ACN. The three *N*-glycopeptide fractions were combined to form the enriched glycopeptide mixture, which was dried and desalted on a primed Oligo R3 reversed phase SPE micro-column. R3 resin suspended in ACN was packed into a p10 pipette tips, washed three times with 50 μL ACN, and then washed with 0.1% (v/v) aqueous TFA before sample loading. The sample loading was repeated once to ensure high recovery ahead of three washing steps with 0.1% (v/v) aqueous TFA and elution with 50 μL 0.1% TFA in 50% ACN (both v/v) and then with 50 μL 0.1% TFA in 70% ACN (both v/v). The desalted glycopeptide fractions were combined, dried and stored at −20°C (*22*).

### High-pH reversed phase prefractionation

The HILIC enriched *N*-glycopeptides and HILIC flow-through fractions containing the non-glycosylated peptides were resuspended separately in 50 μL 25 mM aqueous ammonium bicarbonate for high pH pre-fractionation using Oligo R2 reversed phase SPE micro-columns packed on supporting C18 discs (Empore) in standard p10 pipette tips. The SPE micro-columns were primed three times with 50 μL ACN and then washed with 50 μL 25 mM aqueous ammonium bicarbonate. Samples were loaded on the columns followed by two washing steps with 50 μL 25 mM aqueous ammonium bicarbonate. The peptides were eluted in three fractions i.e. fraction 1: 25 mM ammonium bicarbonate in 10% (v/v) ACN; fraction 2: 25 mM ammonium bicarbonate in 20% (v/v) ACN, and fraction 3: 25 mM ammonium bicarbonate in 60% (v/v) ACN. Each eluted fraction was dried and resuspended in 0.1% (v/v) aqueous FA for separate LC-MS/MS analysis.

### (Glyco)proteome profiling by LC-MS/MS

Approximately 1 μg (glyco)peptide material was injected per LC-MS/MS run. The (glyco)peptides were loaded on a trap column (2 cm length x 100 μm inner diameter) custom packed with ReproSil-Pur C18 AQ 5 μm resin (Dr. Maisch, Ammerbuch-Entringen, Germany) and separated at a constant flow rate of 250 nL/min on an analytical column (Reprosil-Pur C18-Aq, 25 cm length x 75 μm inner diameter, 3 μm particle size, Dr. Maisch, Ammerbuch-Entringen, Germany) using an UltiMate™ 3000 RSLCnano System. The mobile phases were 0.1% FA in 99.9% (both v/v) ACN (solvent B) and 0.1% (v/v) aqueous FA (solvent A). The gradient was 2-30% B over 100 min, 30-50% B over 18 min, 50-95% B over 1 min and 9 min at 95% B. The nanoLC was connected to a Q-Exactive HF-X Hybrid Quadrupole-Orbitrap mass spectrometer (Thermo Fisher Scientific) operating in positive ion polarity mode. The Orbitrap acquired full MS1 scans with an AGC of 3 x 10^6^ ions and 50 ms maximum accumulation time. Full MS1 scans were acquired at high-resolution (60,000 FWHM at *m/z* 200) in the *m/z* 350-1,800 range. The 20 most abundant precursor ions were selected from each MS1 full scan using data-dependent acquisition and were fragmented utilising higher energy collision-induced dissociation (HCD) with a NCE of 35%. Only multicharged precursors (z ≥ 2) were selected for fragmentation. Fragment spectra were acquired at 45,000 resolution with an AGC of 1 x 10^5^ and 90 ms maximum accumulation time using a precursor isolation window of *m/z* 1.0 and a dynamic exclusion of 30 s after a single isolation and fragmentation of a given precursor ion.

### *N*-glycoproteomics data analysis

The HCD-MS/MS data of intact *N*-glycopeptides were searched with Byonic v2.6.46 (Protein Metrics Inc, CA, USA) (*76*) using 10/20 ppm as the precursor/product ion mass tolerance, respectively. Cys carbamidomethylation (+57.021 Da) and TMT (+229.163 Da) at N-term and lysine (K) were considered fixed modifications. Trypsin specific cleavages were considered with a maximum of two missed cleavages allowed per peptide. The following variable modifications were considered: Met oxidation (+15.994 Da), and *N*-glycosylation of sequon-localised Asn with a glycomics-informed *N*-glycan database comprising the *N*-glycan compositions identified in the OSCC tissue *N*-glycome data (Supplementary Table S3) and HexNAc_1_, HexNAc_1_Fuc_1_ and HexNAc_2_ that were manually added to the database. A maximum of two common modifications and a maximum of one rare modification were allowed. The HCD-MS/MS data were searched against a protein database composed of all reviewed UniProtKB human proteins (20,300 sequences, released December 11, 2019). All searches were filtered to <1% false discovery rate (FDR) at the protein level and 0% at the peptide level by using a protein decoy database (*77*). Only *N*-glycopeptides confidently identified with PEP 2D scores < 0.001 were considered (*22*). Glycopeptides identified with low confidence, and those in the reverse database and contaminant database were excluded. The identified *N*-glycopeptides were quantified using the ‘Report Ion Quantifier’ available as a node in Proteome Discoverer v2.2 (Thermo Scientific) and the reporter ion intensities from the MS/MS scans were extracted from the QuantSpectra table (*22*). Glycopeptides were manually grouped by summing the reporter ion intensities from the glycopeptide spectral matches (glycoPSMs) belonging to the same UniProtKB identifier, same glycosylation site within the protein, and same glycan composition. The abundances of the unique glycopeptides from each channel were first normalised by dividing each unique glycopeptide reporter ion intensity by the reference reporter intensity (from the peptide pool, see above) within that specific experiment and further normalised by the total sum intensity of each channel to correct for any inter-sample variation in the total yield during the labelling reactions.

### Proteomics data analysis

LC-MS/MS-based proteomics data were acquired of both unenriched peptide samples not subjected to glycopeptide enrichment as well as to the HILIC flow-through fractions. All data were processed using MaxQuant v1.6.12.0 (*78*). The HCD-MS/MS data were searched against the reviewed UniProtKB Human Protein Database (20,300 sequences, released December 11, 2019) using the Andromeda search engine (*79*) with a tolerance of 4.5 ppm for precursor ions and 20 ppm for product ions. Report fragment ions “10plex TMT” were enabled in the quantification settings and the enzyme specificity was set to trypsin with a maximum of two missed cleavages permitted. Carbamidomethylation of Cys (+57.021 Da) was considered a fixed modification, and oxidation of Met (+15.994 Da) and protein N-terminal acetylation (+42.010 Da) were considered variable modifications. Both the protein and peptide identifications were filtered to 1% FDR. Processing and statistical analyses (Student’s t-test) of the resulting data table (MaxQuant output) were performed in Perseus v1.6.14.0 (*78*). Proteins identified using the reverse database, proteins only identified through modified peptides and proteins identified from the MaxQuant contaminant database were excluded (except human keratins, which were not excluded since they are of interest in the study of squamous tissues (*80*)). The identified proteins were quantified by their reporter ion intensities using at least one razor/unique peptide per protein. Protein abundances of each channel were first normalised by dividing the reporter ion intensity by the reference protein reporter ion intensity (131C channel of peptide pool) within that specific LC-MS/MS run and further normalised by dividing by the sum of total intensity of each channel to correct for any inter-sample variation in the total yield during the labelling reactions.

### Statistical analysis

For the glycome profiling and glycosylation site analyses, statistical significance was assessed using unpaired two-tailed Student’s t-tests in which *p* < 0.05 was used as the confidence threshold. For the glycoproteome profiling, the statistical significance of the summed intensities of the glycopeptides grouped based on their *N*-glycan classes or tissue cluster was assessed using unpaired two-tailed Student’s t-tests in which *p* < 0.05 was used as the confidence threshold. For data visualisation, heat maps with z-score values of normalised intensities were generated using the open-source statistical programming language R.

### Clinical association and survival analysis

Linear regression analysis was performed using the R code to evaluate the linear relationship between glycan or glycopeptide abundance and the following clinicopathological variables: age (>63 or ≤63, mean age of patient cohorts chosen as the threshold), sex, smoking habit, alcohol consumption, tumour size, lymph node metastasis (N+, N0), clinical stage, type of treatment (surgery, surgery and radiotherapy, or a combination of surgery, radiation and chemotherapy), disease-free survival, presence of the worst pattern of invasion (*60*), presence of inflammatory infiltrate, and perineural invasion. Linear regressions with *p* < 0.05 were considered significant. The Pearson product−moment correlation coefficient (R) was also calculated to measure the strength of the association between variables. Only associations with R < −0.5 or 0.5 < R with at least six valid values per group and at least three valid values per clinical feature were considered (*81*). For cluster analysis and associations to clinical features, Fisher’s Exact Test (for two group comparisons) or Pearson Chi-Square test (for comparisons of more than two groups) were performed using IBM SPSS Statistics v 28.0.0.0 (190). Furthermore, a survival analysis was calculated on GraphPad Prism v.9.1.2 using the Kaplan-Meier methodology and compared with the log-rank test, evaluating the survival probability comparing a higher and lower abundance of specific glycoproteins, glycopeptides and glycans.

### ROC curve analysis by logistic regression and random forest models

The potential of select *N*-glycans and *N*-glycopeptides to stratify N0 and N+ patients was evaluated by the area-under-the-curve (AUC) of the constructed receiver operating characteristic (ROC) curves generated using random forest and logistic regression models. The area-under-the-curve (AUC-ROC) with a 95% confidence interval was used for comparison. Optimal cut-off by highest sensitivity (true positive rate) as a function of the specificity (false positive rate) was calculated, and 60% was chosen as the decision threshold. For all statistical comparisons, an ANOVA *p* < 0.05 was considered as significance threshold. The data analysis was performed using the package pROC (*82*) and the R environment version 3.6.0.

### Enrichment analysis of gene ontology (GO) terms

GO enrichment analysis was performed using the DAVID (*83*) or Enrichr (*84*) with corrected *p* < 0.05 chosen as the significance threshold. The entire human proteome GO annotation file was used as a reference set.

## ABBREVIATIONS

ACN: acetonitrile
DTT: dithiothreitol
EIC: extracted ion chromatogram
EPPS: *N*-(2- hydroxyethyl)piperazine-*N*′-(3-propanesulfonic acid)
FA: formic acid
FDR: false discovery rate
FFPE: formalin-fixed paraffin-embedded
FWHM: full width half maximum
HCD: higher energy collision-induced dissociation
HLB: hydrophilic − lipophilic-balanced
IAA: iodoacetamide
LC: liquid chromatography
LTQ: linear trap quadrupole
MS: mass spectrometry
N0: absence of lymph node metastasis
N+: presence of lymph node metastasis
NCE: normalised collision energy
OSCC: oral squamous cell carcinoma
PGC: porous graphitised carbon
PMSF: phenylmethylsulfonyl fluoride
PSM: peptide-to-spectrum match
PVDF: polyvinylidene fluoride
SPE: solid phase extraction
TFA: trifluoroacetic acid
TIC: total ion chromatogram
ZIC-HILIC: zwitterionic hydrophilic interaction liquid chromatography

## Acknowledgments

We thank Dr. Sami Yokoo for the assistance with tissue recovery and Dr. Daniela Granato for assistance with sample digestion. We also thank Dr Edward S.X. Moh and Dr Krishnatej Nishtala for assistance with the acquisition of glycomics data.

## Funding

Fundação de Amparo à Pesquisa do Estado de São Paulo (FAPESP) grant 2018/02180-0 (CMC)

Fundação de Amparo à Pesquisa do Estado de São Paulo (FAPESP) grant 2019/17840-8 (CMC)

Fundação de Amparo à Pesquisa do Estado de São Paulo (FAPESP) grant 2018/18496-6 (AFPL)

Cancer Institute NSW Grant ECF181259 (RK)

Australian Research Council (ARC) Future Fellowship grant (FT210100455) (MTA)

Macquarie University Enterprise Partnership Scheme (175232162) (MTA)

## Author contributions

Conceptualisation: CMC, AFPL, RK, MTA

Collected the clinical samples: ACPR, TBB, ES, LLM

Prepared tissues slides and assigned tumour area: TMLM

Methodology: CMC, AFPL, RK, MTA

Investigation: CMC, RK

Visualisation: CMC, FP, RK

Supervision: AFPL, RK, MTA

Writing-original draft: CMC

Writing-review & editing: AFPL, RK, LPK, MTA

## Data and materials availability

All proteomics and glycoproteomics LC-MS/MS raw data files supporting the conclusions presented herein have been deposited to the ProteomeXchange Consortium via the PRIDE (*85*) partner repository with the dataset identifier PXD037134. All glycomics LC-MS/MS raw data files are available via the GlycoPOST (*86*) with the identifier GPST000296.

## Supplementary Materials

Please see the attached Supplementary Materials.

## Supplementary Materials for

**Supplementary Figure S1.**
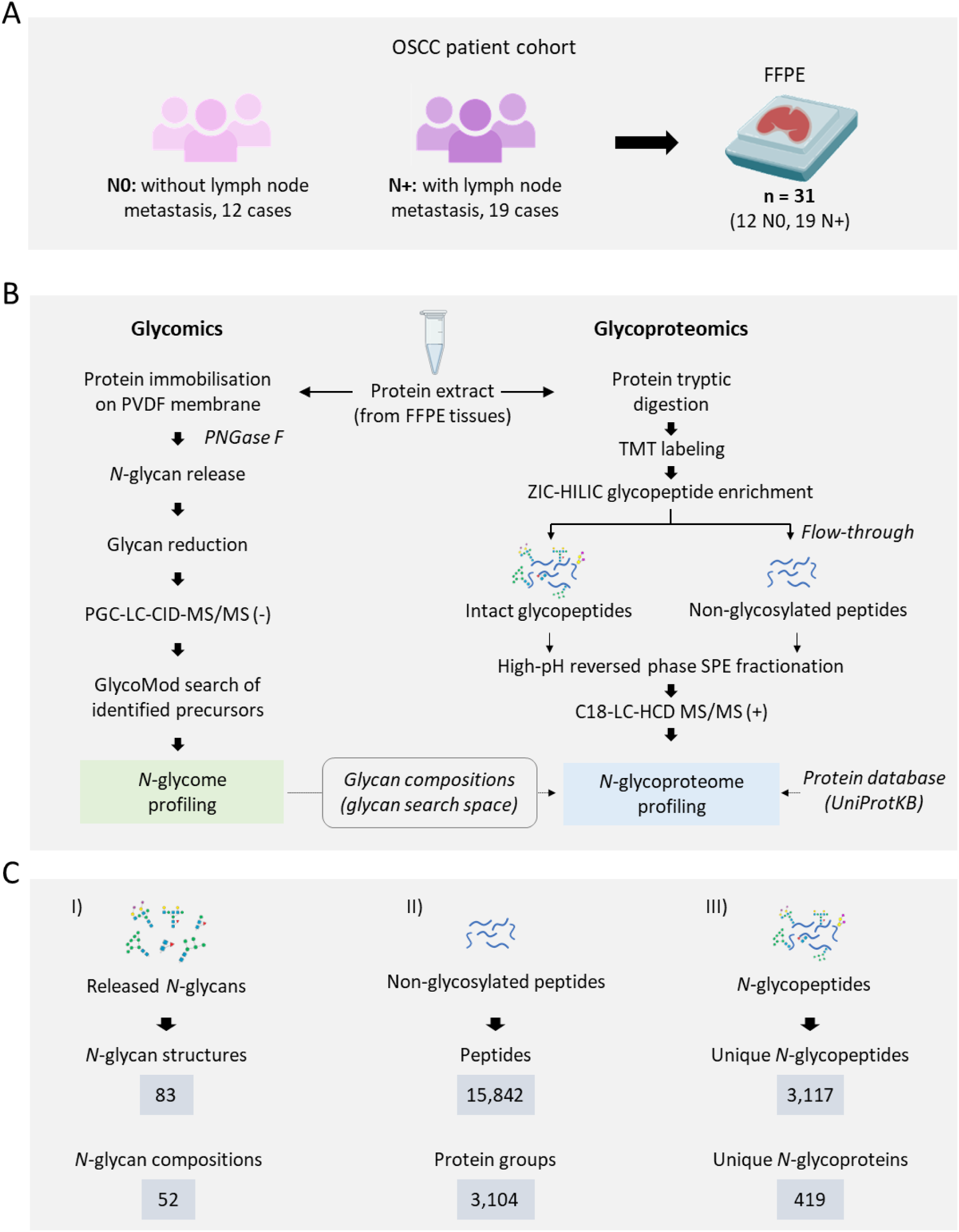
Study design and overview of the integrated glycomics and glycoproteomics approach. A) Overview of the sample cohort comprising resected tumour tissues (FFPE slides) from N0 and N+ OSCC patients. B) Overview of the integrated glycomics and glycoproteomics workflows. C) Total number of identifications for I) the *N*-glycome profiling including the *N*-glycan structures (including isomeric variants) and compositions, II) proteome profiling including the unique peptides and corresponding source proteins identified in the non-enriched (ZIC-HILIC flowthrough) fraction, III) *N*-glycoproteome profiling including the unique intact *N*-glycopeptides and corresponding source *N*-glycoproteins identified after ZIC-HILIC enrichment.

**Supplementary Figure S2.**
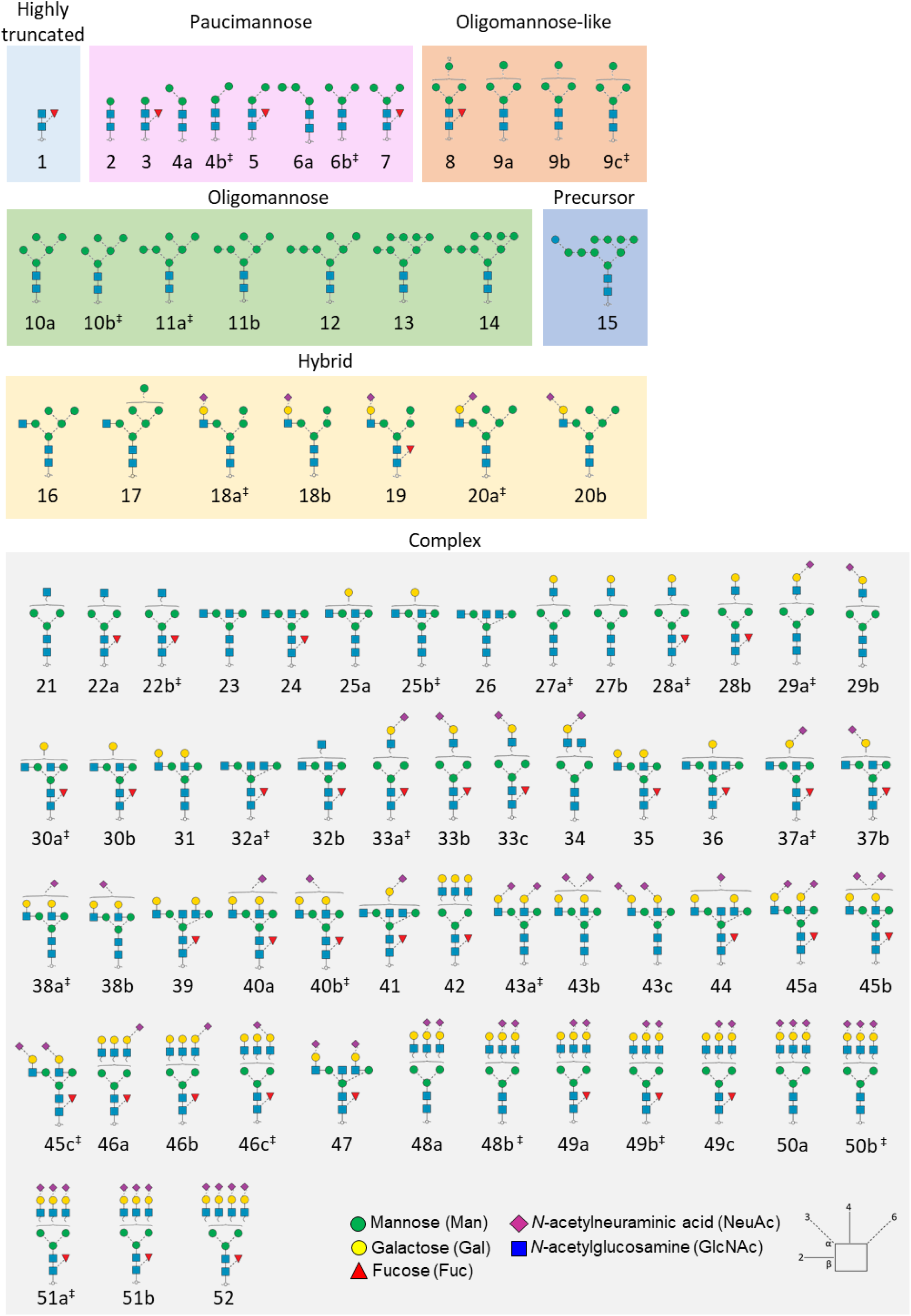
*N*-glycome map of the investigated OSCC tissues. Map of the confidently identified *N*-glycan isomers and the assigned glycan identifiers used consistently in this study (Glycan 1-52). The *N*-glycan isomers observed for each composition are denoted with lower case letters (a, b, c…). ^‡^The most abundant *N*-glycan isomer.

**Supplementary Figure S3.**
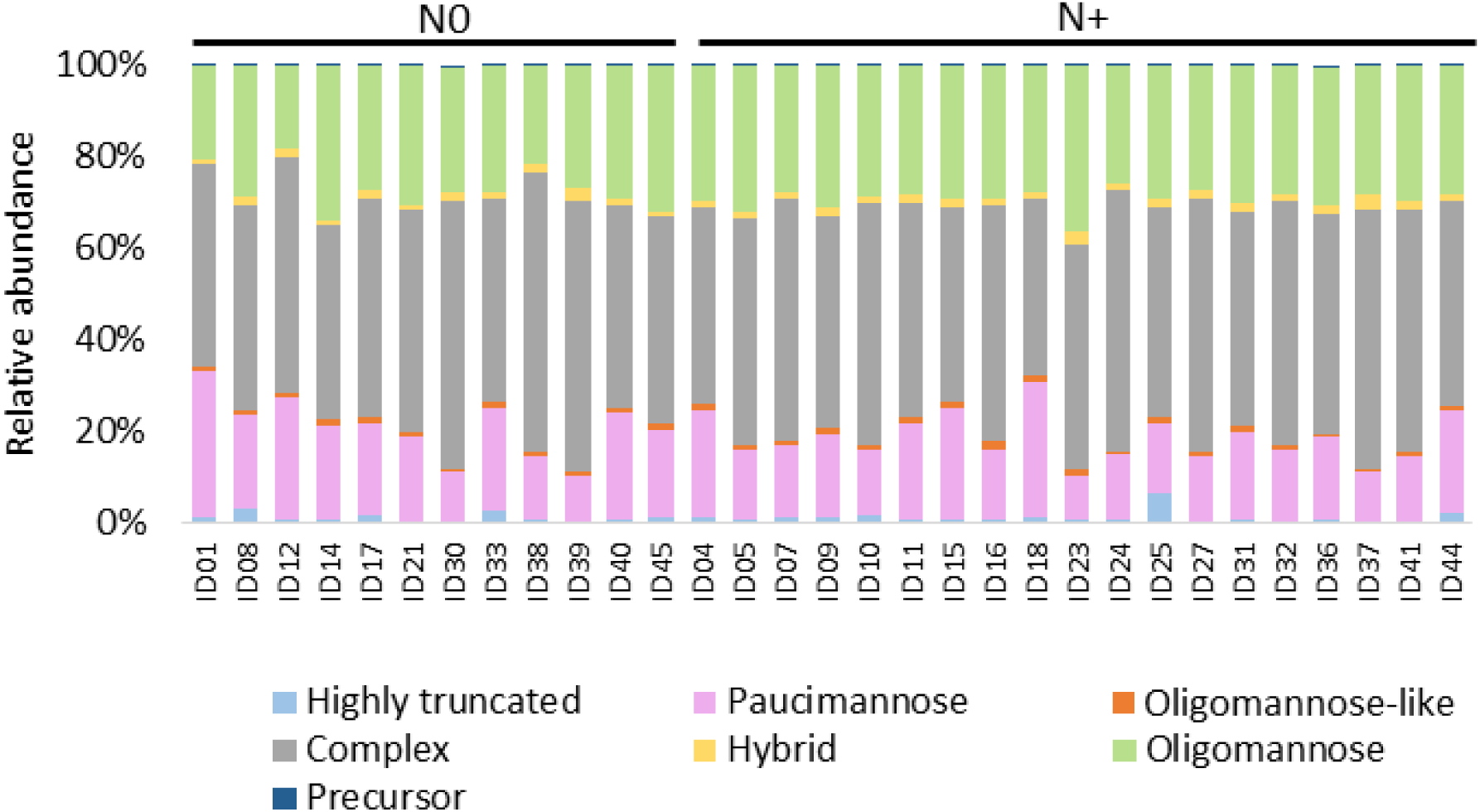
*N*-glycan type distribution across the studied OSCC tumour tissue samples. Relative abundance of the identified *N*-glycan types within each N0 and N+ sample classified into highly truncated, paucimannose, complex, hybrid, oligomannose, oligomannose-like and precursor-type *N*-glycans, see Supplementary Figure S2 for *N*-glycan structures and classification.

**Supplementary Figure S4.**
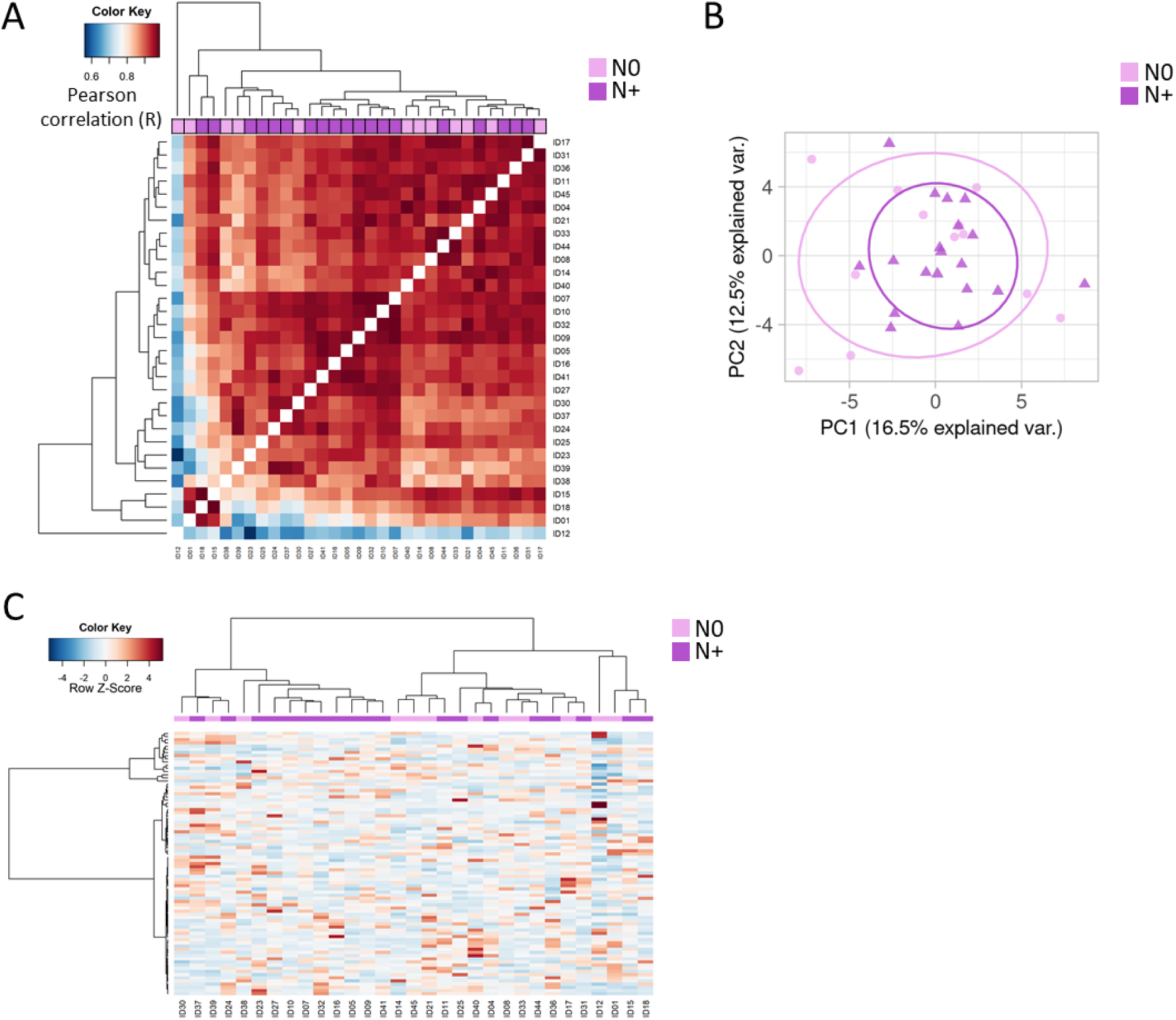
Uniform global *N*-glycosylation across the N0 and N+ OSCC tumour tissues. A) Heat map profile of Pearson correlation coefficients (R) derived from pairwise comparison of the quantitative *N*-glycome datasets comprising 83 *N*-glycan structures collected from the investigated OSCC tissue cohort. Relative *N*-glycan abundance values were used to calculate the correlation coefficient using the Perseus software, and the heat map was constructed using the R language with the function ‘heatmap.3’. The dendrogram was created using Euclidean distance with complete linkage. B) Principal component analysis of the *N*-glycome data, showing no distinct separation of the N0 and N+ patient groups. C) Unsupervised hierarchical clustering analysis of the *N*-glycome from the 12 N0 and 19 N+ samples performed with heatmap function under the R environment using Euclidean distance and Ward linkage.

**Supplementary Figure S5.**
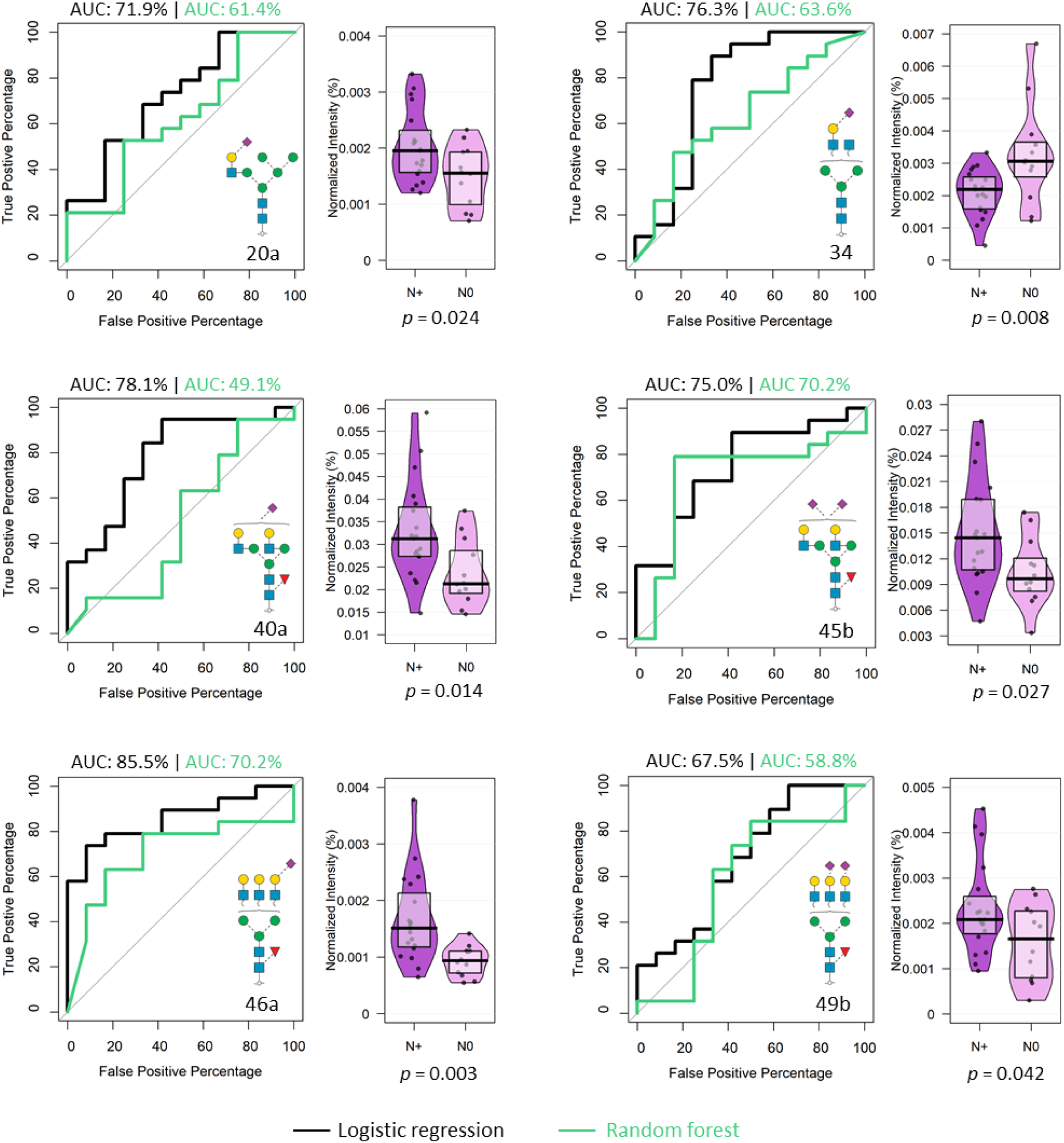
Tumour tissue *N*-glycans show potential for OSCC patient stratification. Five *N*-glycans (Glycan 20a, 40a, 45b, 46a and 49b) displaying increased expression in N+ tissues, and one *N*-glycan (Glycan 34) displaying decreased level in N+ tissues (relative to N0, see pirate plots in purple) show a potential to stratify the N0 and N+ patient groups with an AUC-ROC > 60% using the logistic regression model (black trace). Four of those *N*-glycans were also able to stratify the N0 and N+ patient groups by using a random forest model (AUC-ROC > 60%, green trace). AUC-ROC: area-under-the-curve of the receiver operating characteristic. *p* < 0.05 was used as a threshold to denote statistical significance in the applied Student’s t-tests.

**Supplementary Figure S6.**
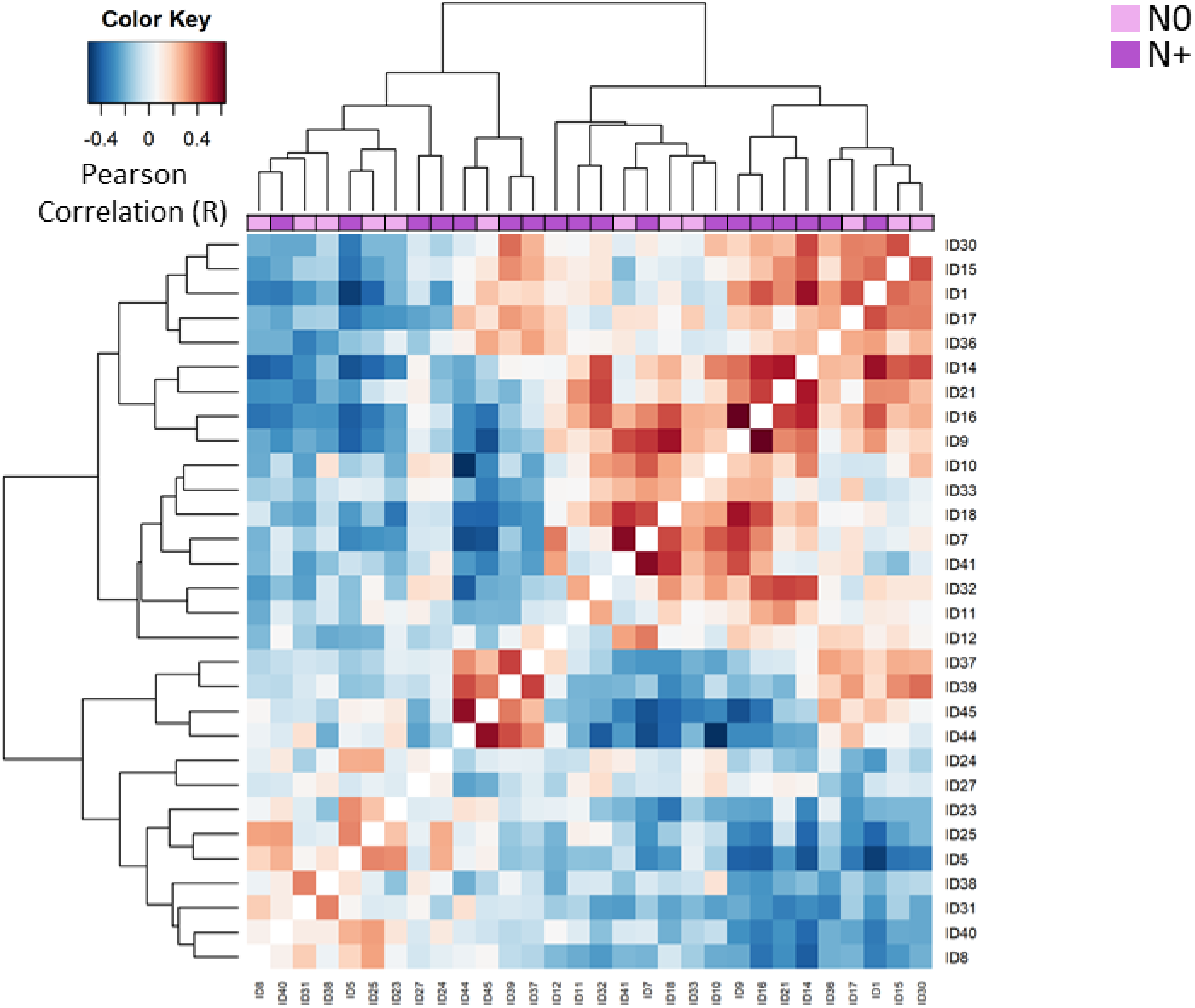
Correlation analysis of the *N*-glycoproteome data from N+ and N0 tumour tissues. Heat map of Pearson correlation coefficients (R) derived from pairwise comparisons of the tissue *N*-glycoproteome data collected across the N+ and N0 OSCC tumour tissues. Normalised intensity values of glycopeptides were used to calculate the correlation coefficient using the Perseus software, and the heat map was constructed using the R language with the function ‘heatmap.3’. The dendrogram was generated using Euclidean distance with complete linkage.

**Supplementary Figure S7.**
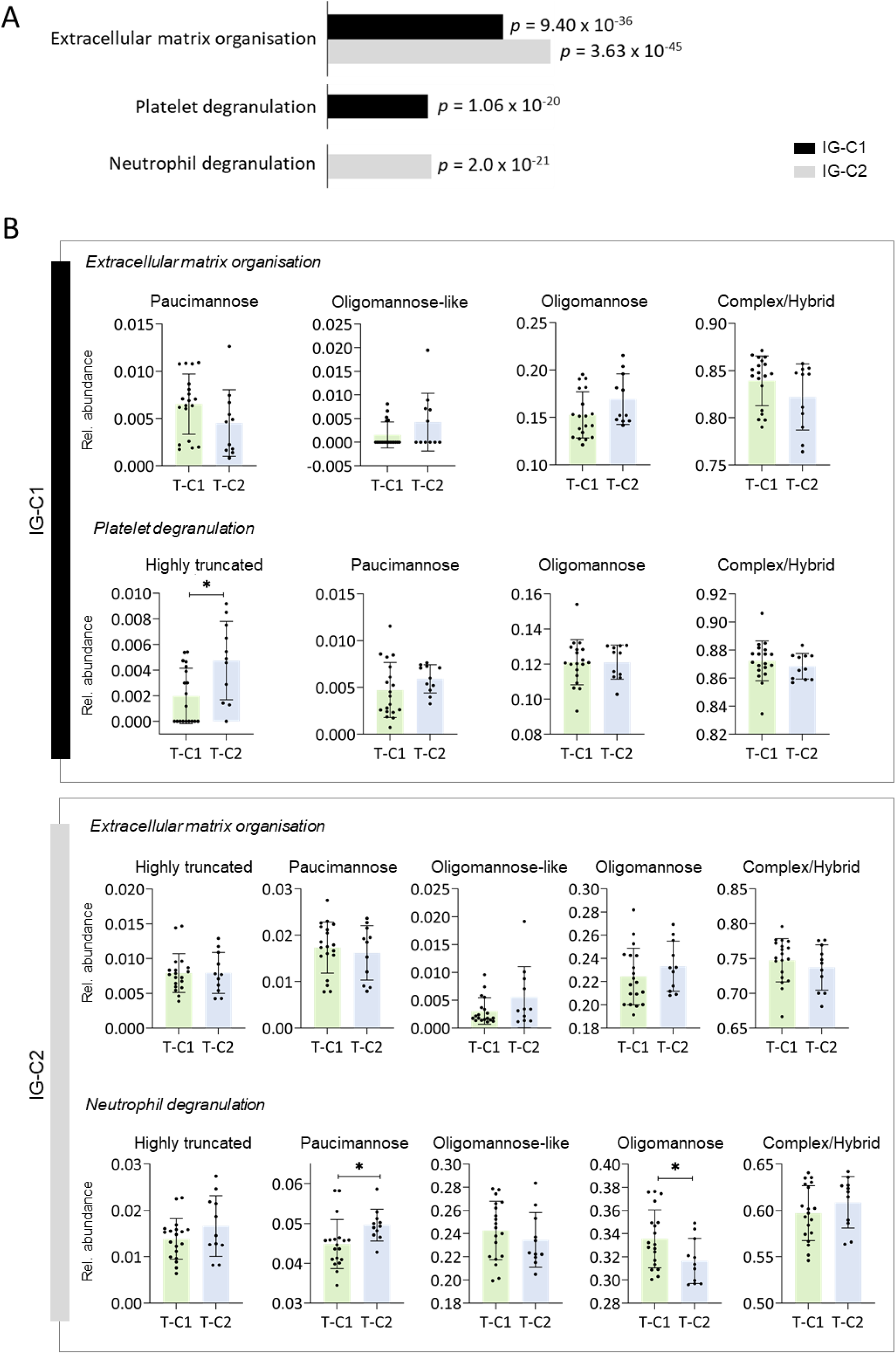
Enriched biological processes and altered glycan class distribution in the *N*-glycopeptide clusters. A) The most enriched biological processes in both *N*-glycopeptide clusters (IG-C1 and IG-C2). B) *N*-glycan class distribution of *N*-glycoproteins involved in the two most enriched biological processes in IG-C1 and IG-C2 within the T-C1 and T-C2 tumour clusters (Figure 5). Student’s t test, two-sided, α = 0.05, \**p* < 0.05, \*\**p* < 0.01, \*\*\**p* < 0.001.

**Supplementary Figure S8.**
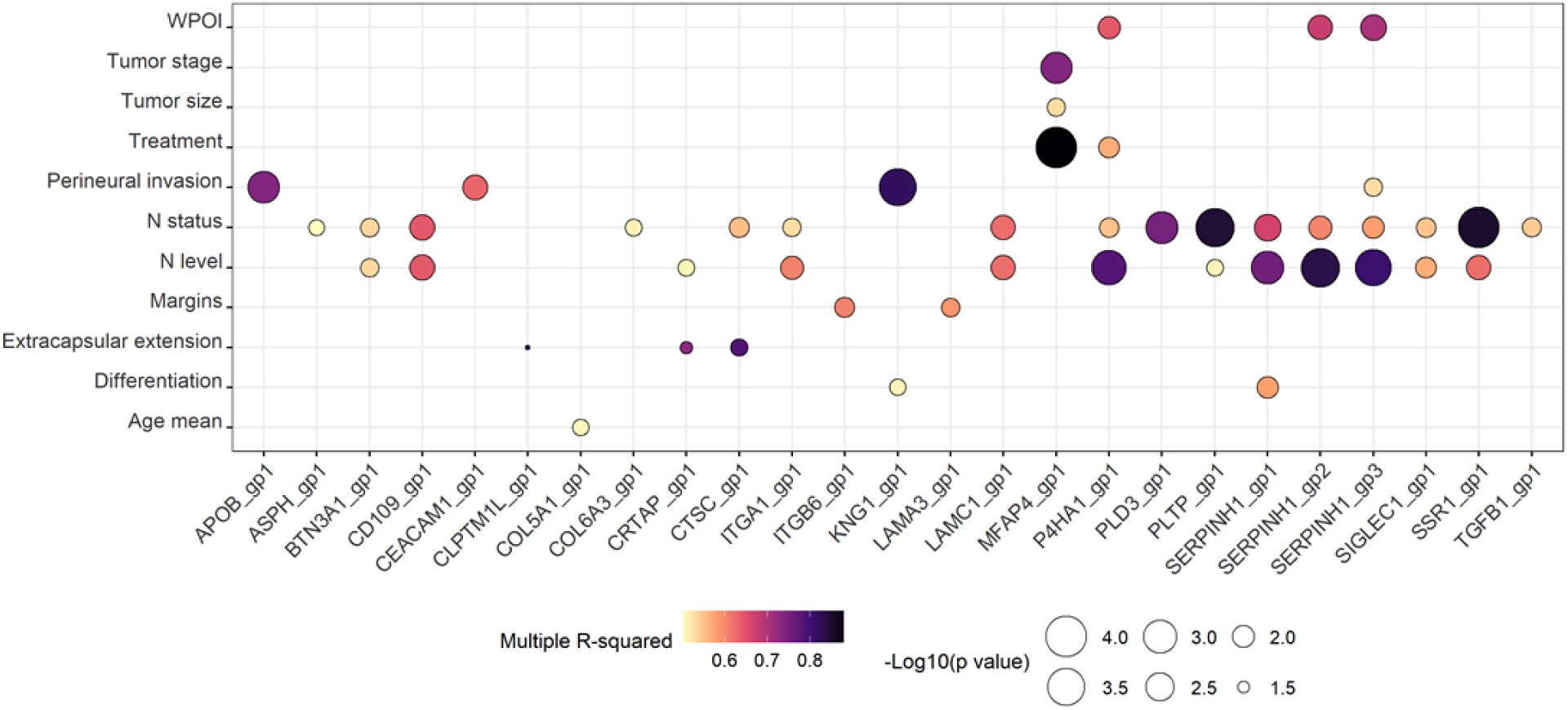
Association of specific *N*-glycopeptides with distinct clinicopathological features. In total 25 of the 79 *N*-glycopeptides that displayed differential abundance between the N0 and N+ patients (plotted on the x axis) were found to associate with a range of clinicopathological features (plotted on the y axis) as measured using Pearson correlation (R). Minimum correlation coefficient +0.7/−0.7, multiple R^2^ > 0.5.

**Supplementary Figure S9.**
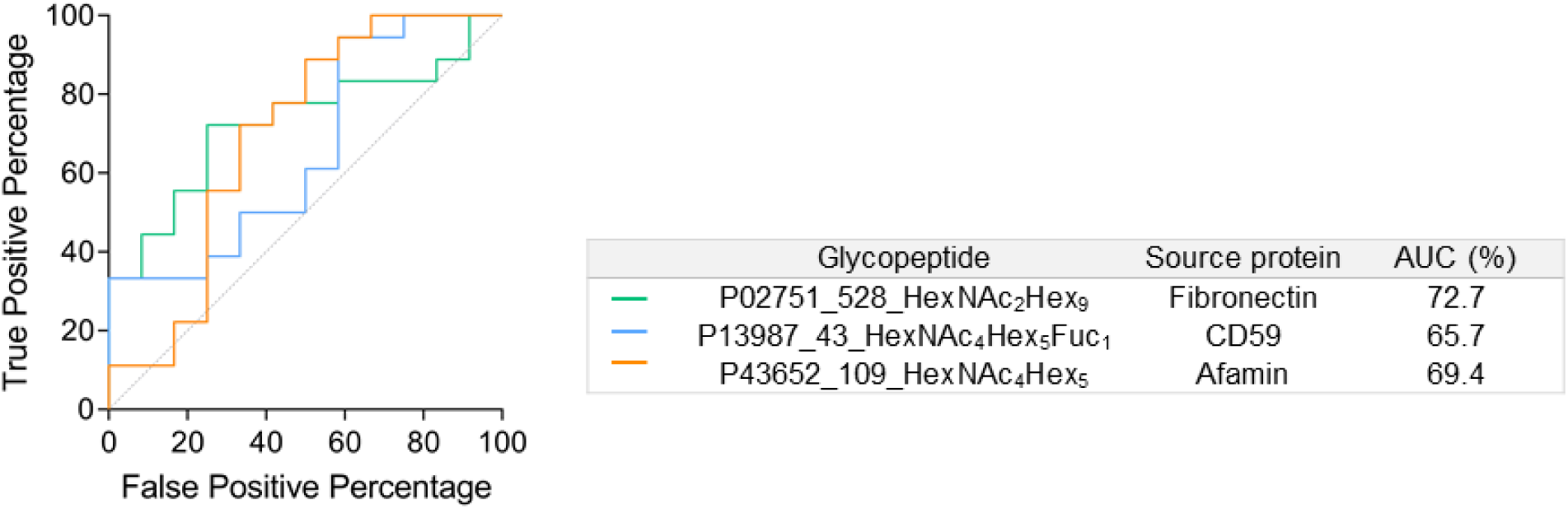
ROC analysis of three *N*-glycopeptides associated with patient survival. *N*-glycopeptides that displayed differential abundance between the N0 and N+ patients and associations with patient survival were evaluated for their potential to stratify patients with and without lymph node metastasis using a logistic regression model. Three *N*-glycopeptides from three different source glycoproteins (fibronectin, green trace; CD59, blue trace; afamin, orange trace) displayed an AUC-ROC > 60%. AUC-ROC: area-under-the-curve of the receiver operating characteristic.

**Supplementary Table S1-8.** (provided as a separate .xlsx file) Information of data composition and analysis.

**Supplementary File S1.** (provided as a separate .pptx file)

Spectral evidence of the reported *N*-glycans observed from the OSCC tissue samples.

## Supplementary File S1

### Annotation and Fragmentation Key

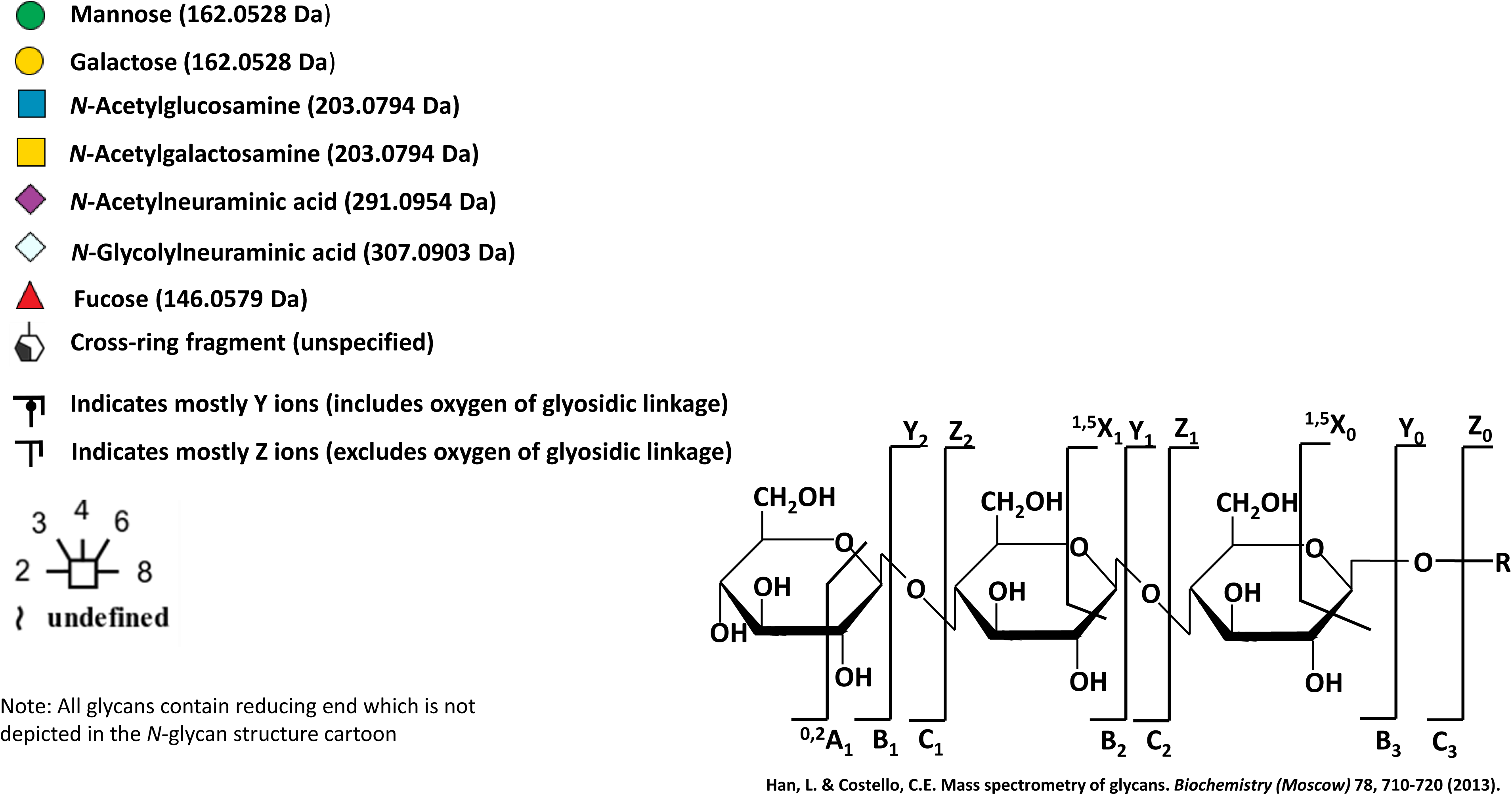

### Spectral evidence for the reported OSCC tumour tissue *N*-glycans

**The *N*-glycans reported in this paper were characterised and quantified by:**

a. EIC (MS) of monoisotopic precursor ions and AUC integration (blue highlights in insert, from Skyline)
b. PGC-LC retention time
c. CID-MS/MS (-) manual annotation

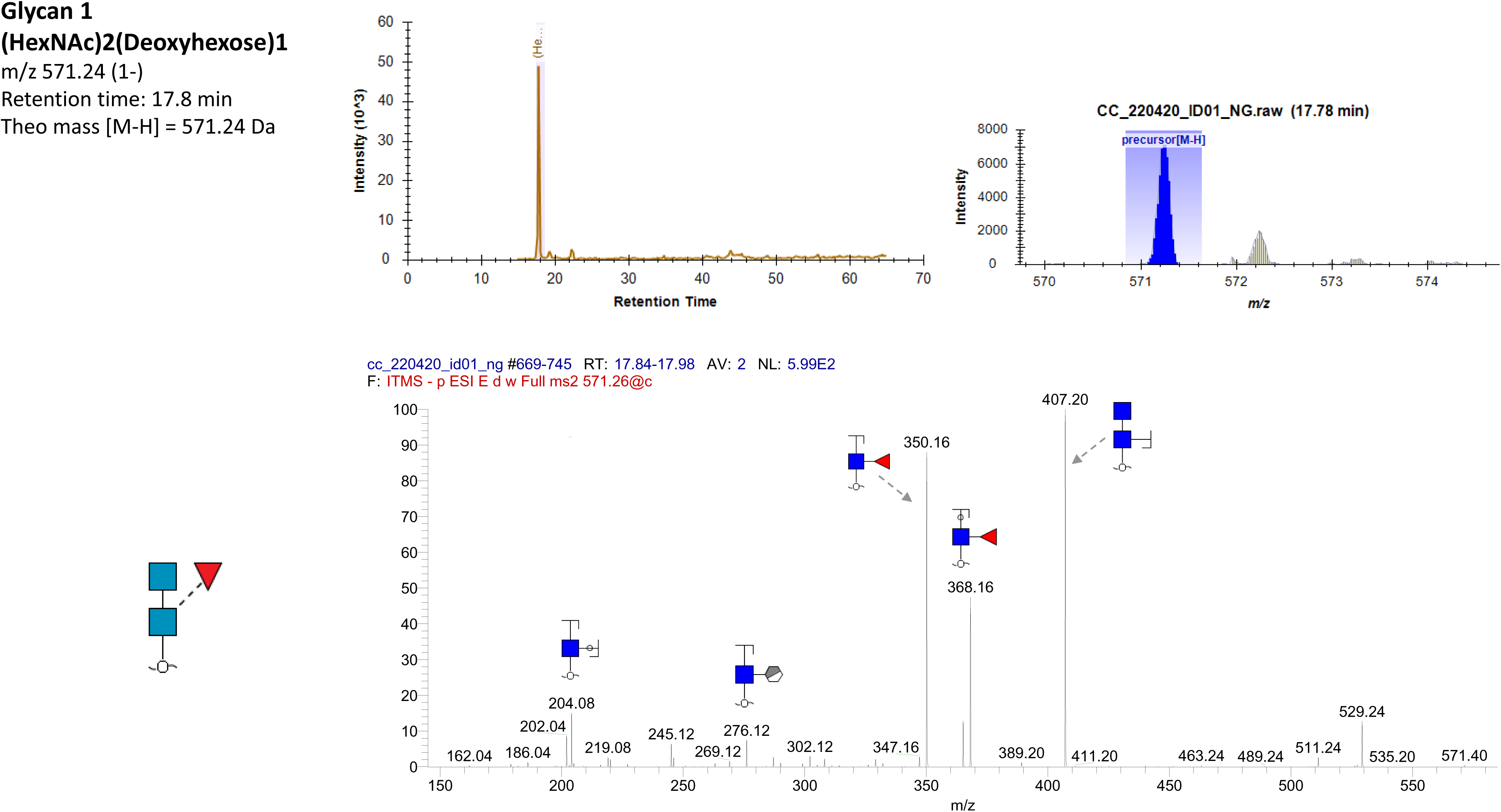
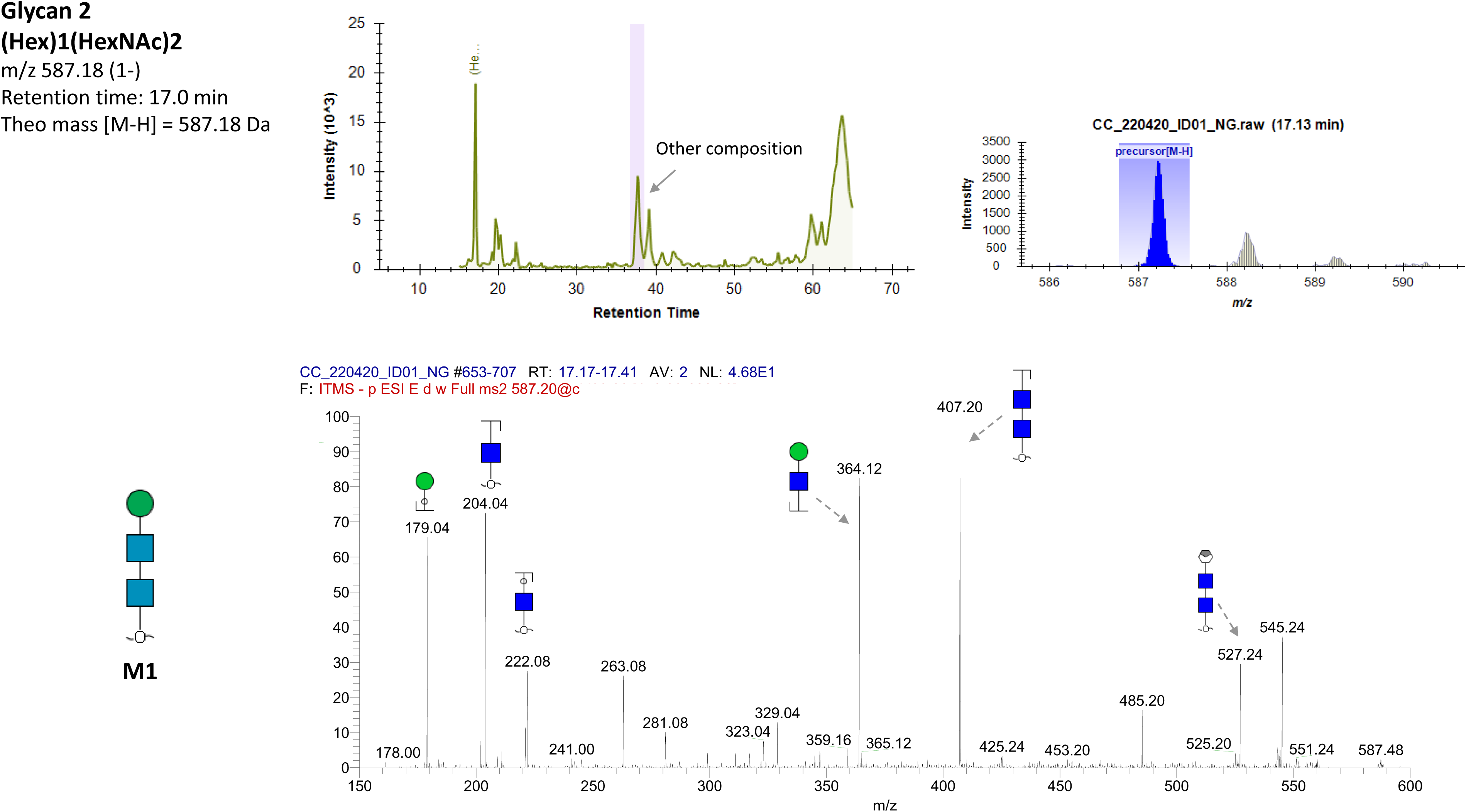
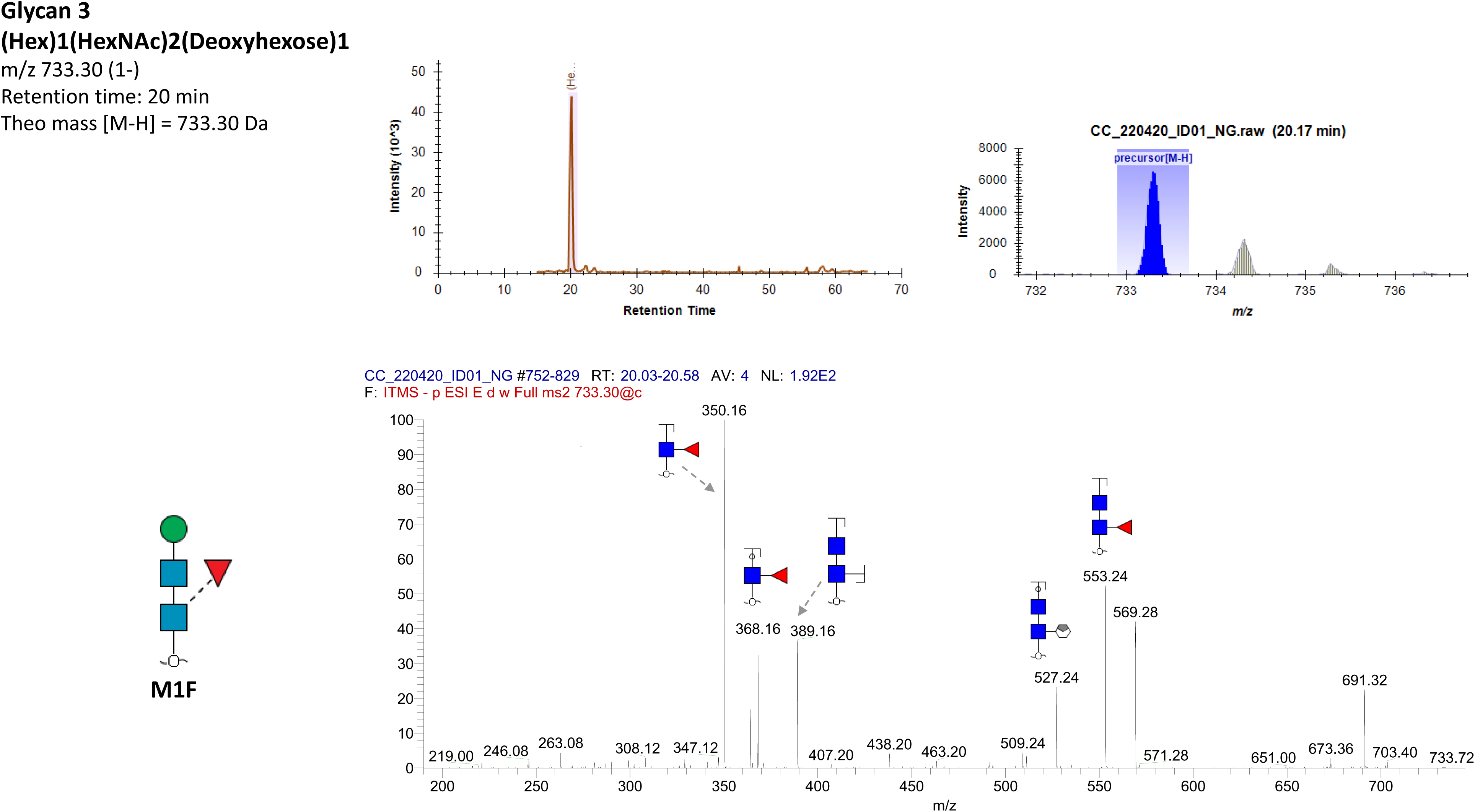
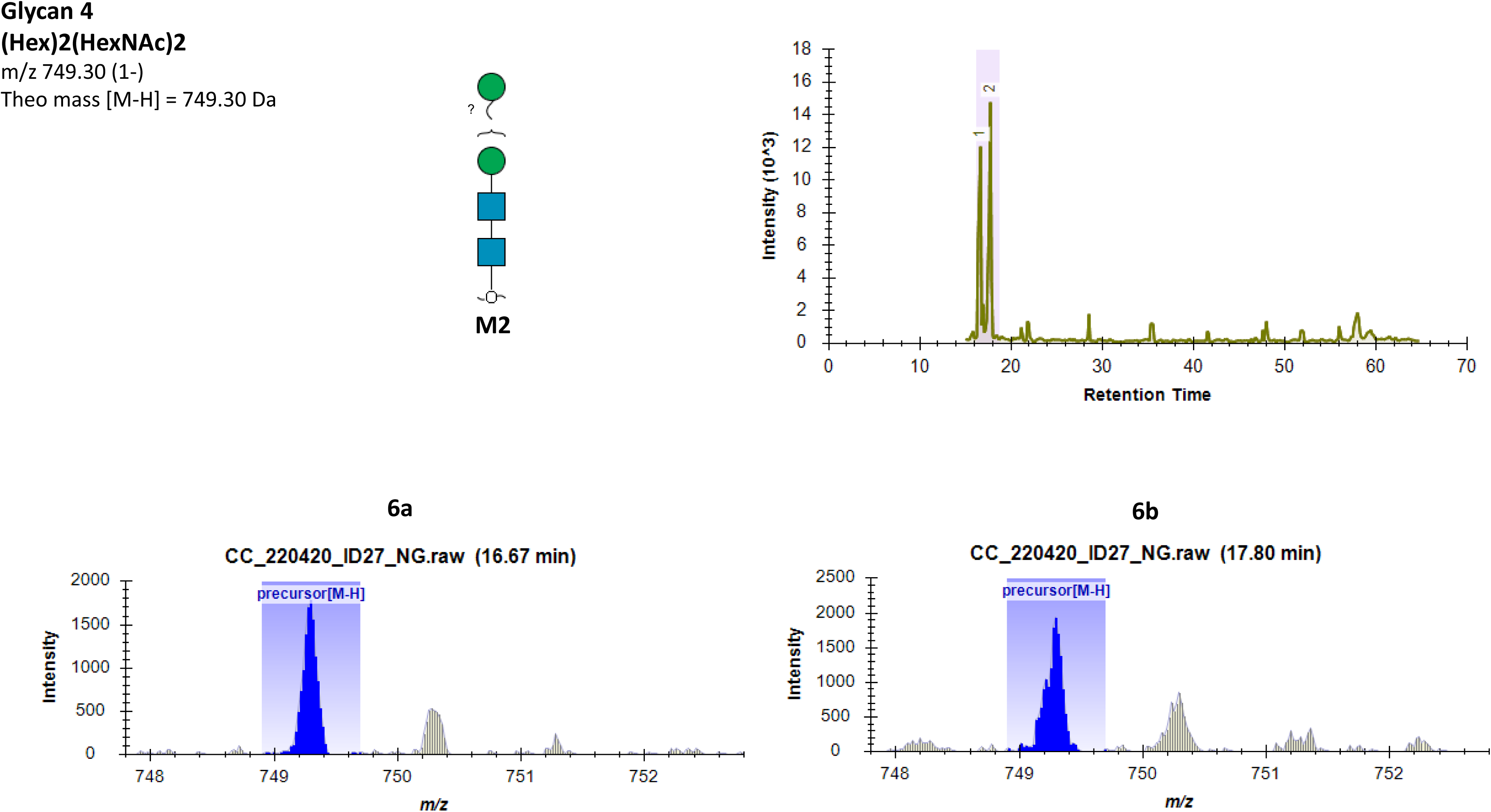
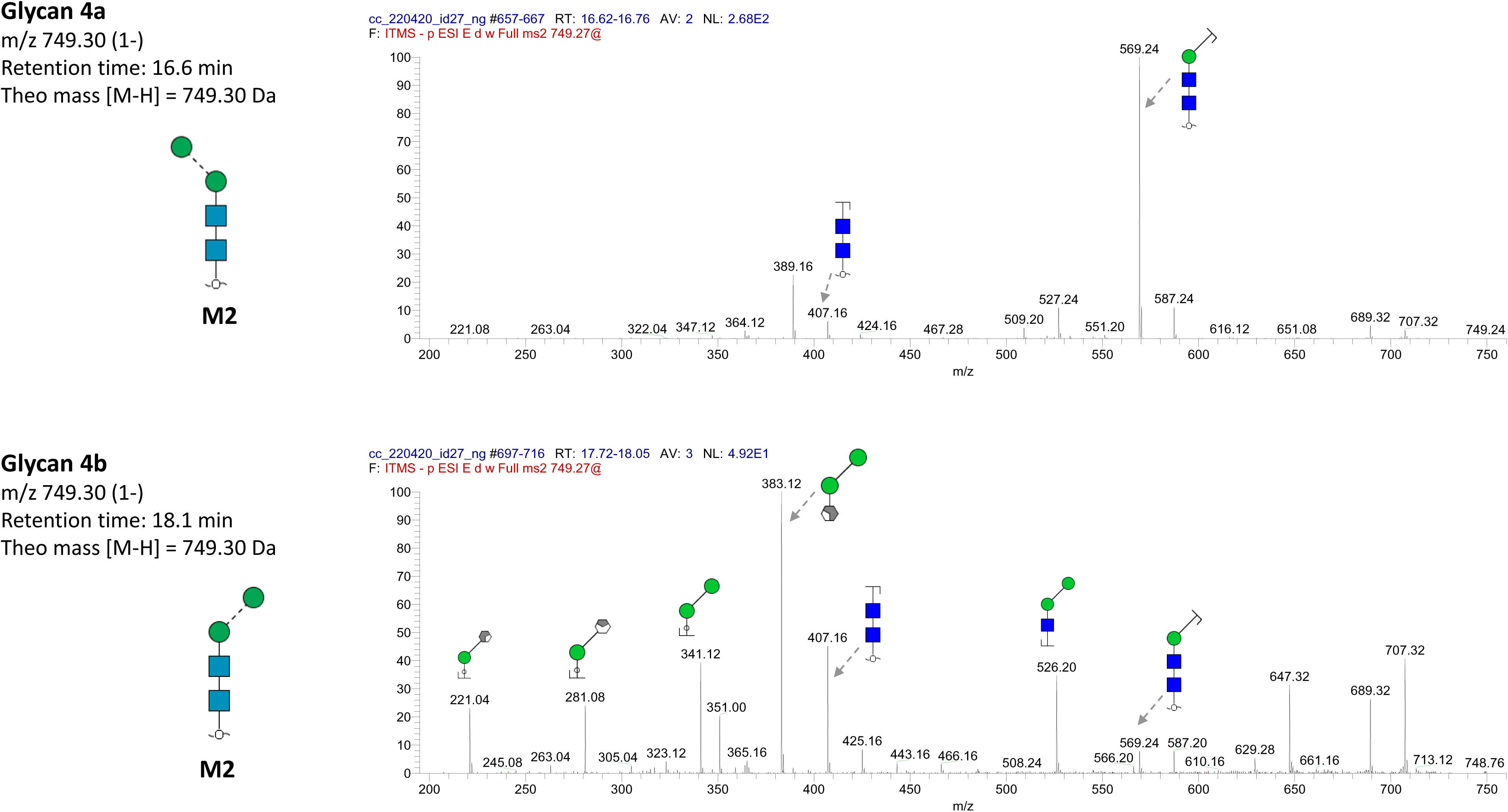
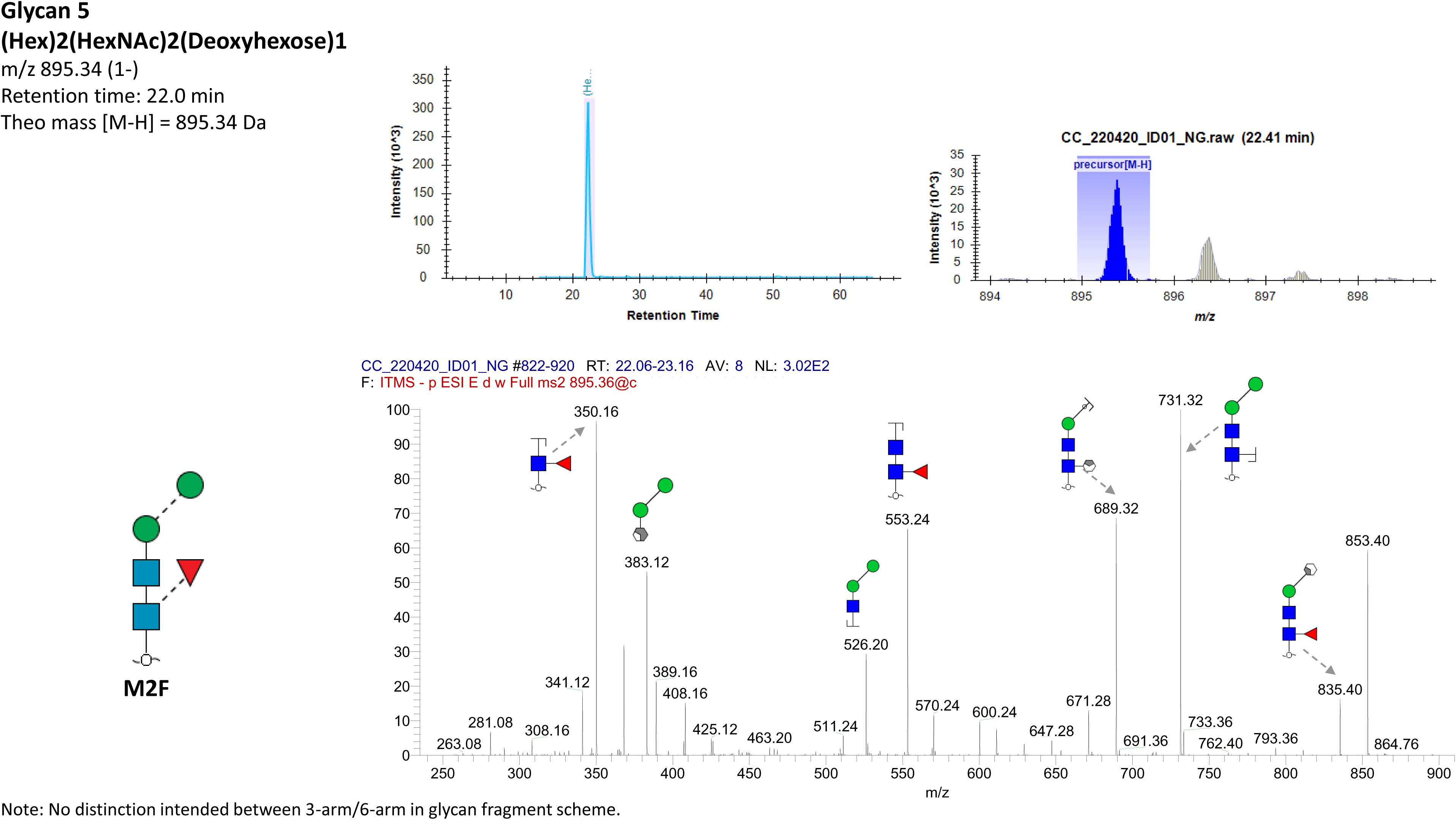
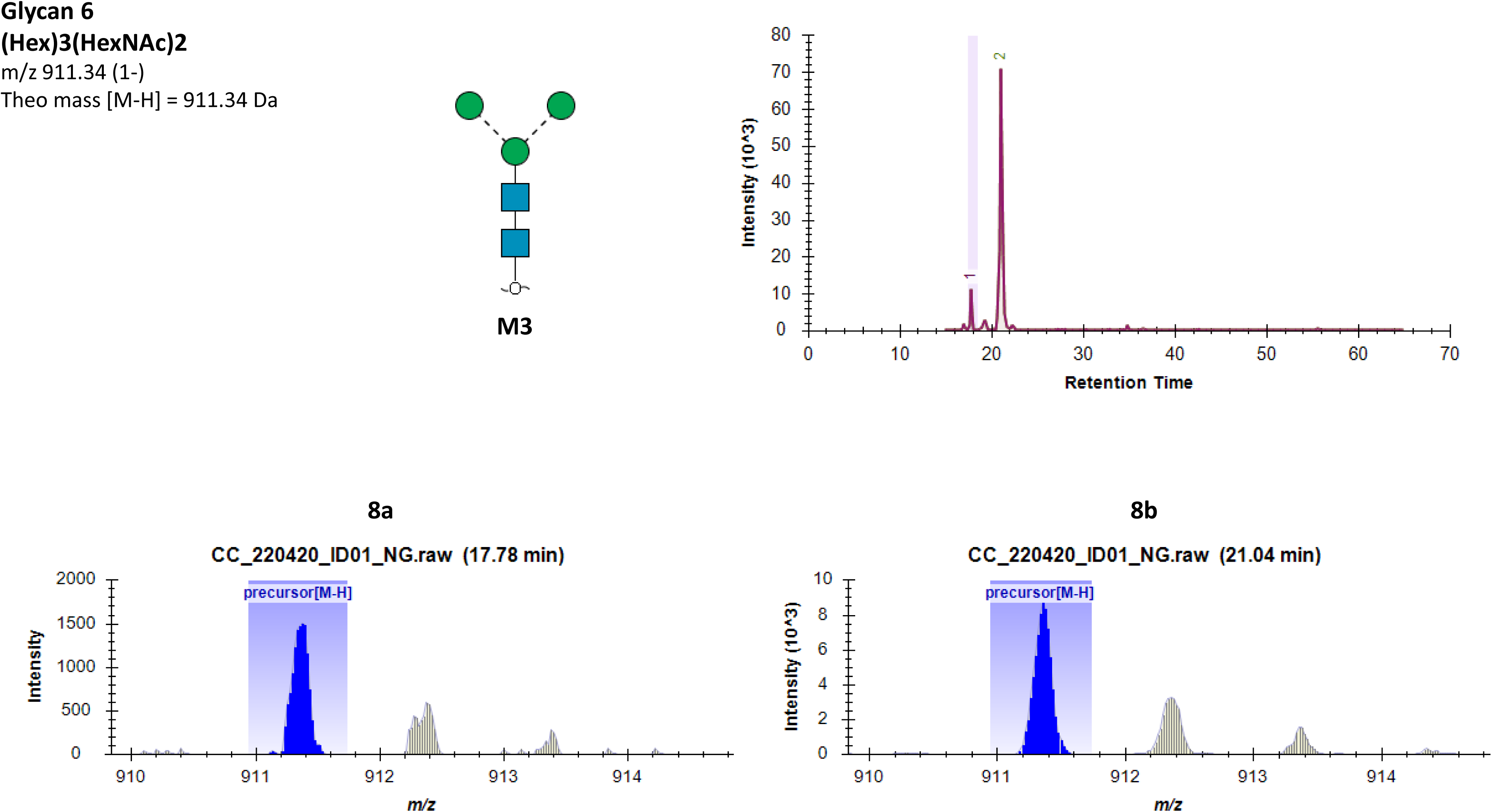
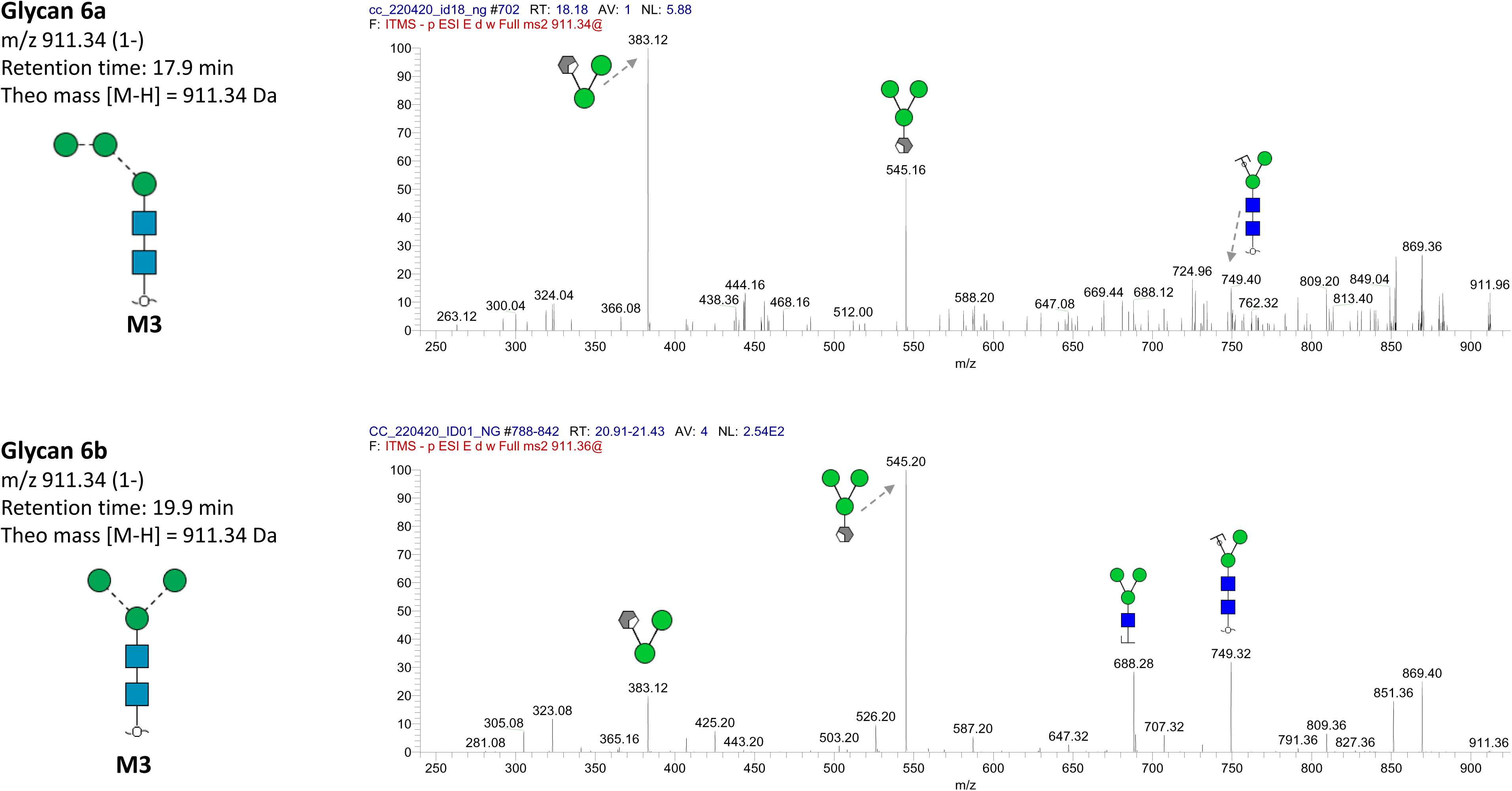
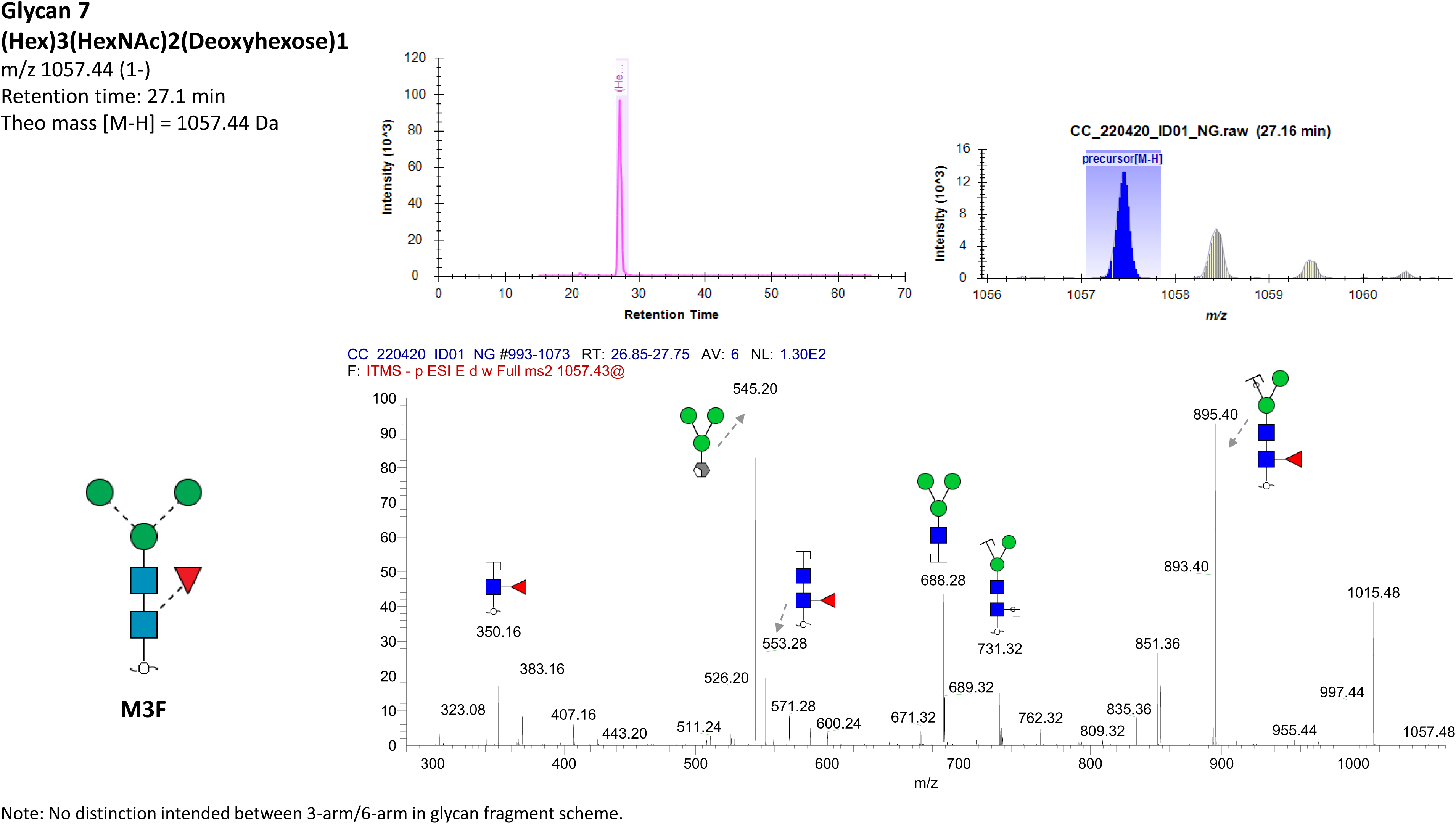
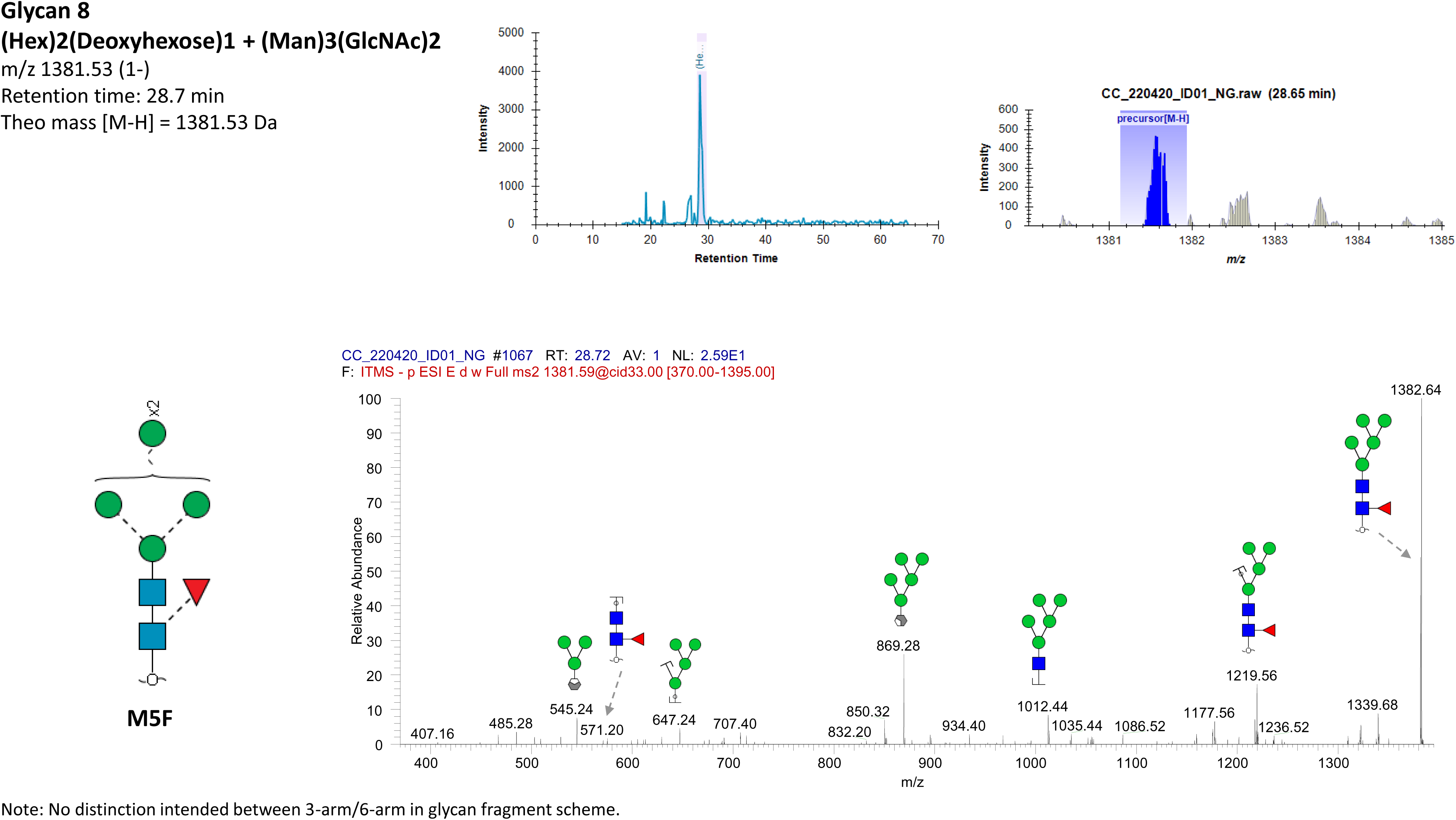
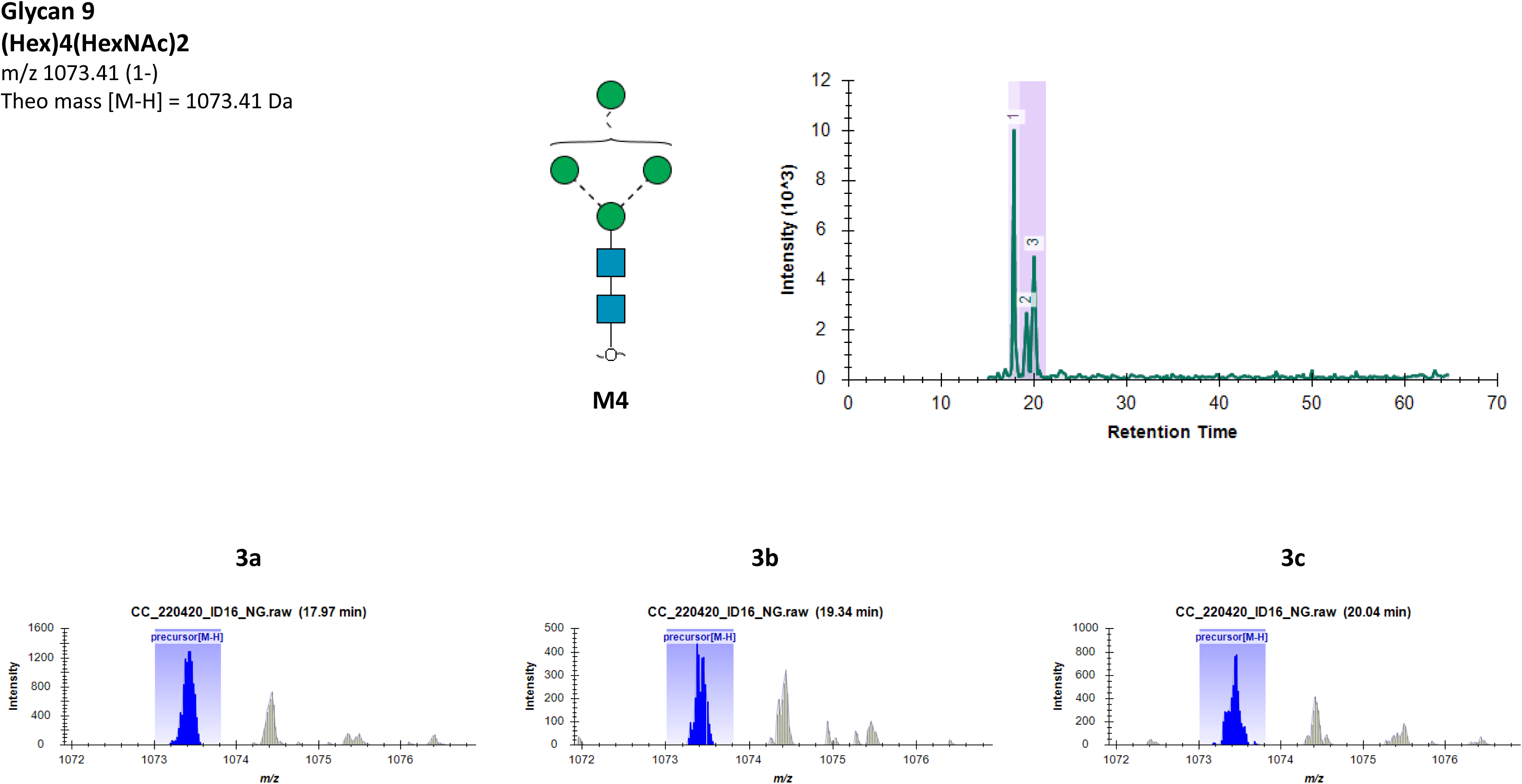
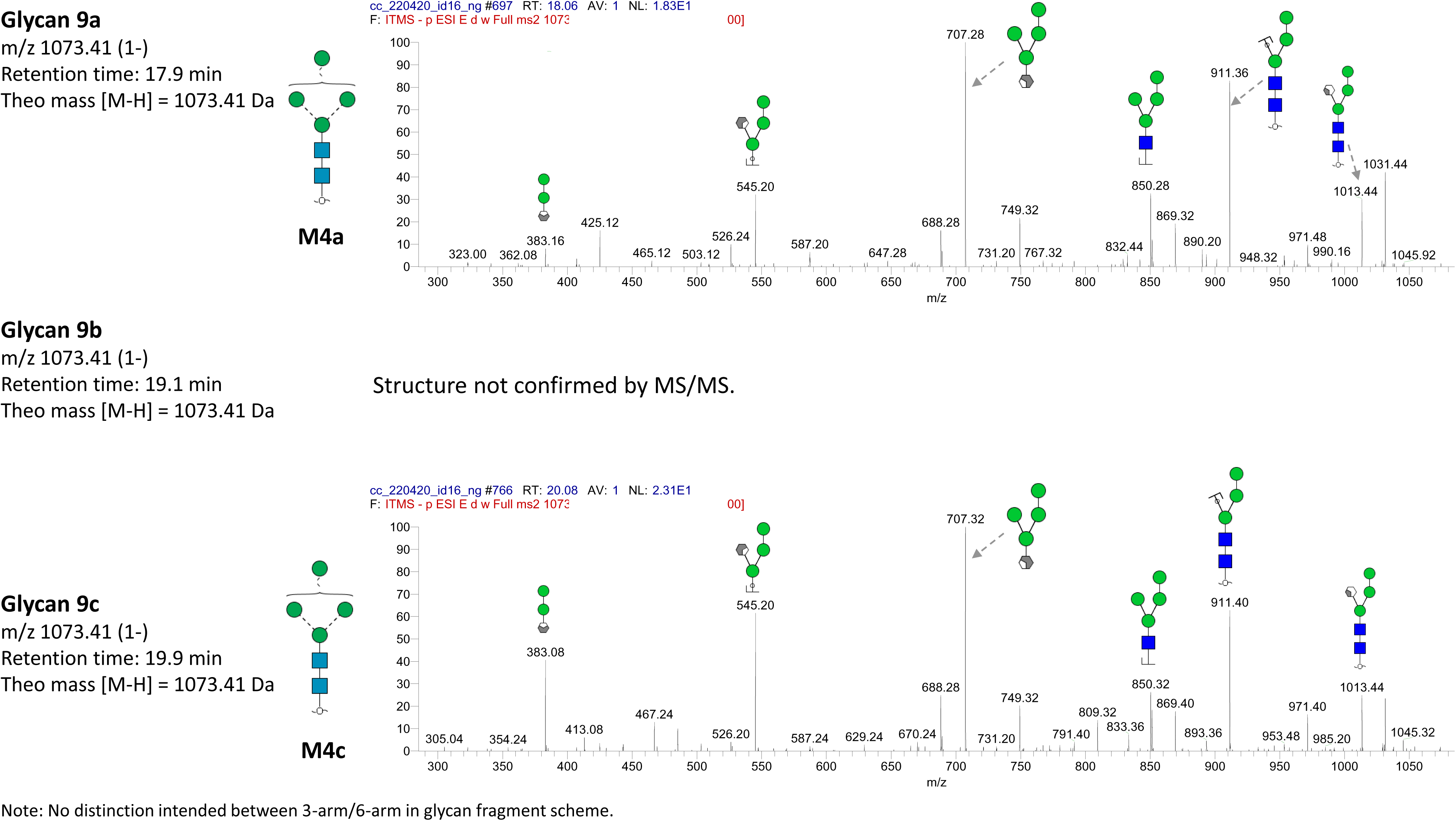
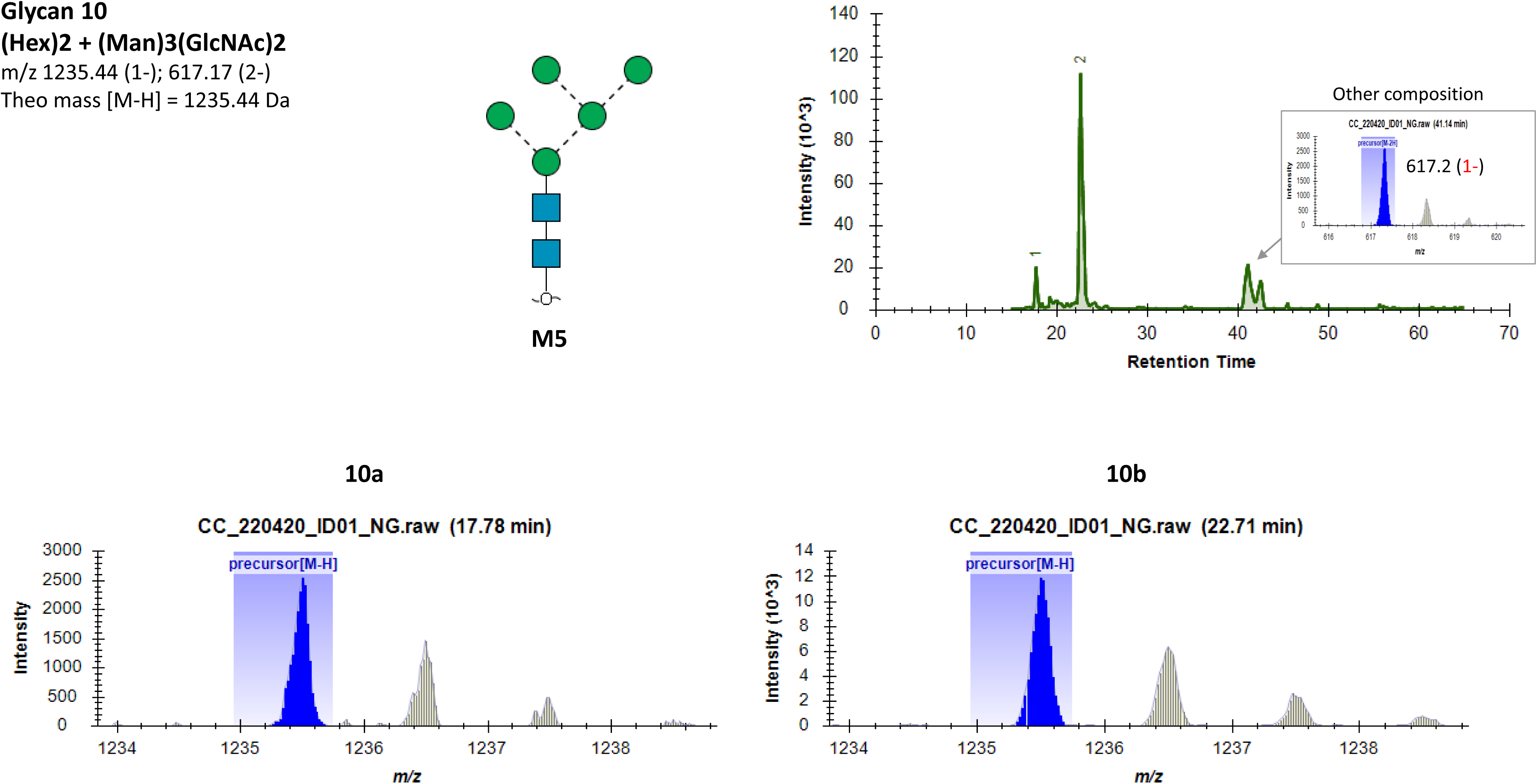
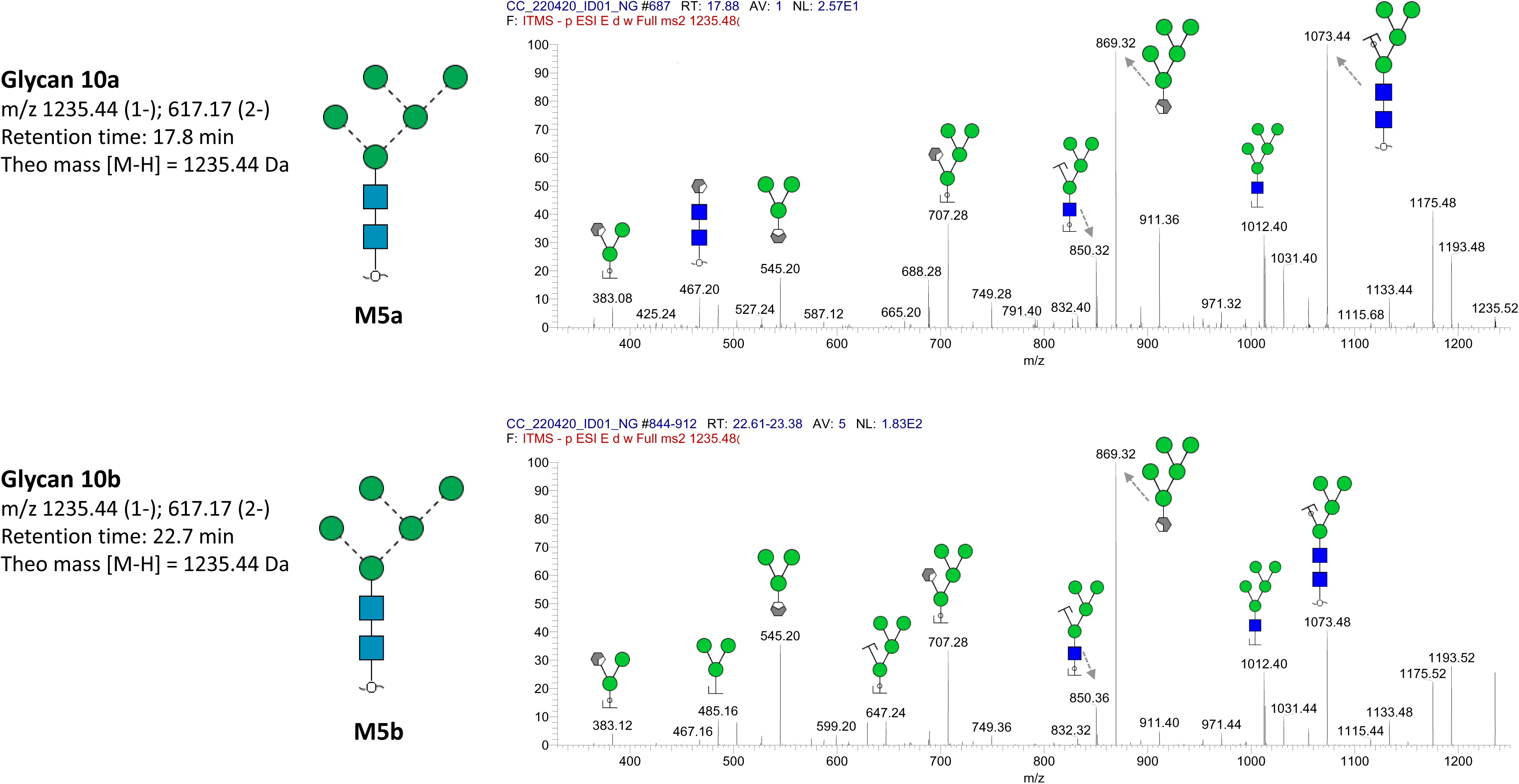
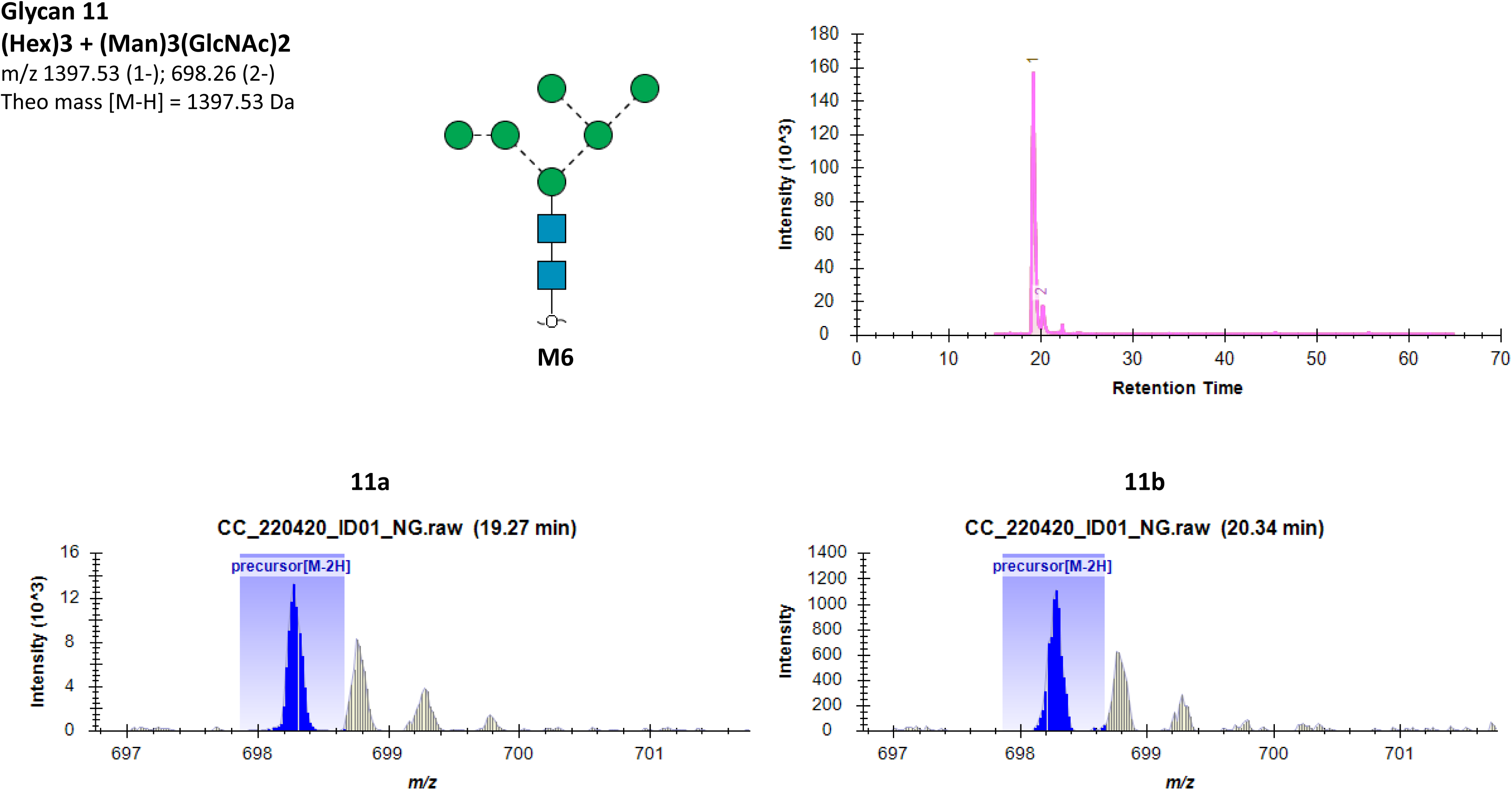
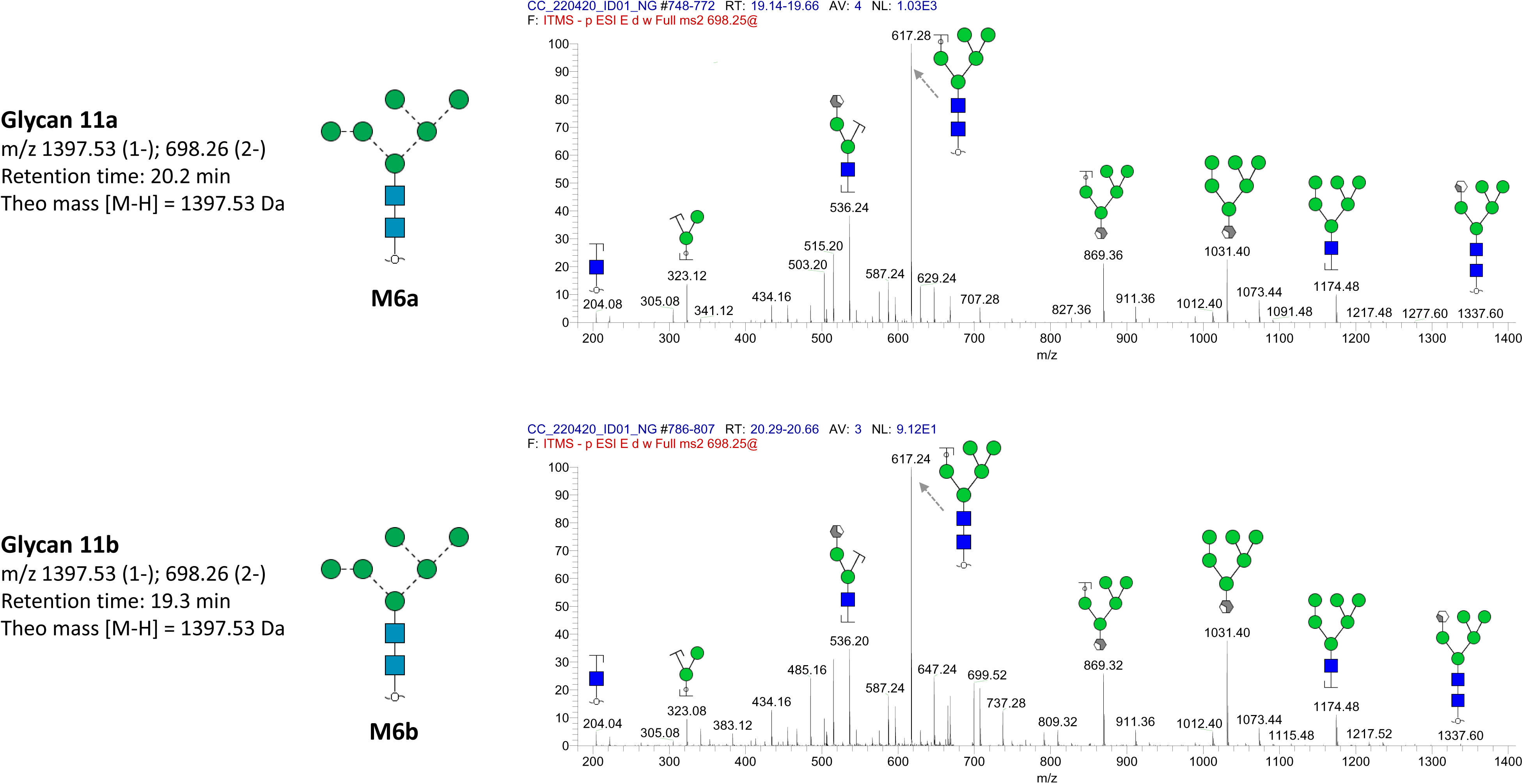
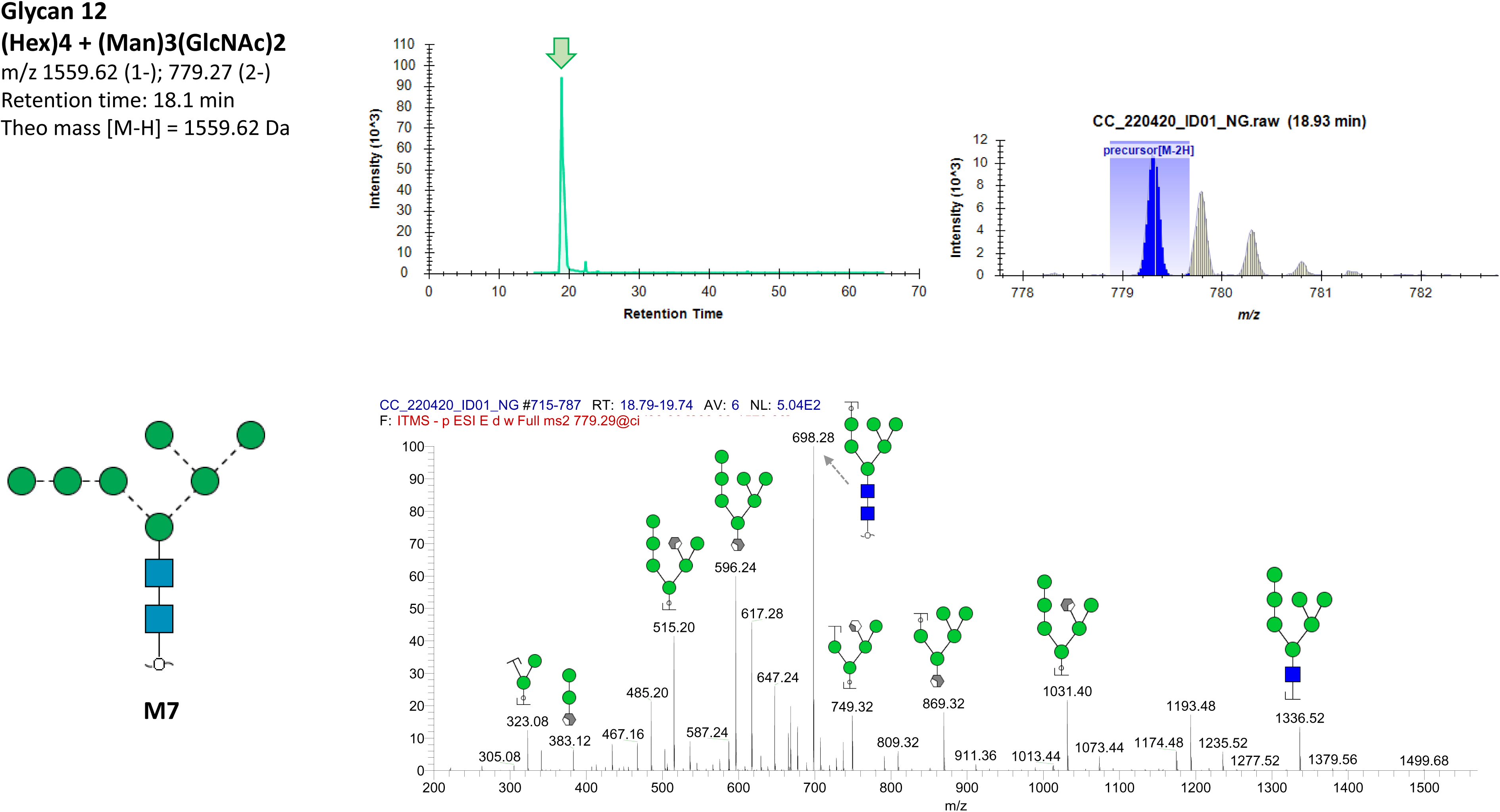
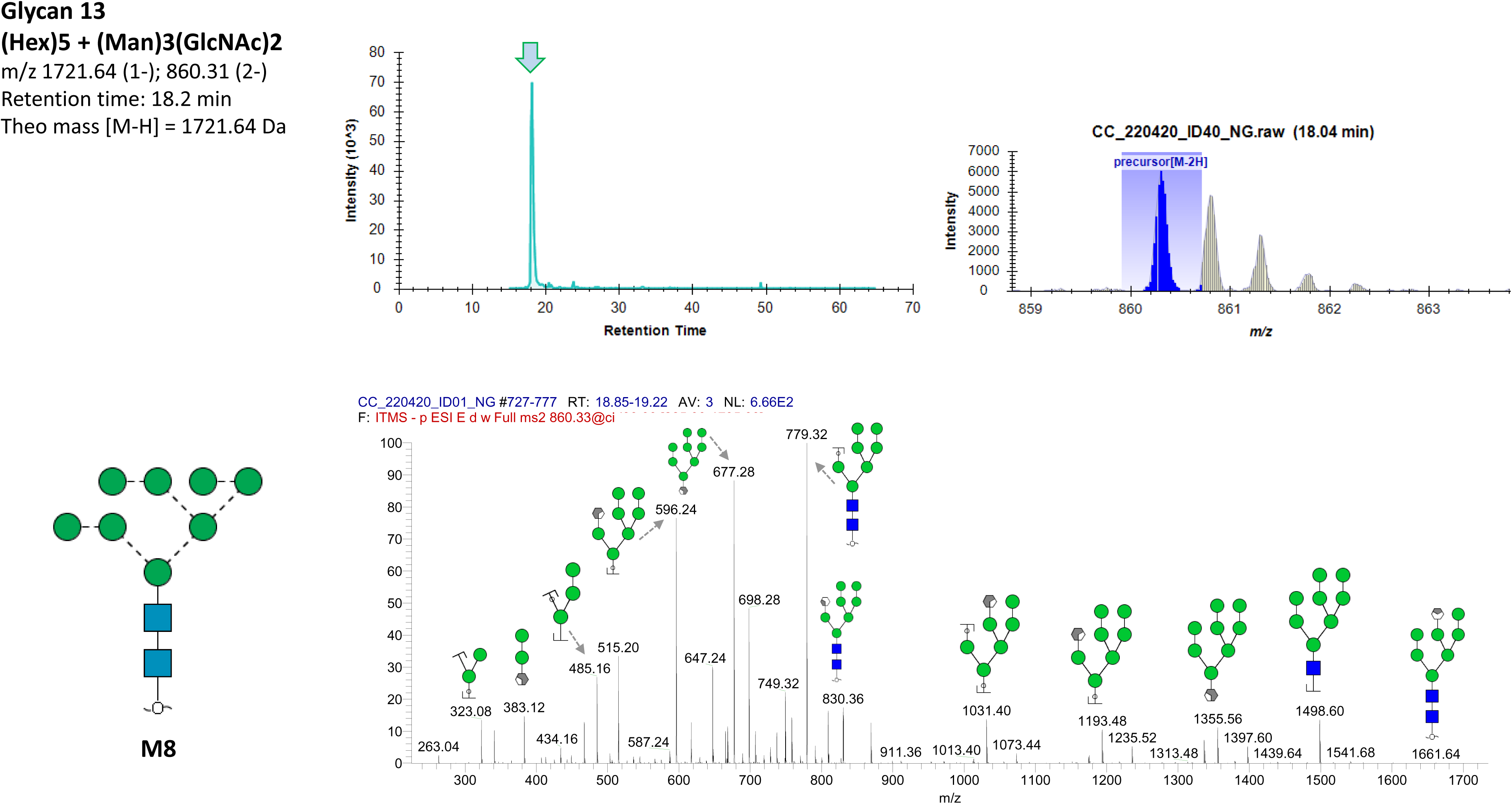
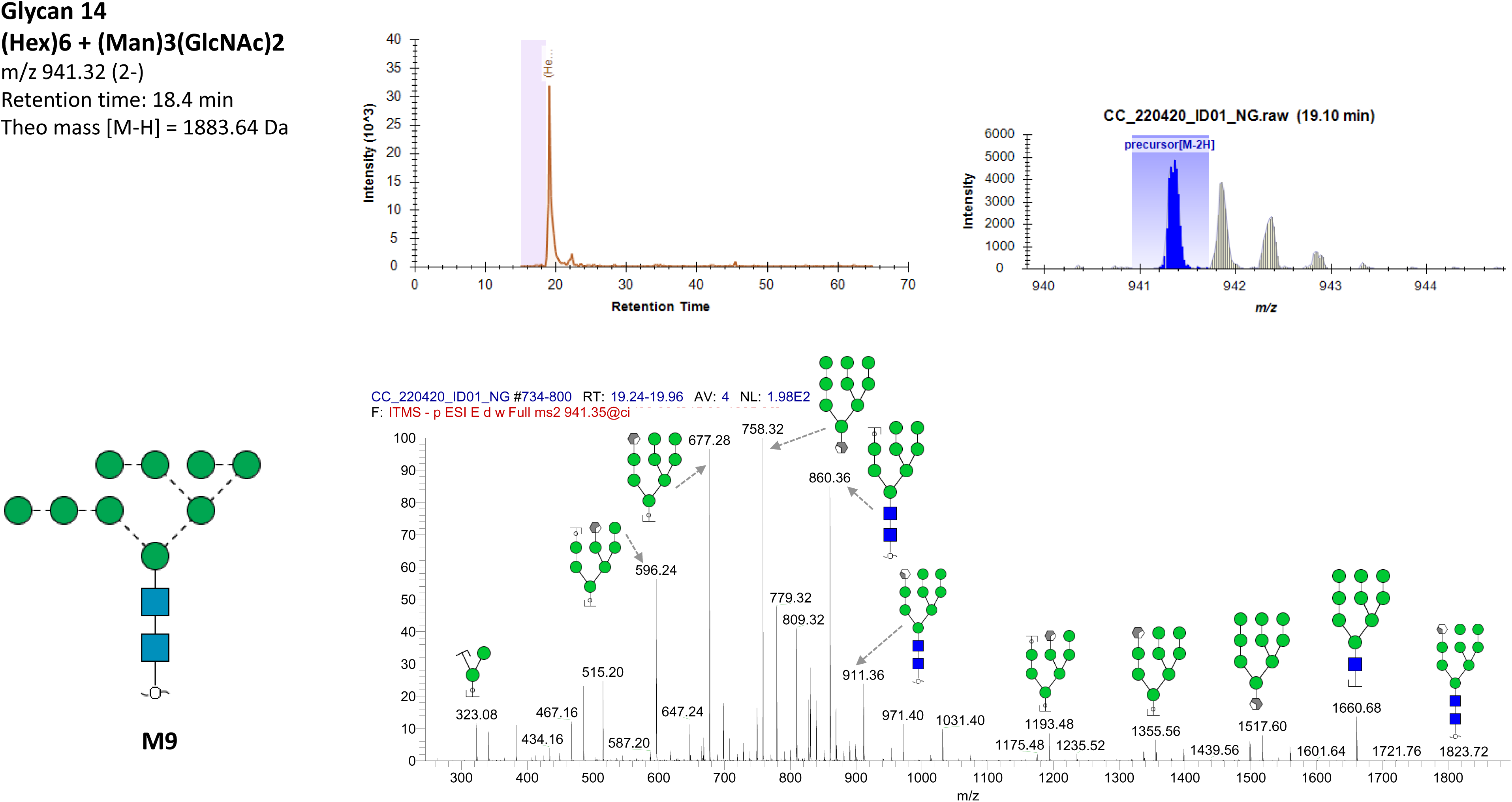
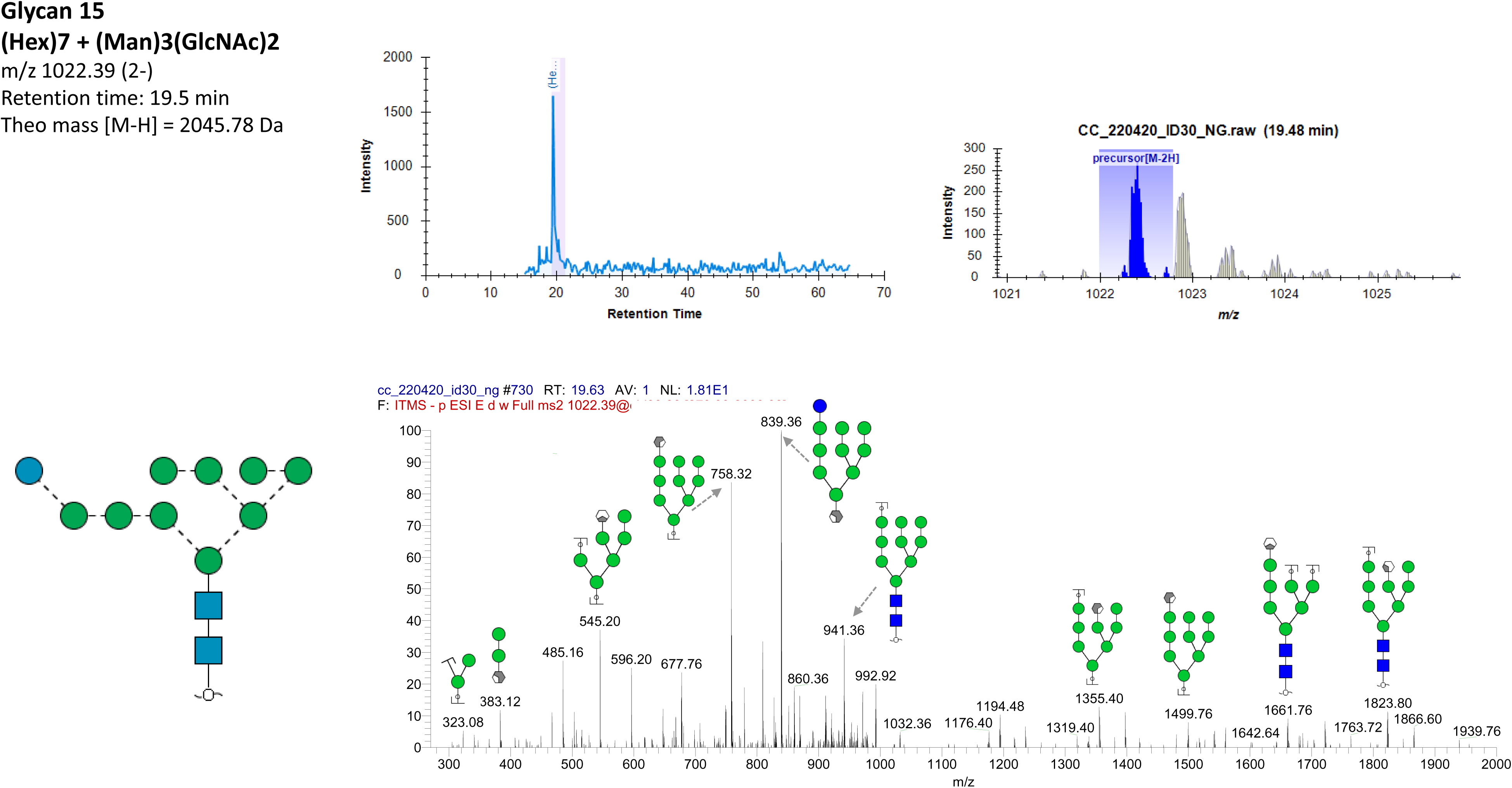
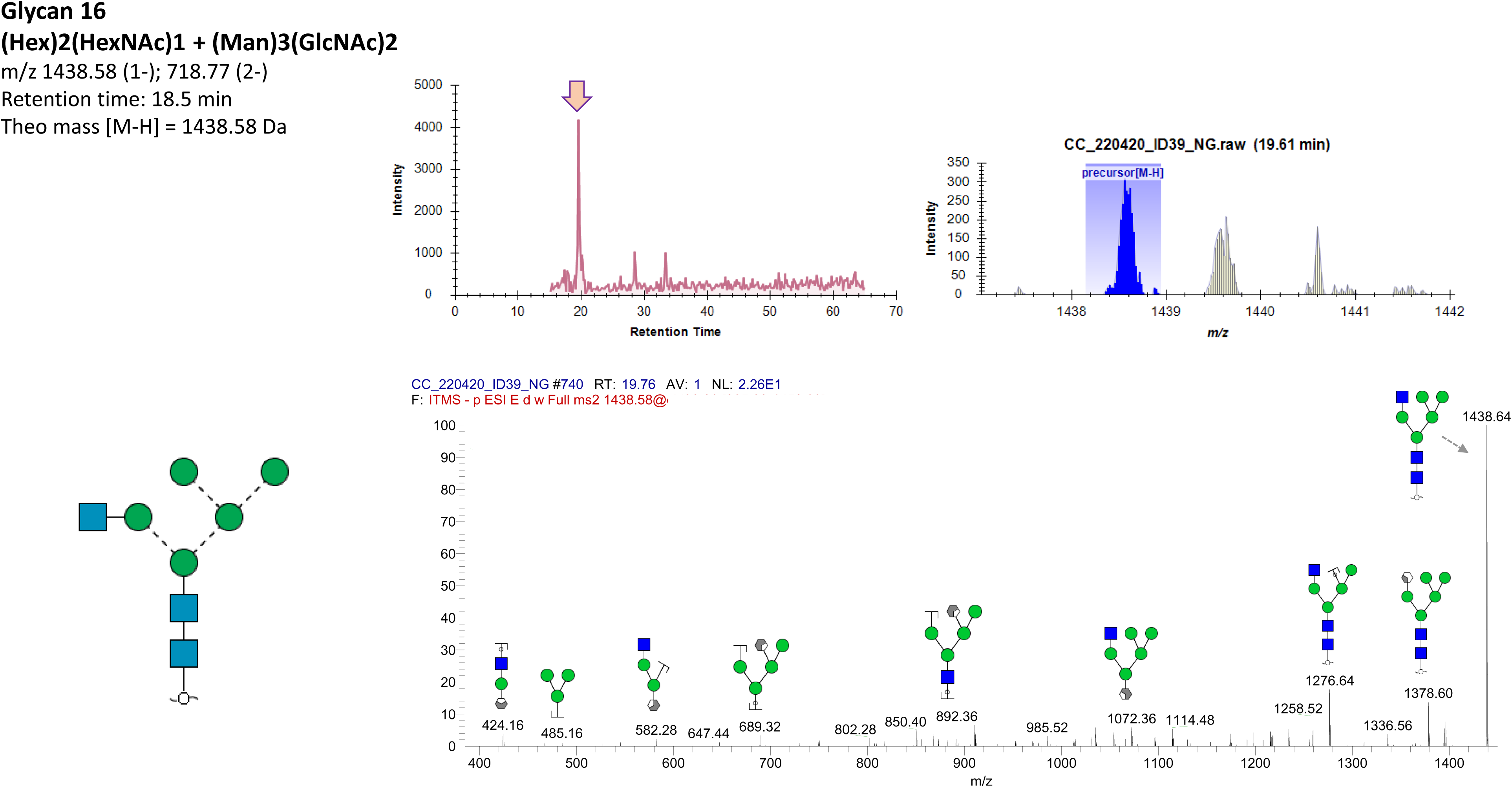
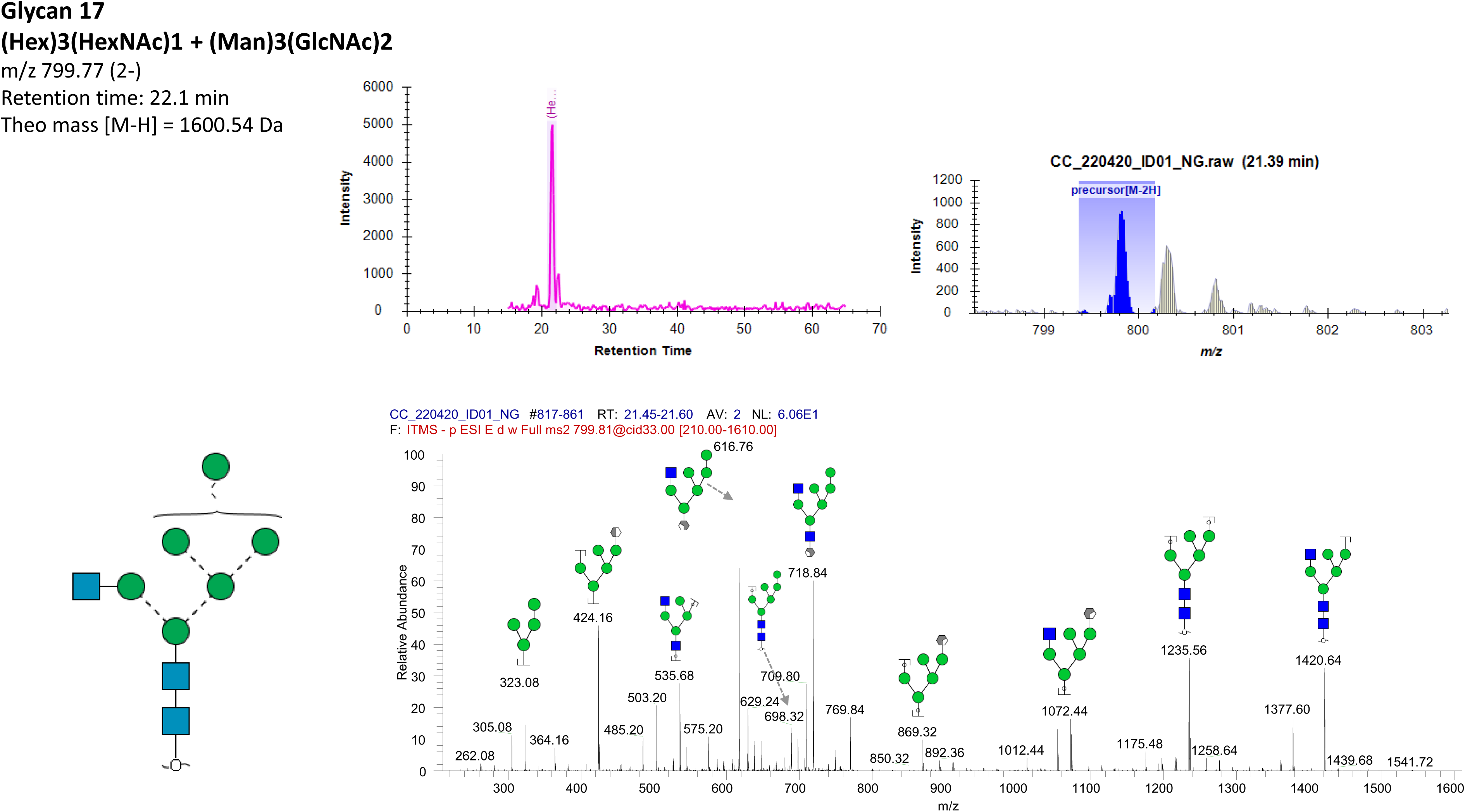
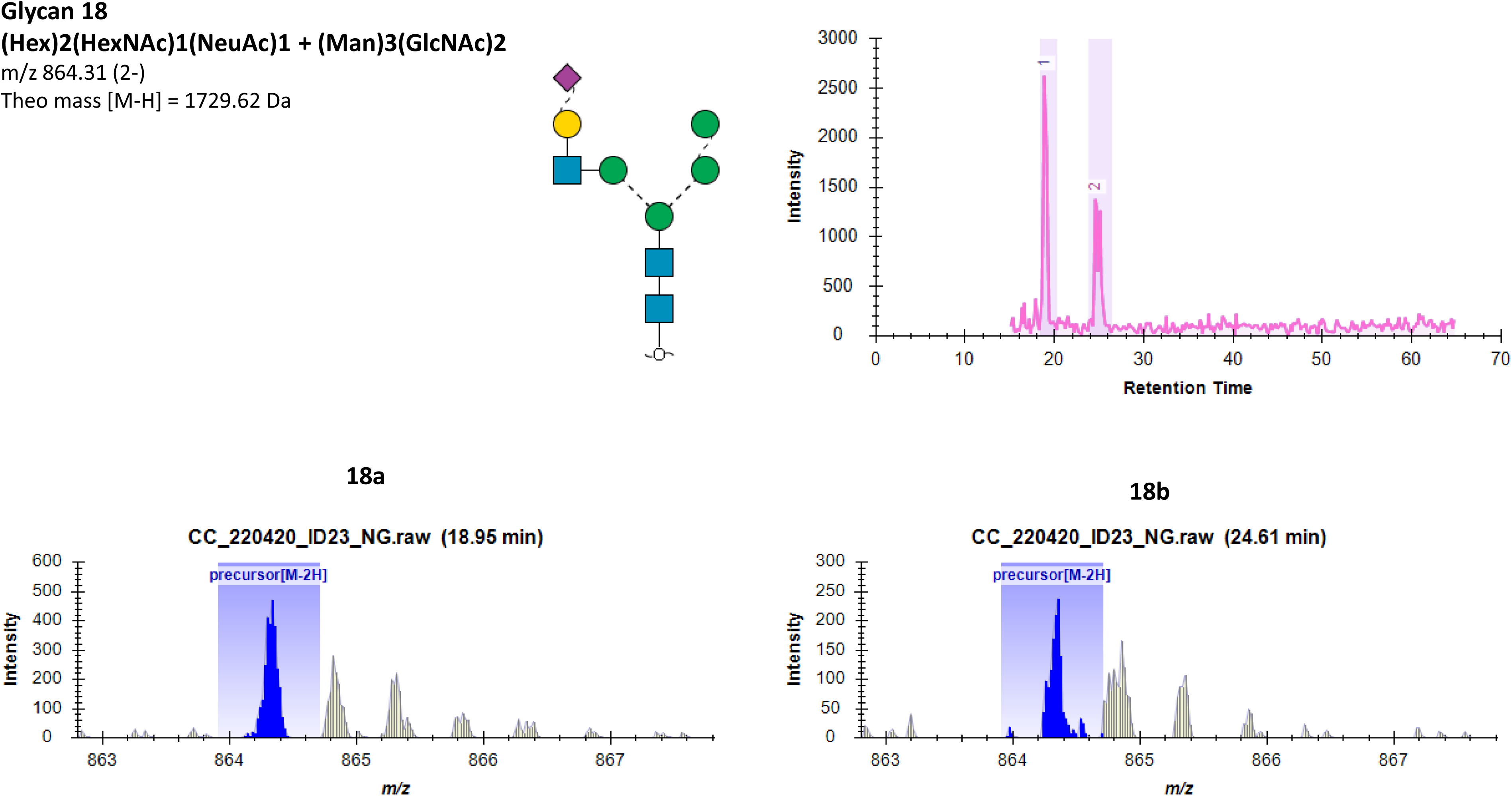
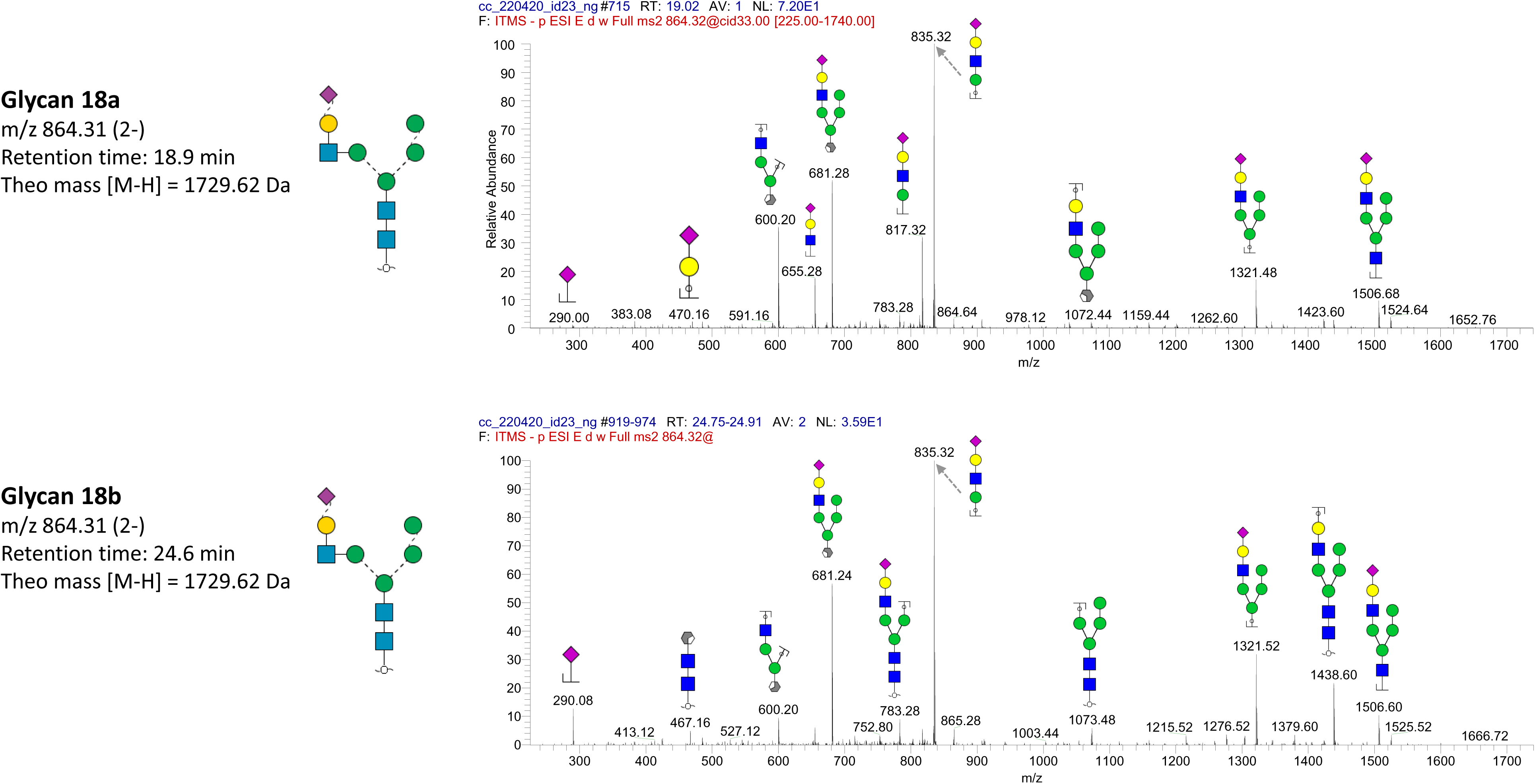
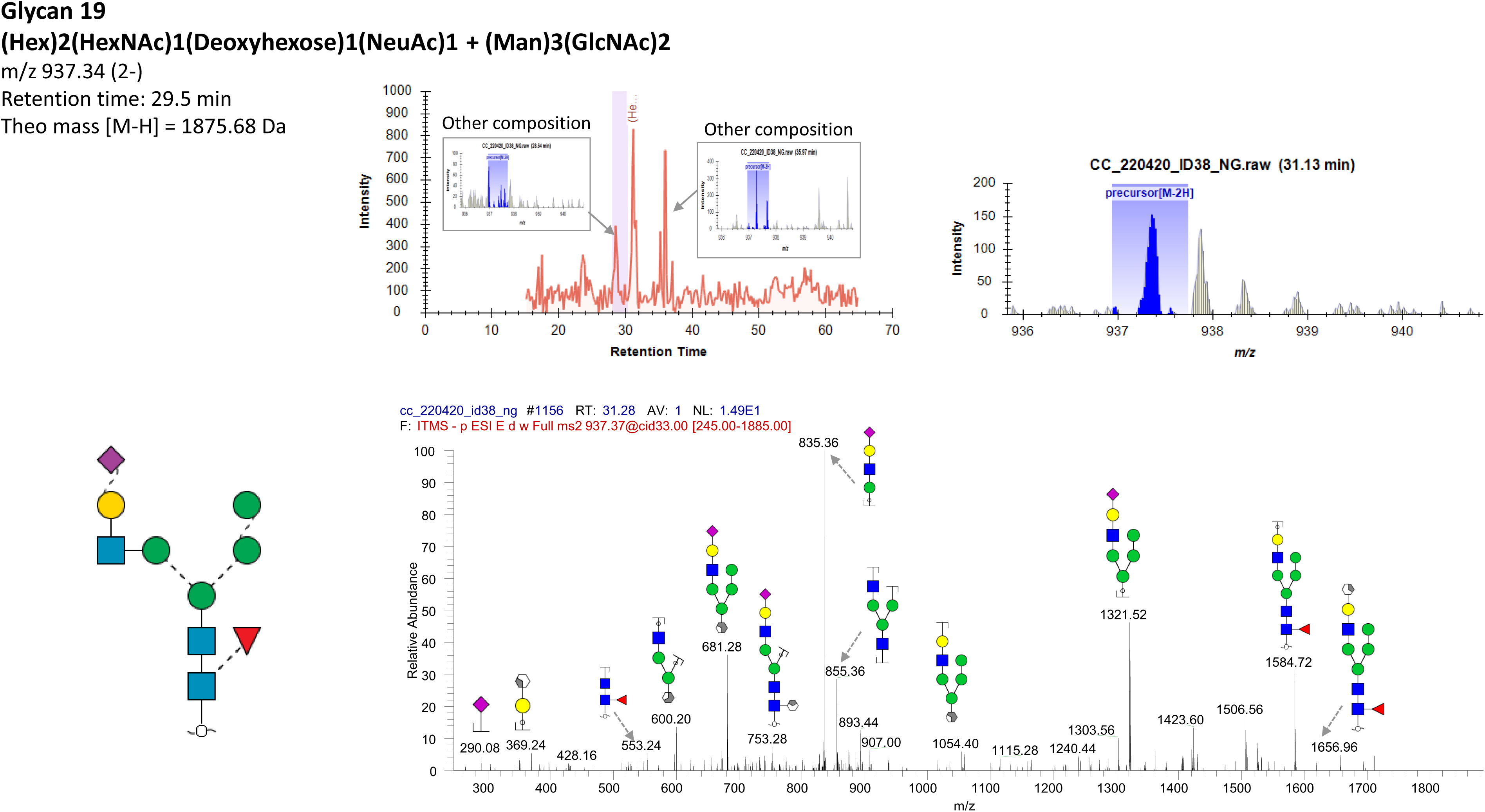
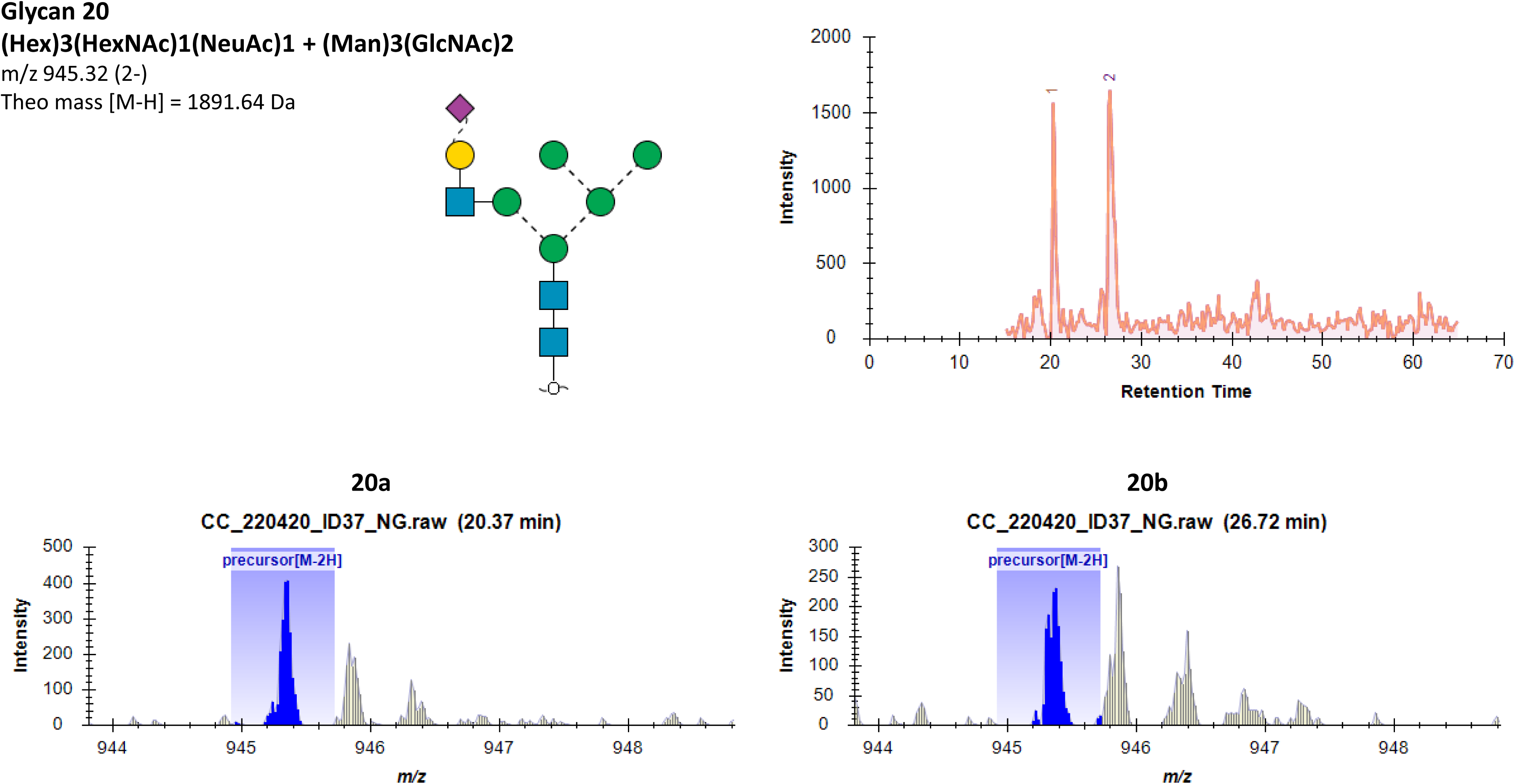
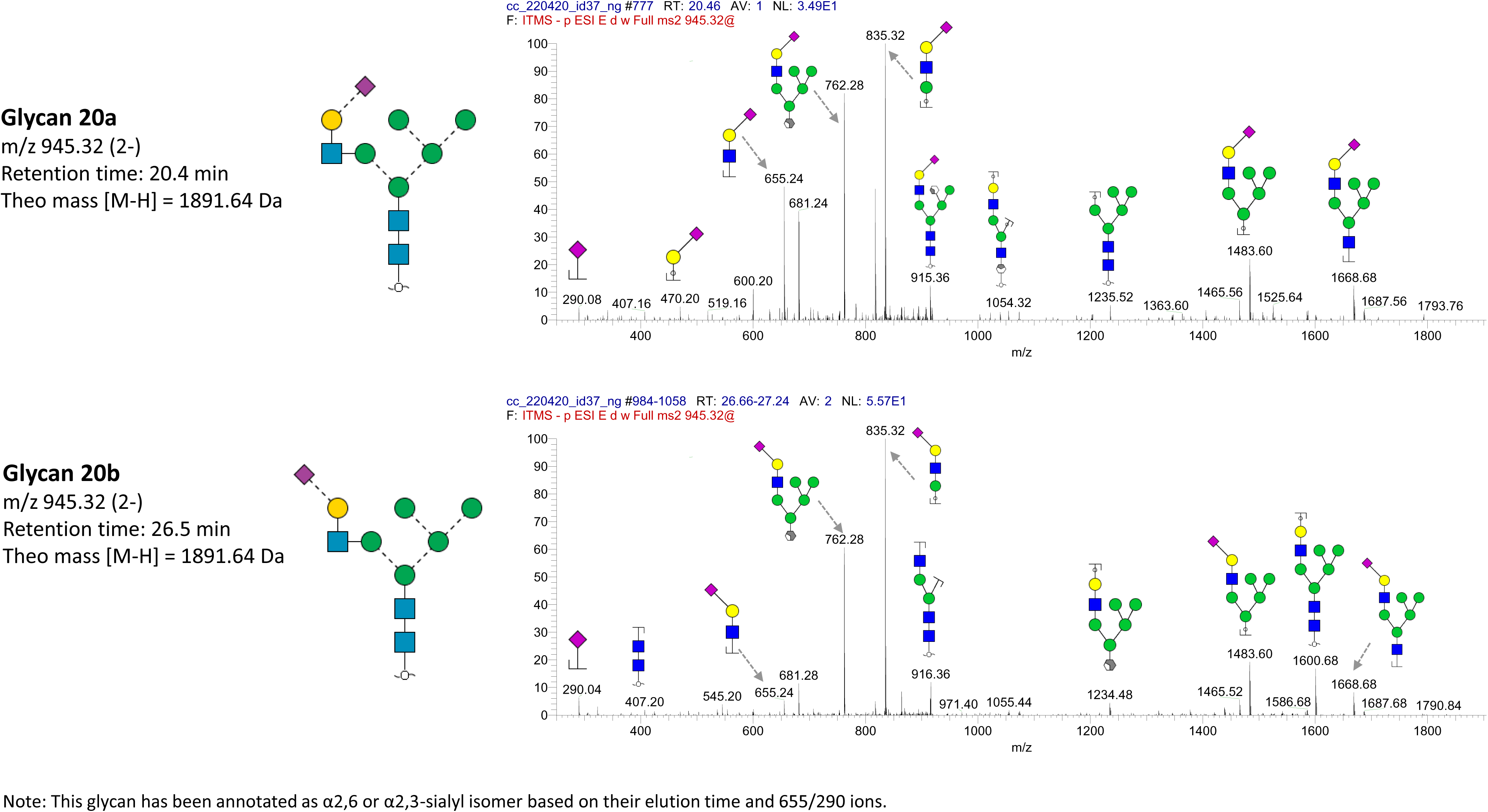
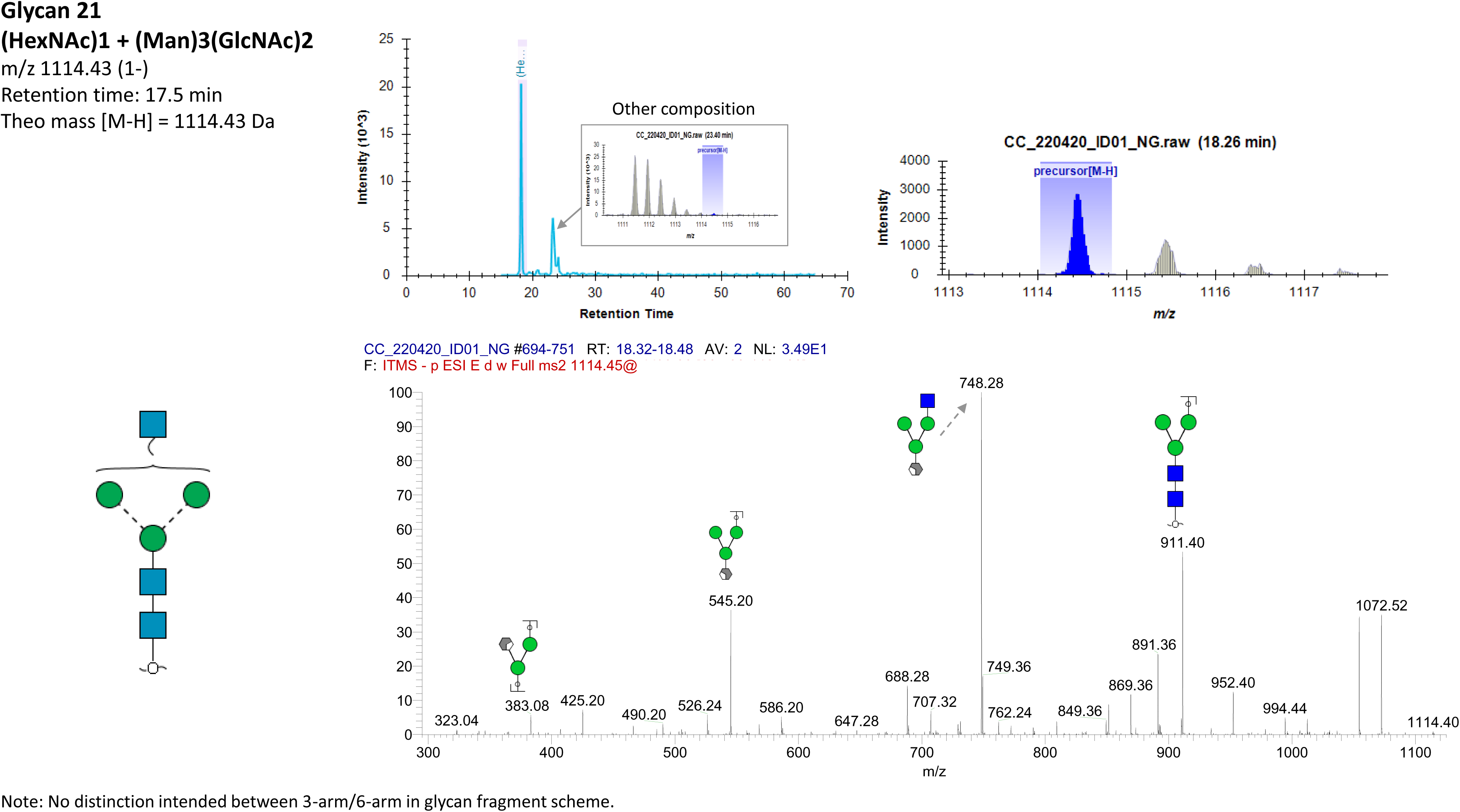
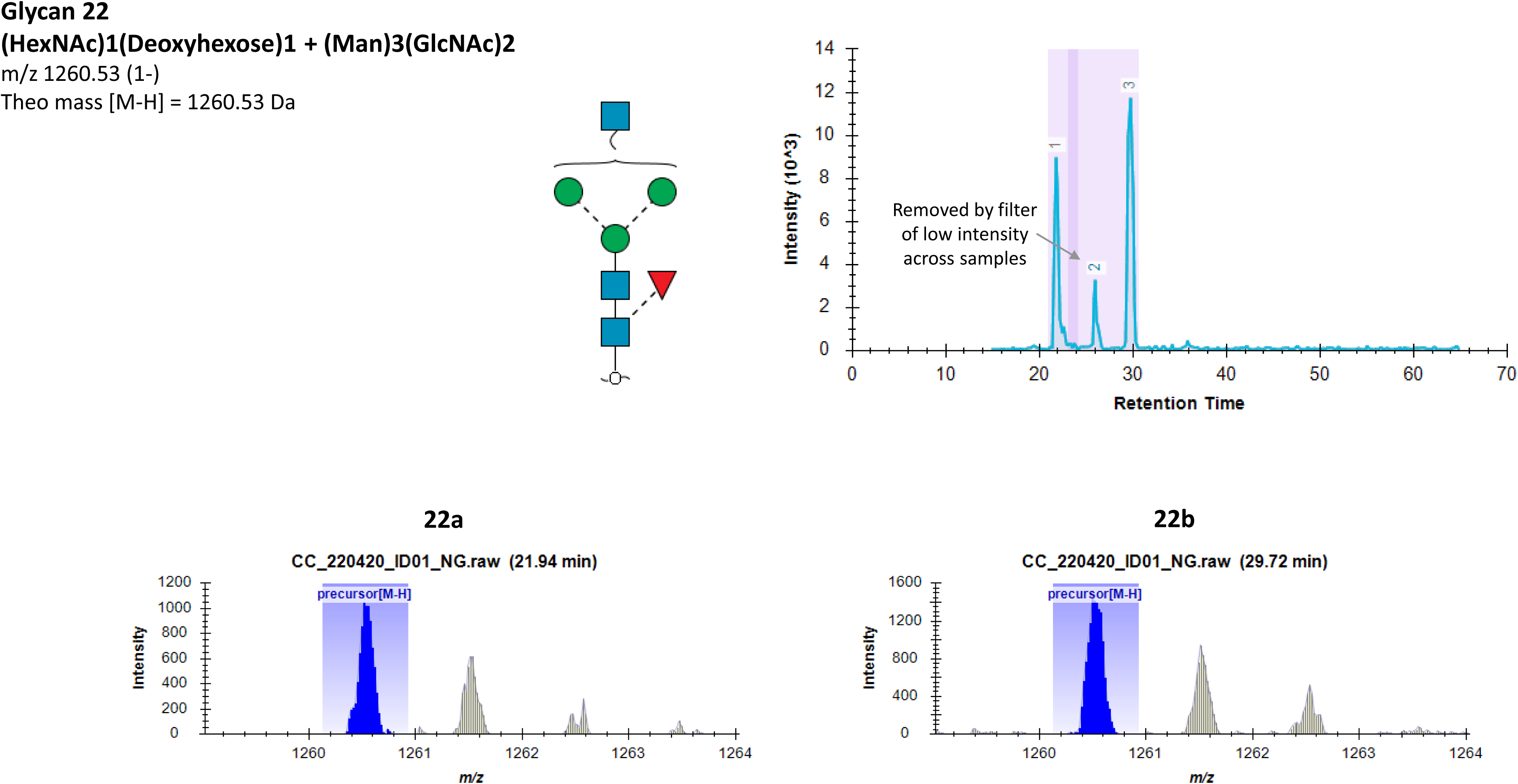
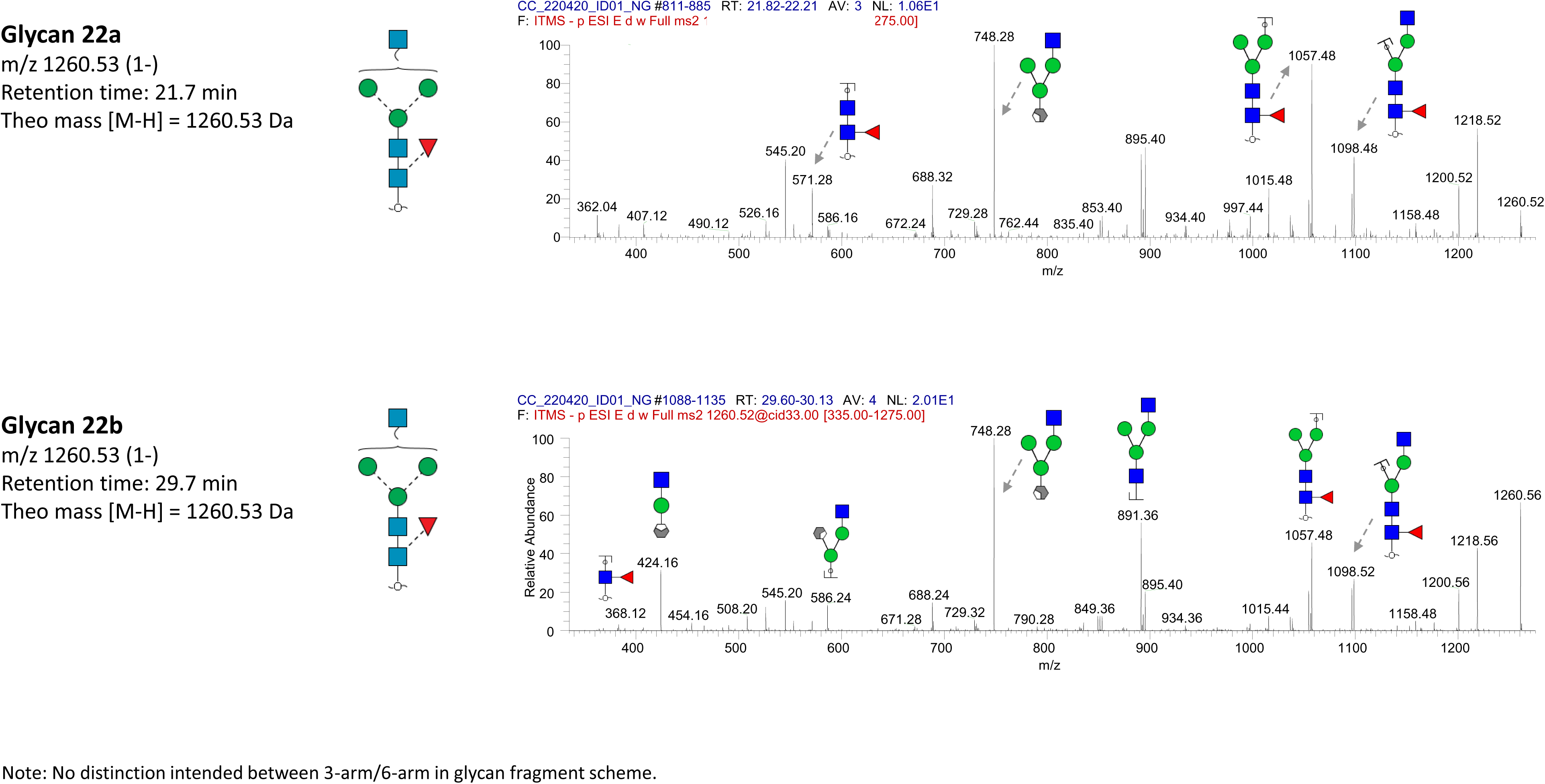
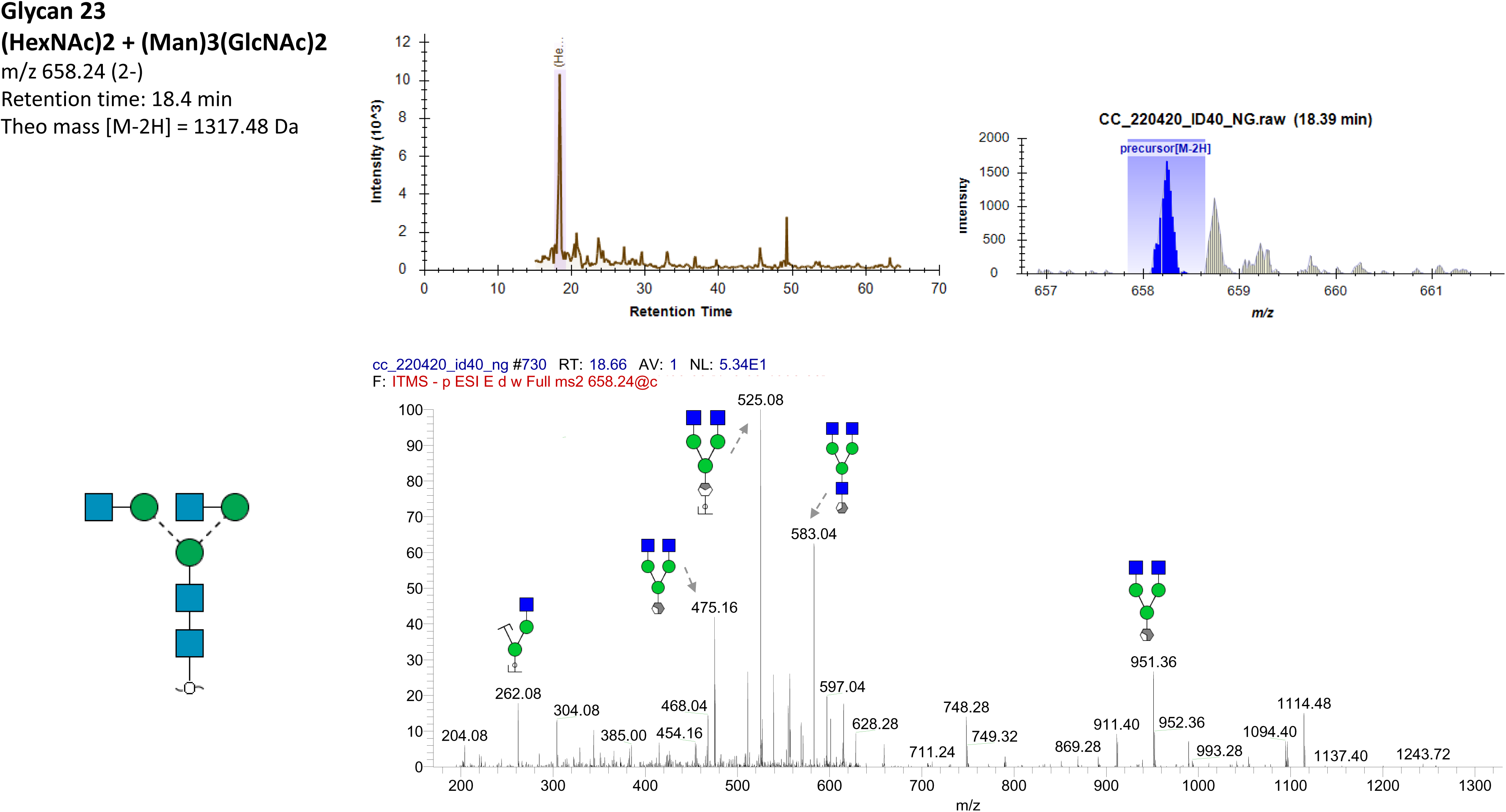
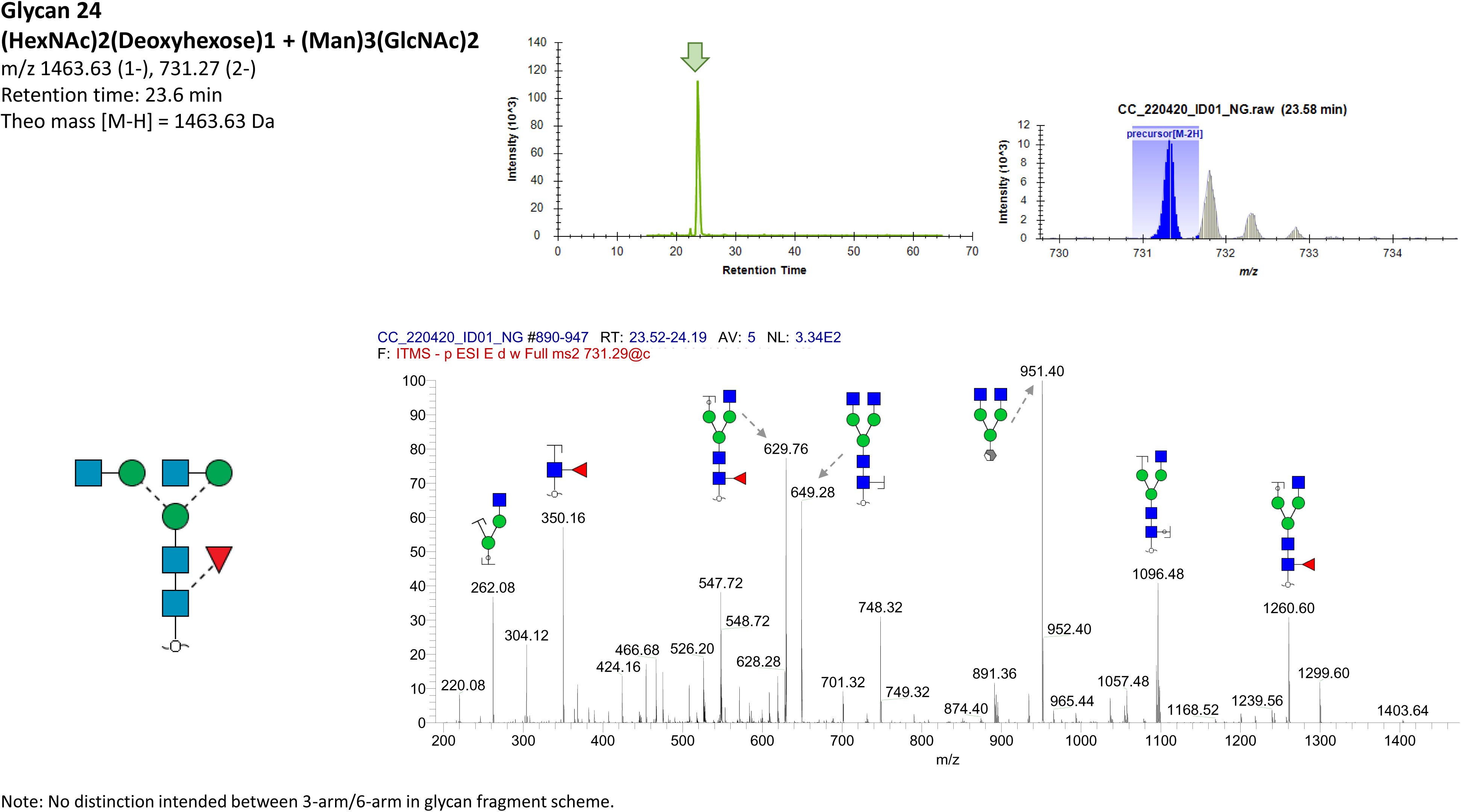
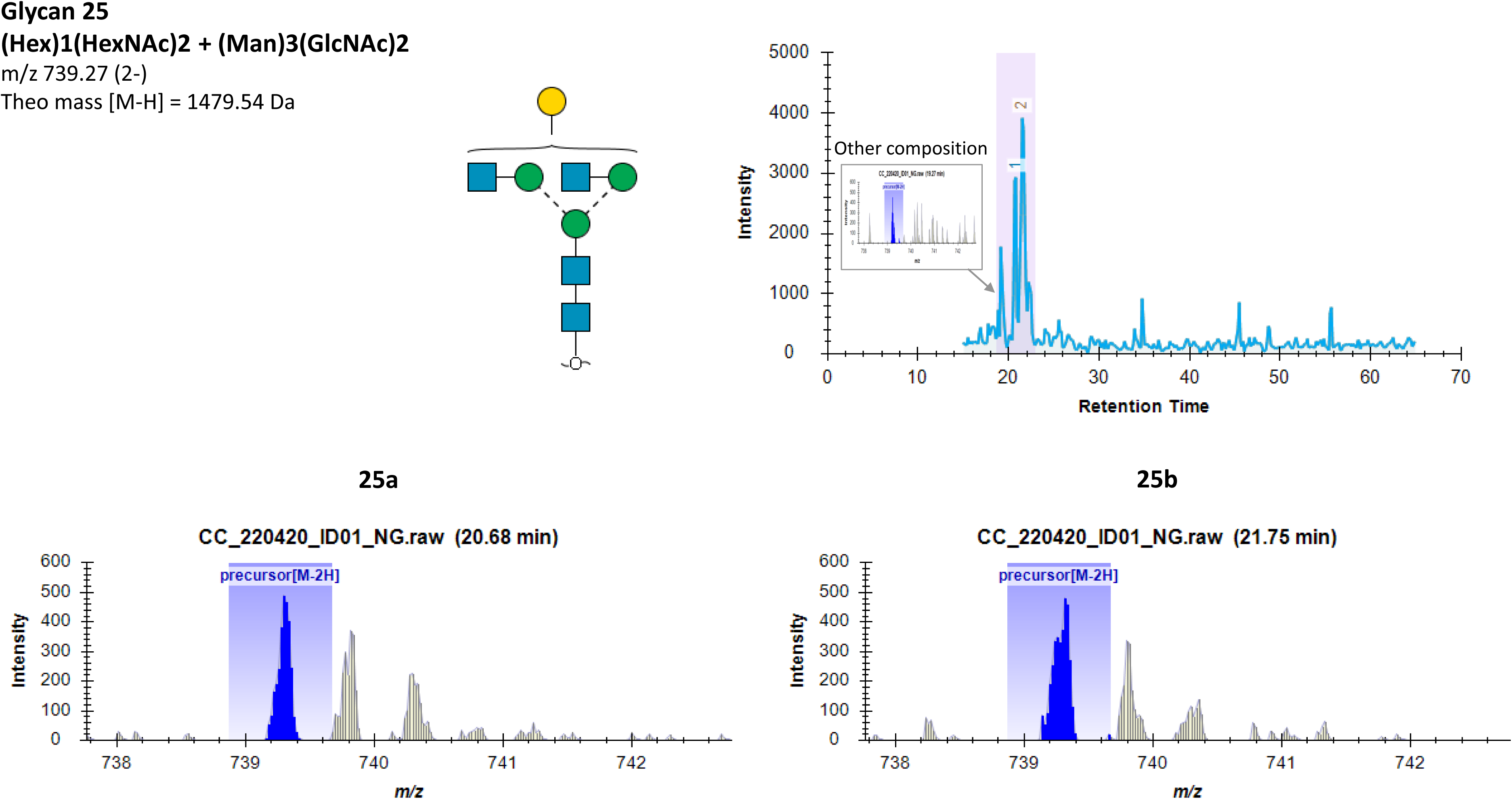
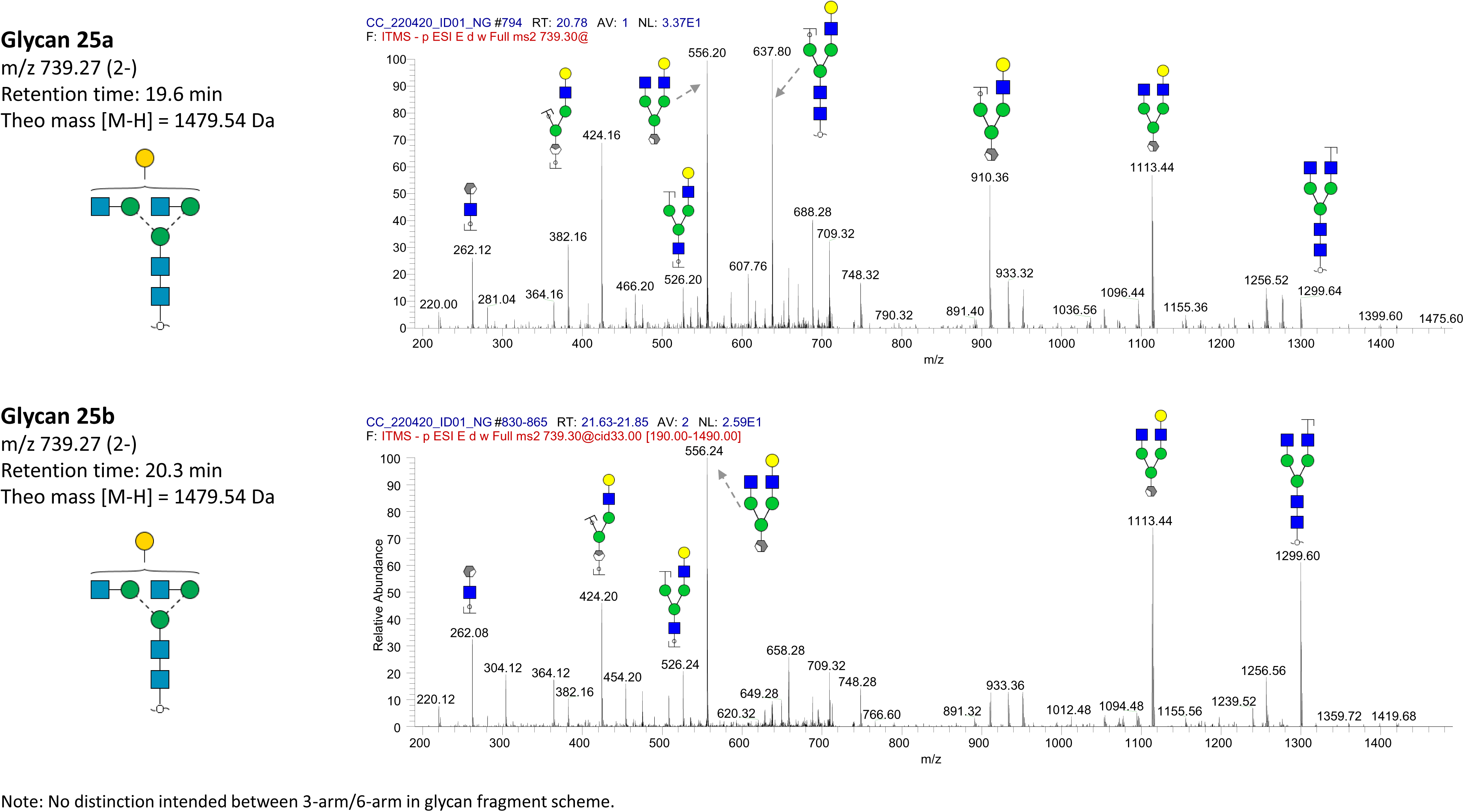
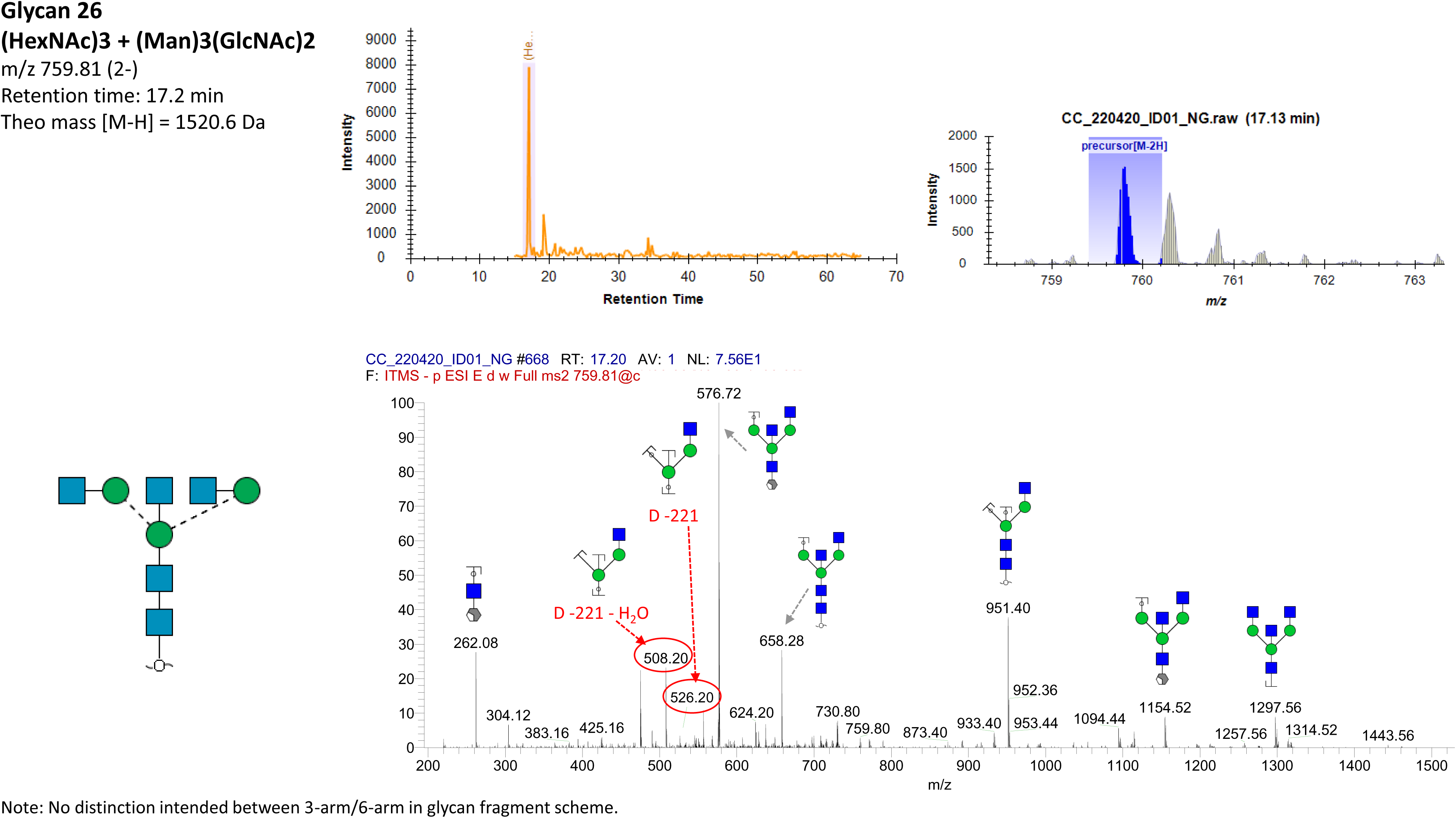
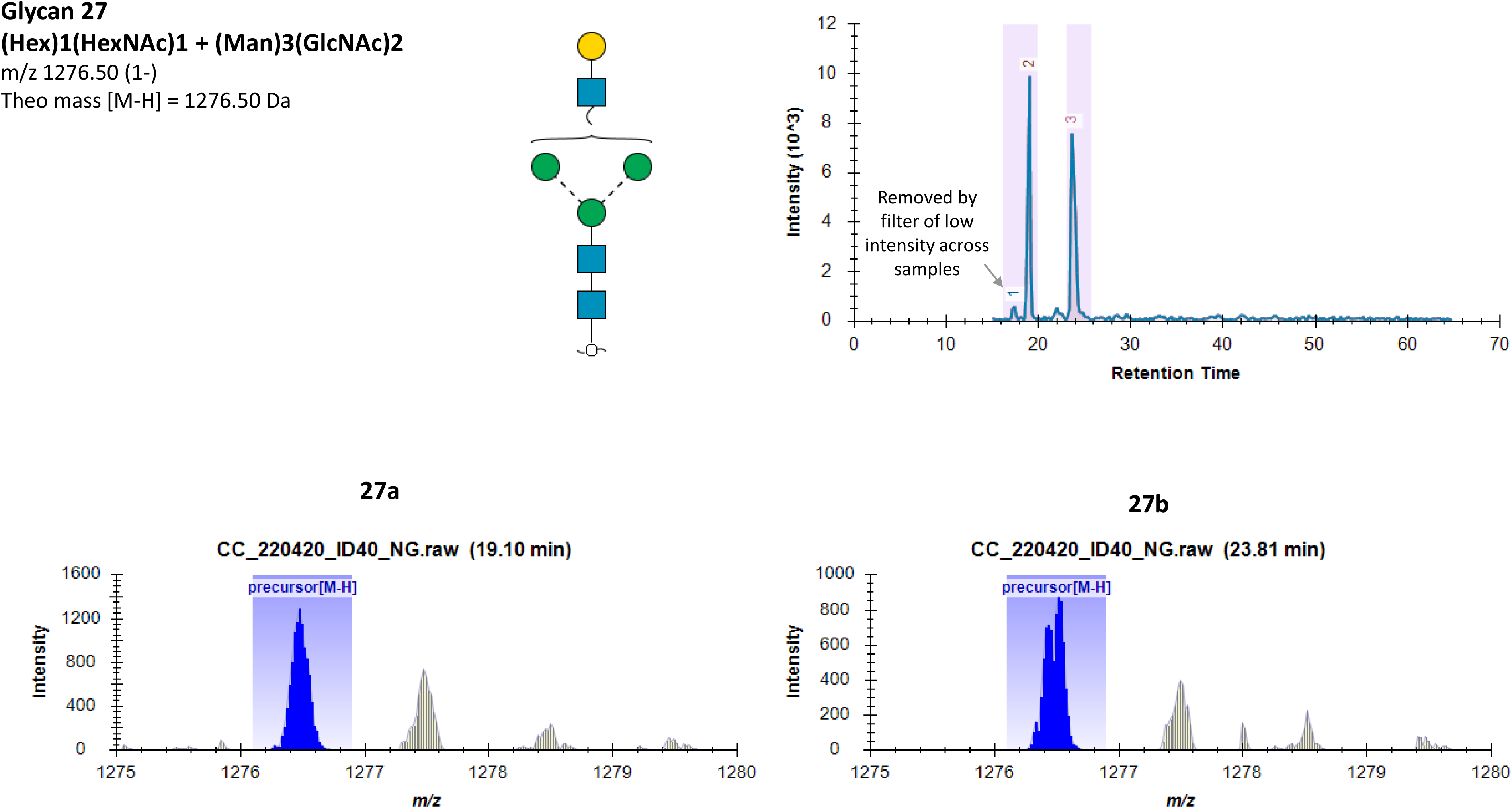
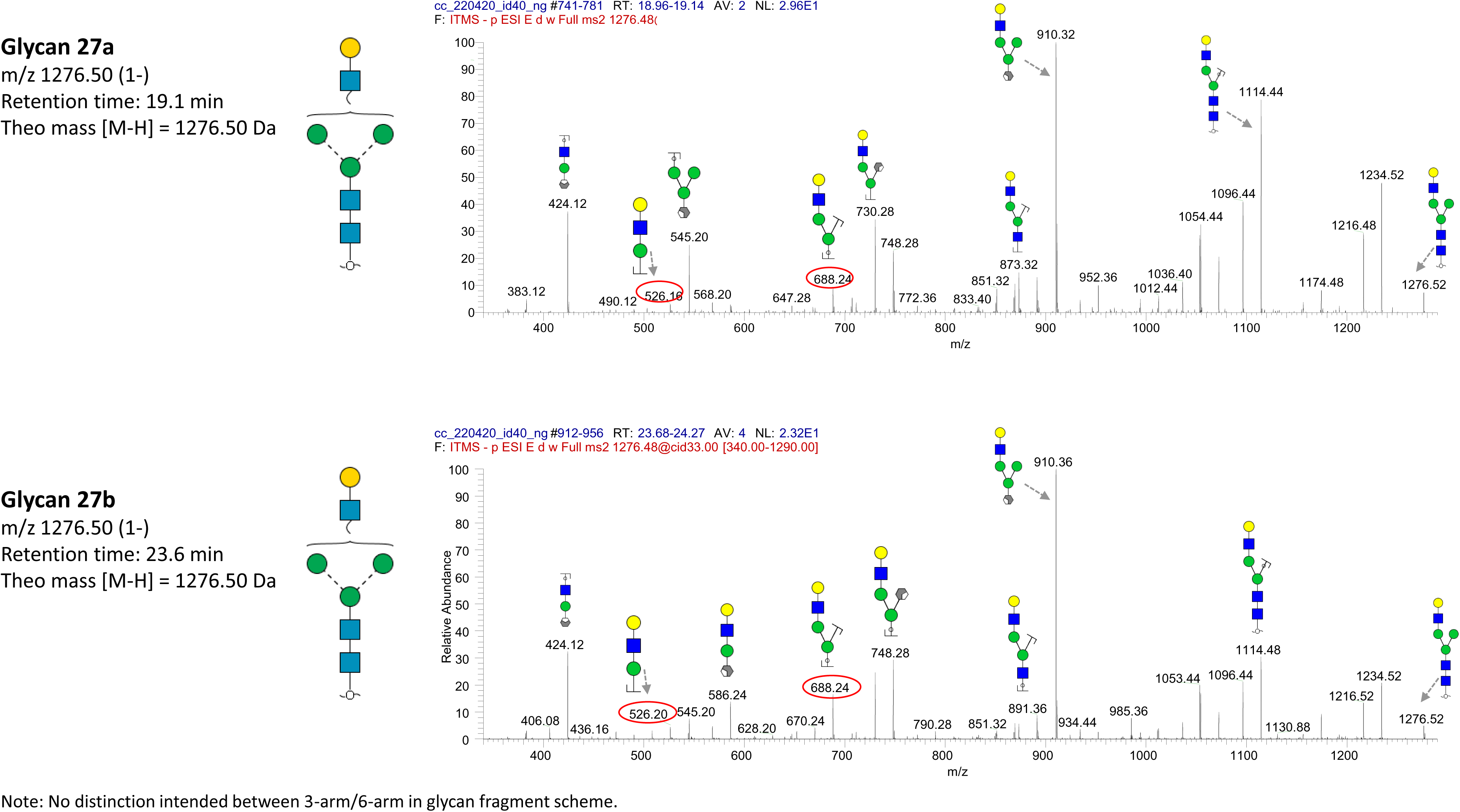
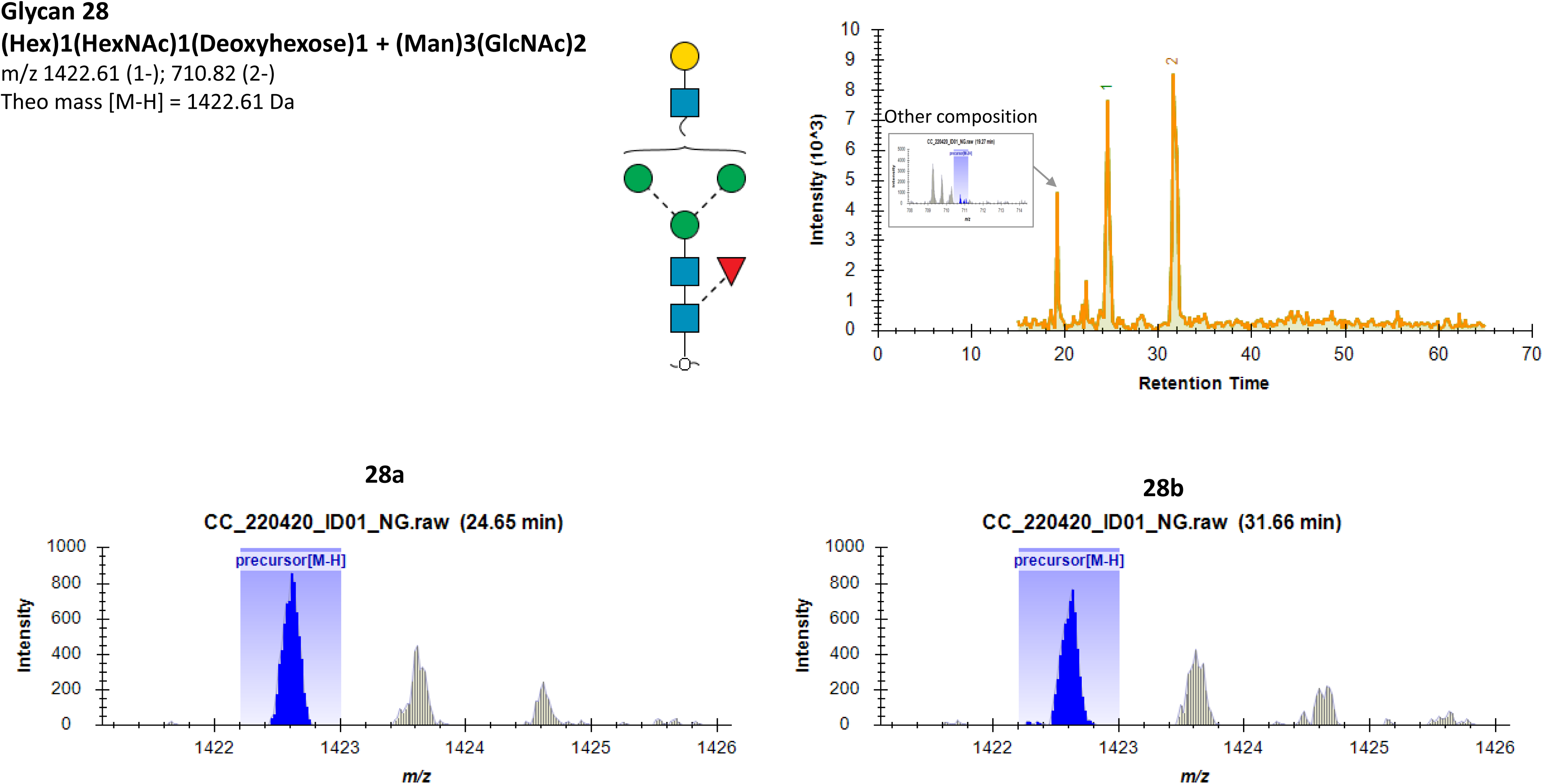
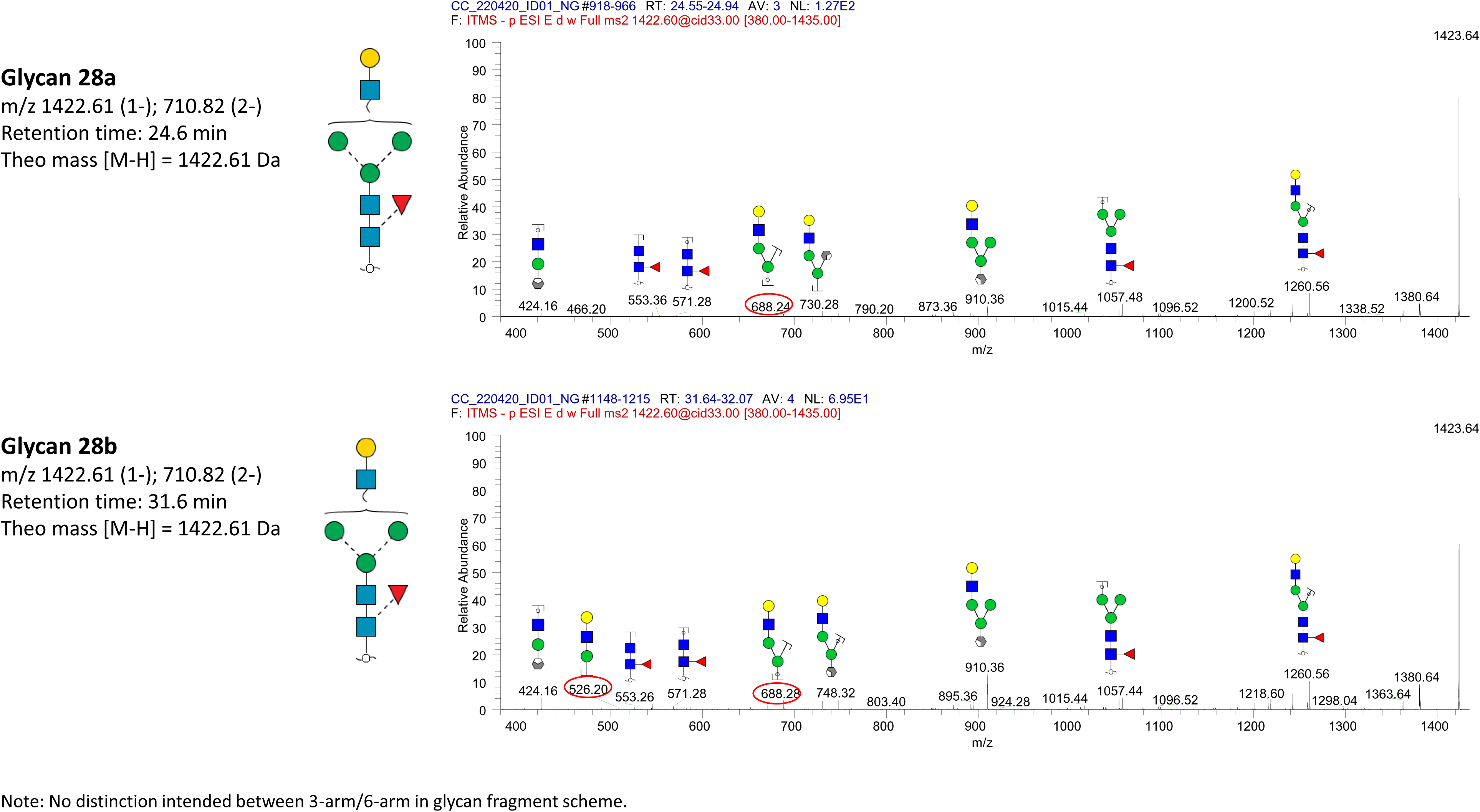
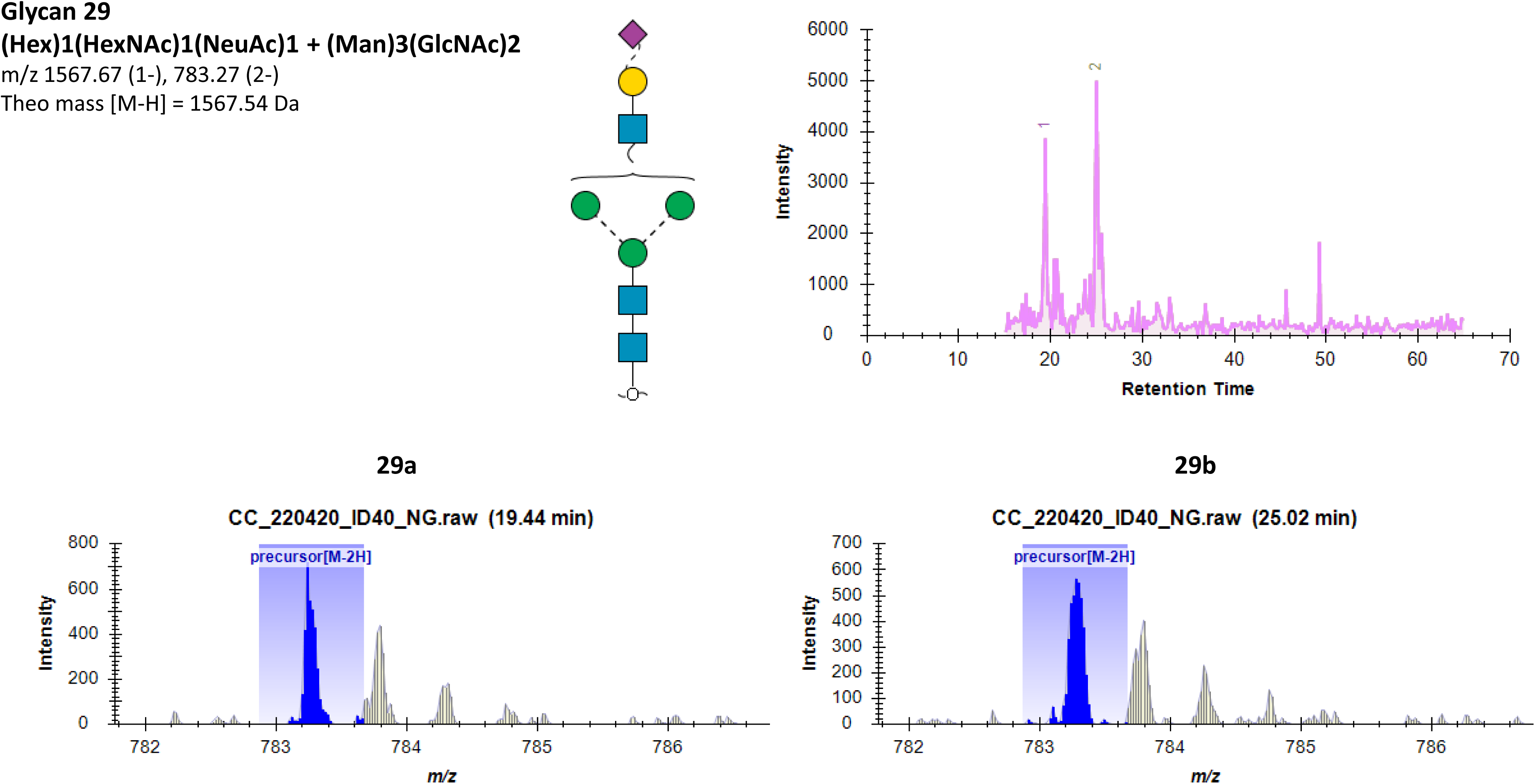
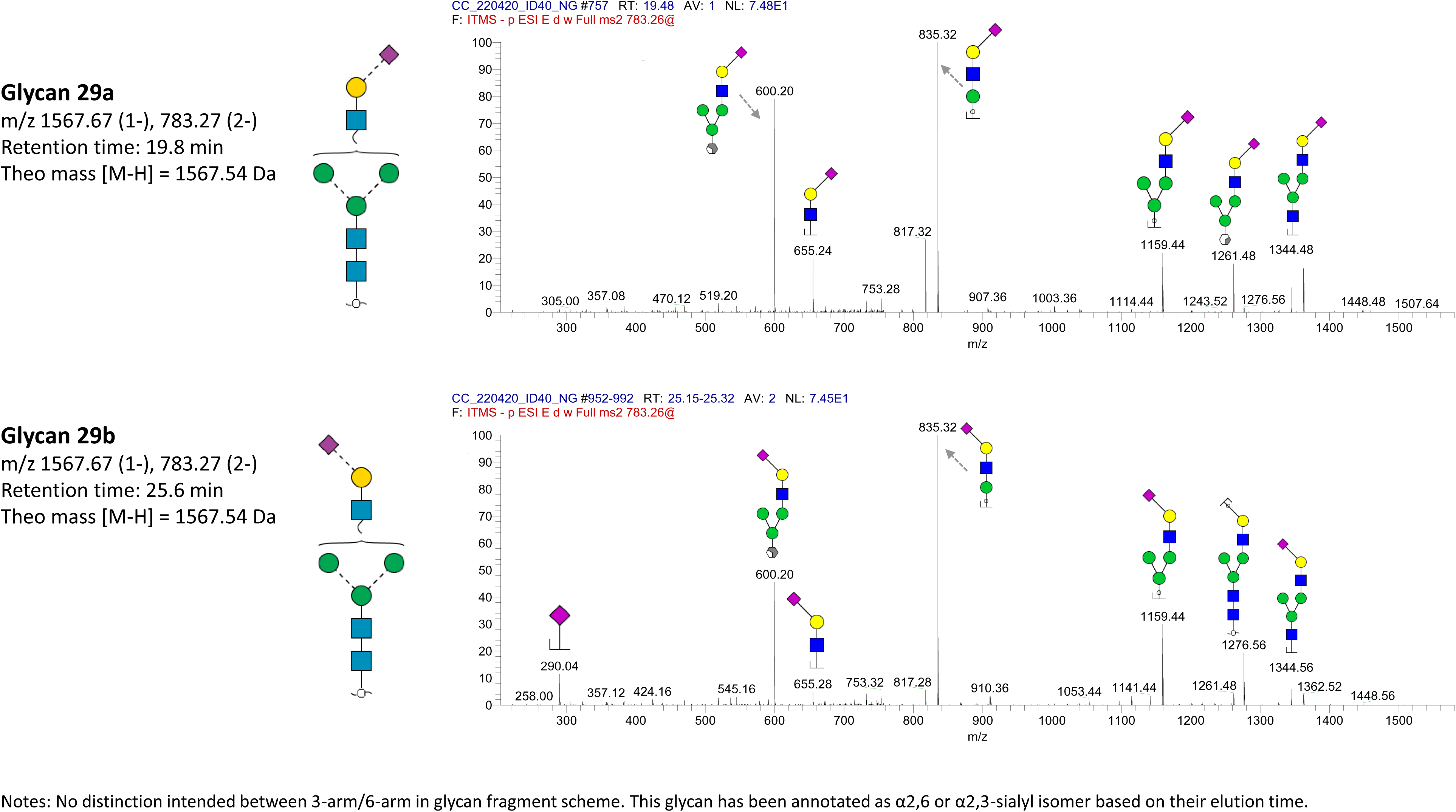
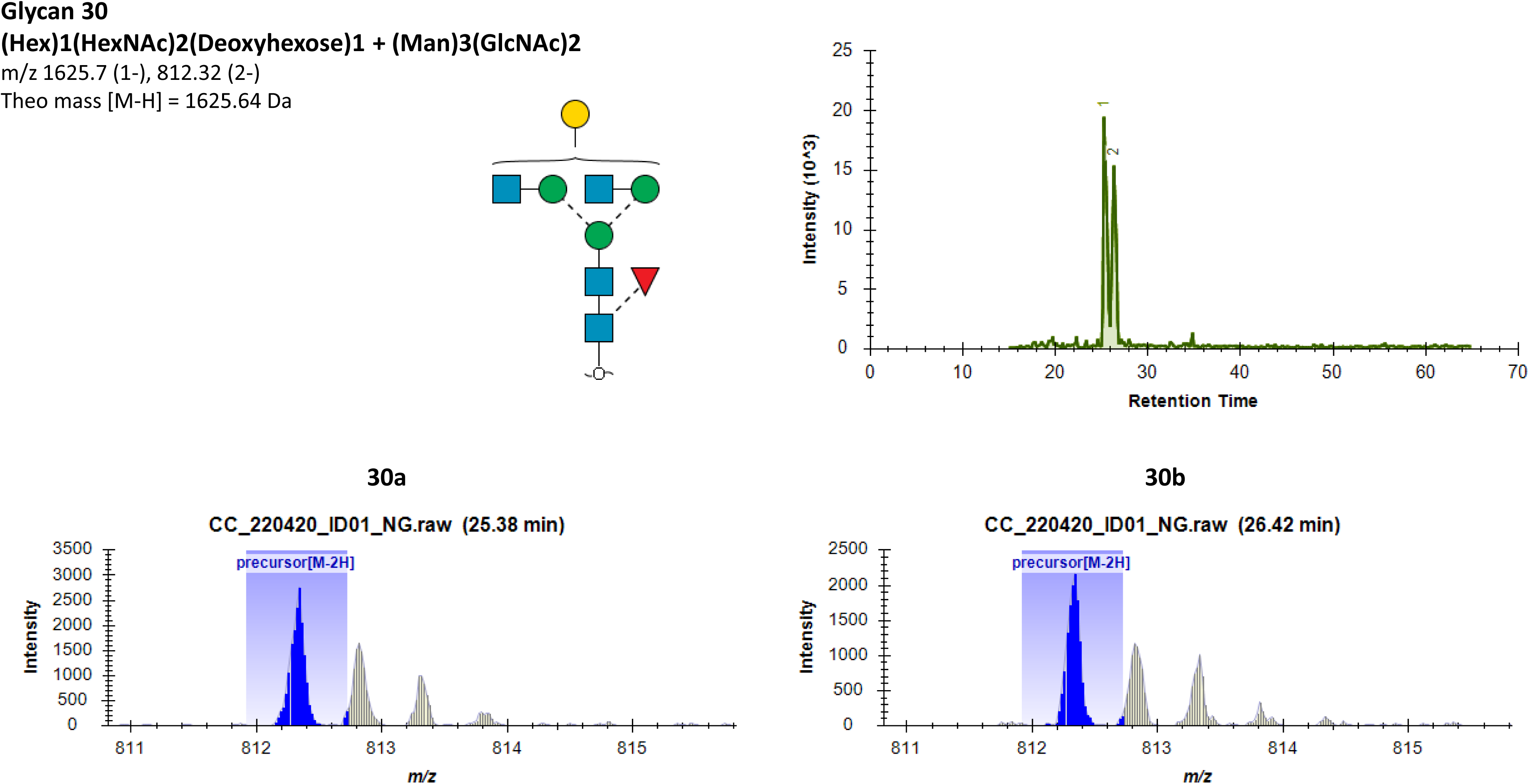
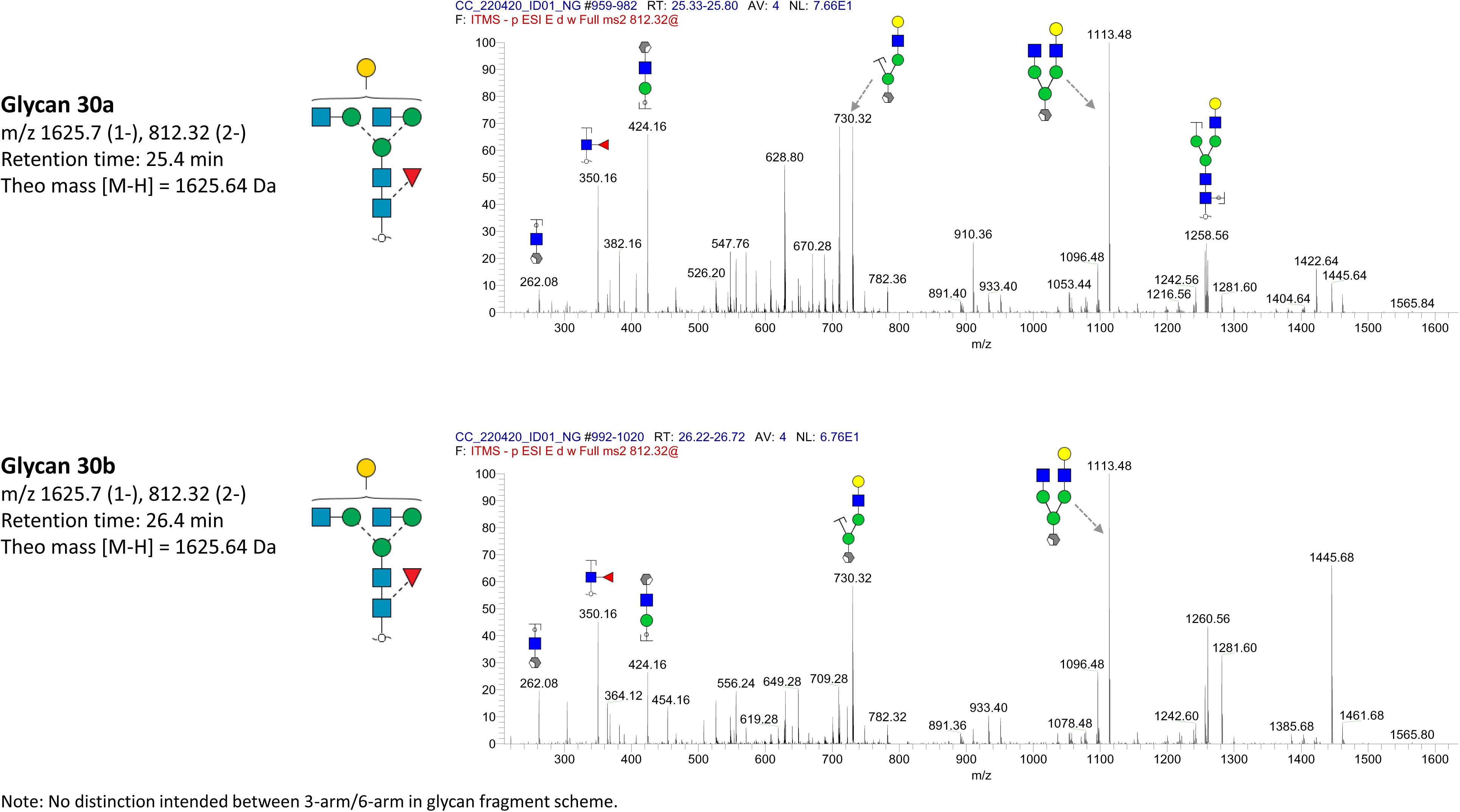
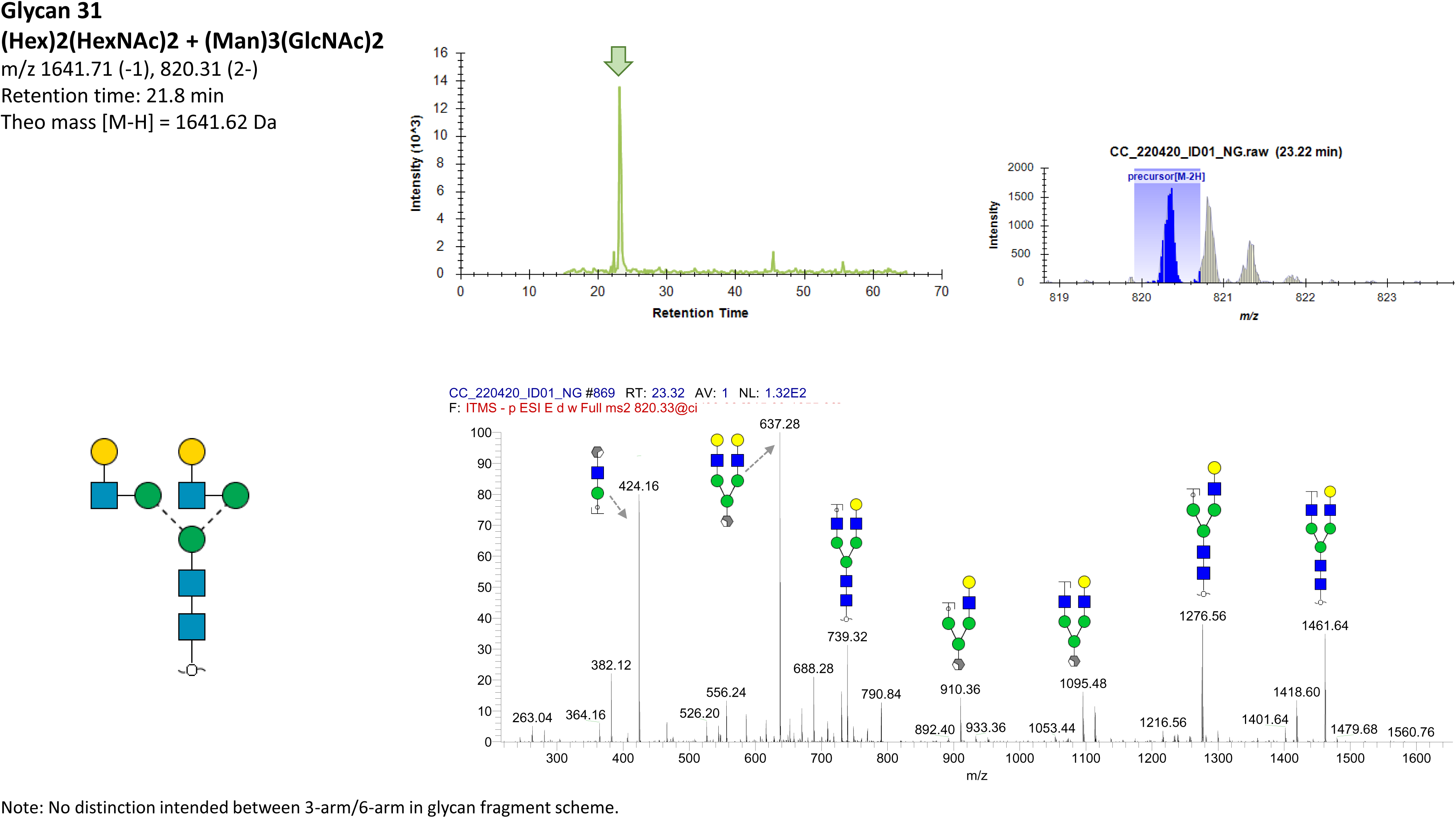
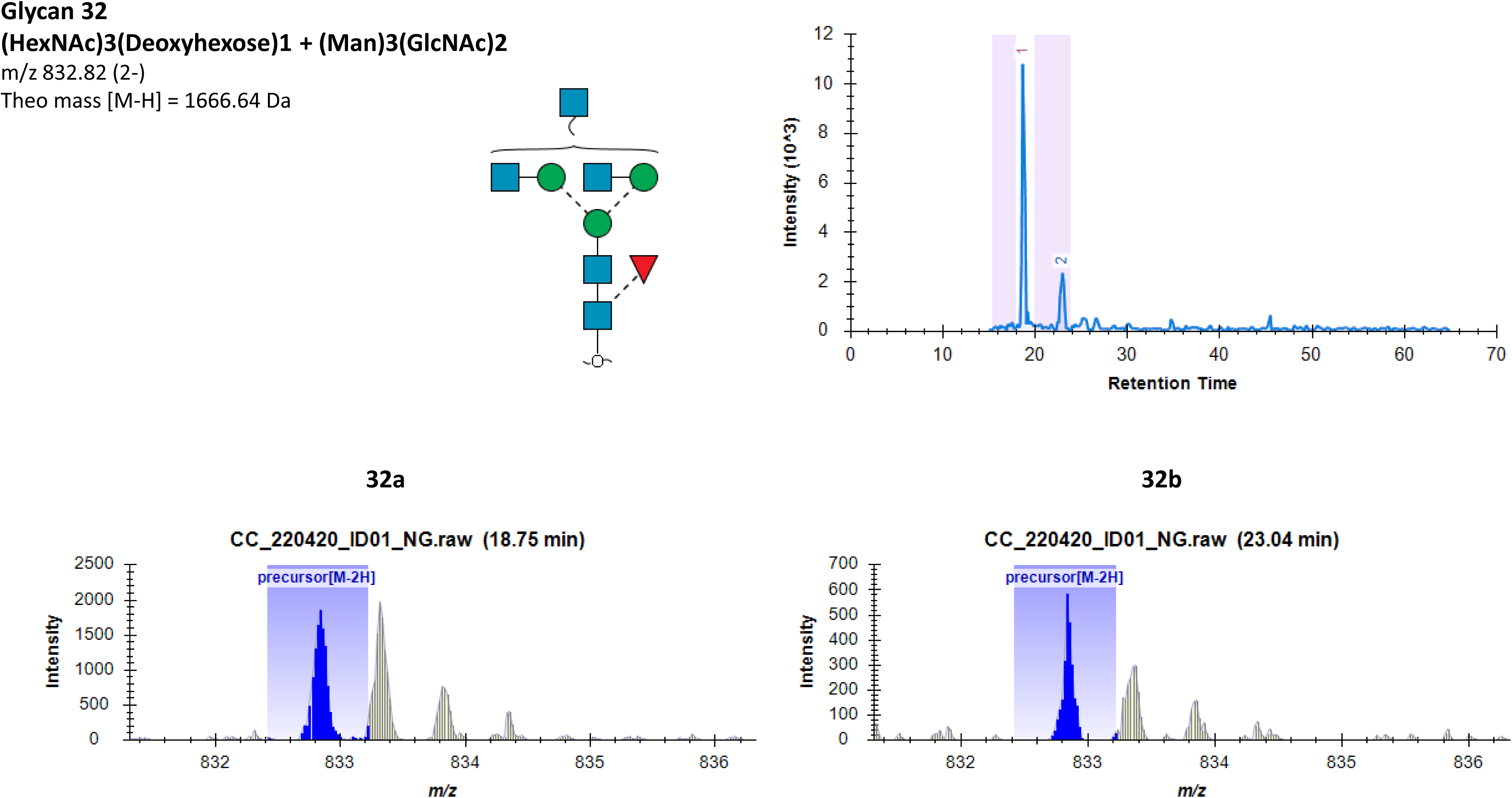
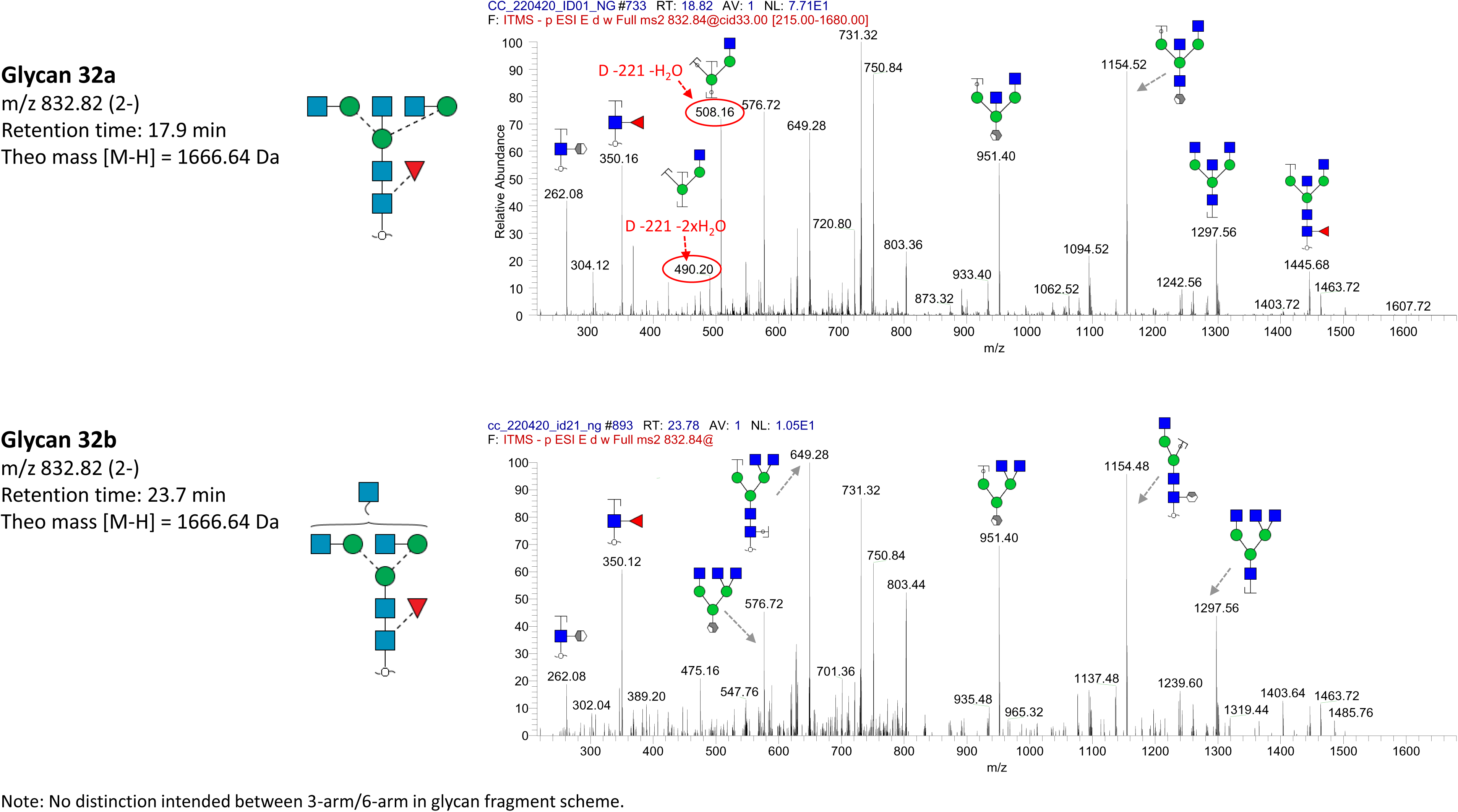
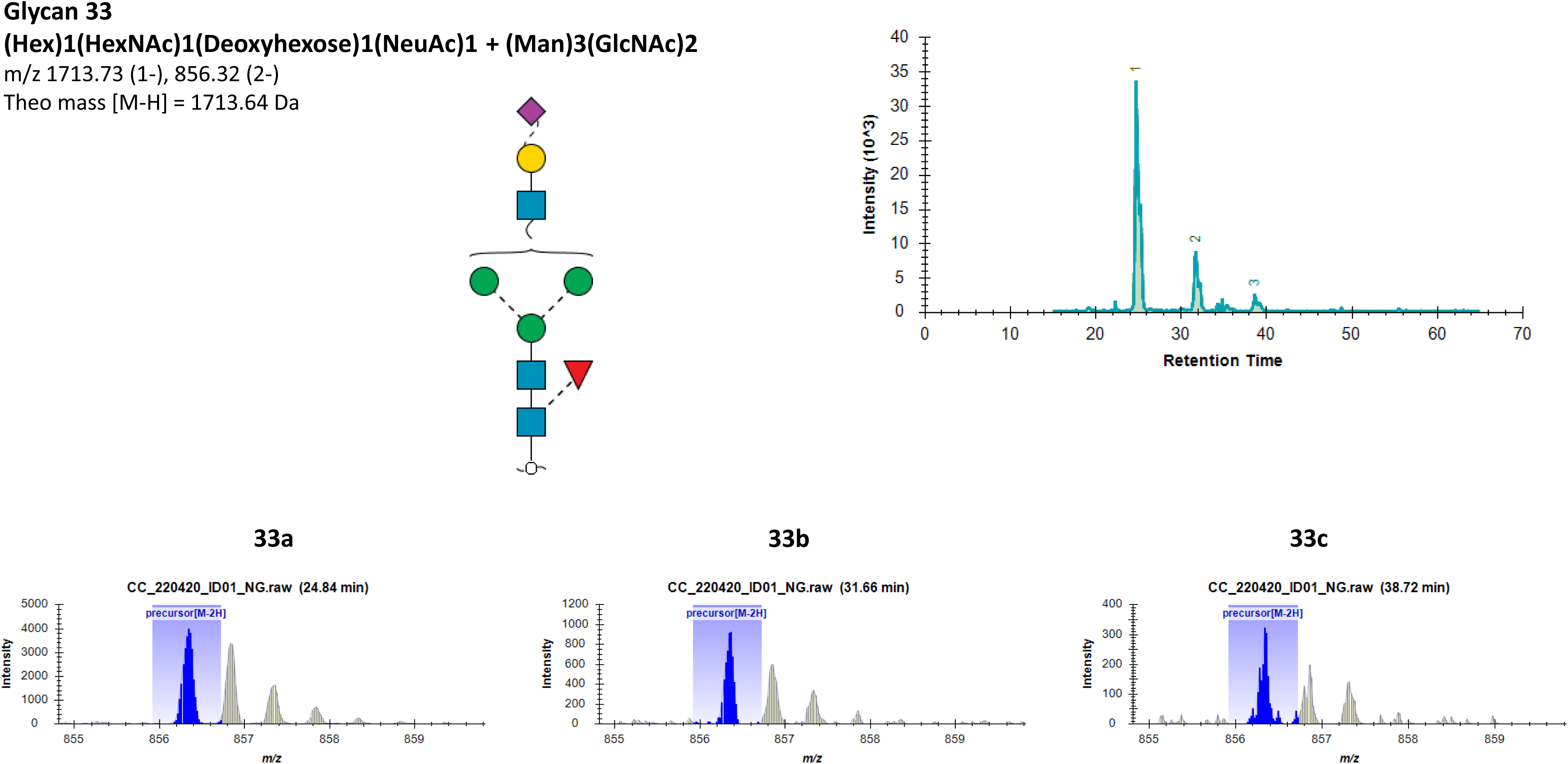
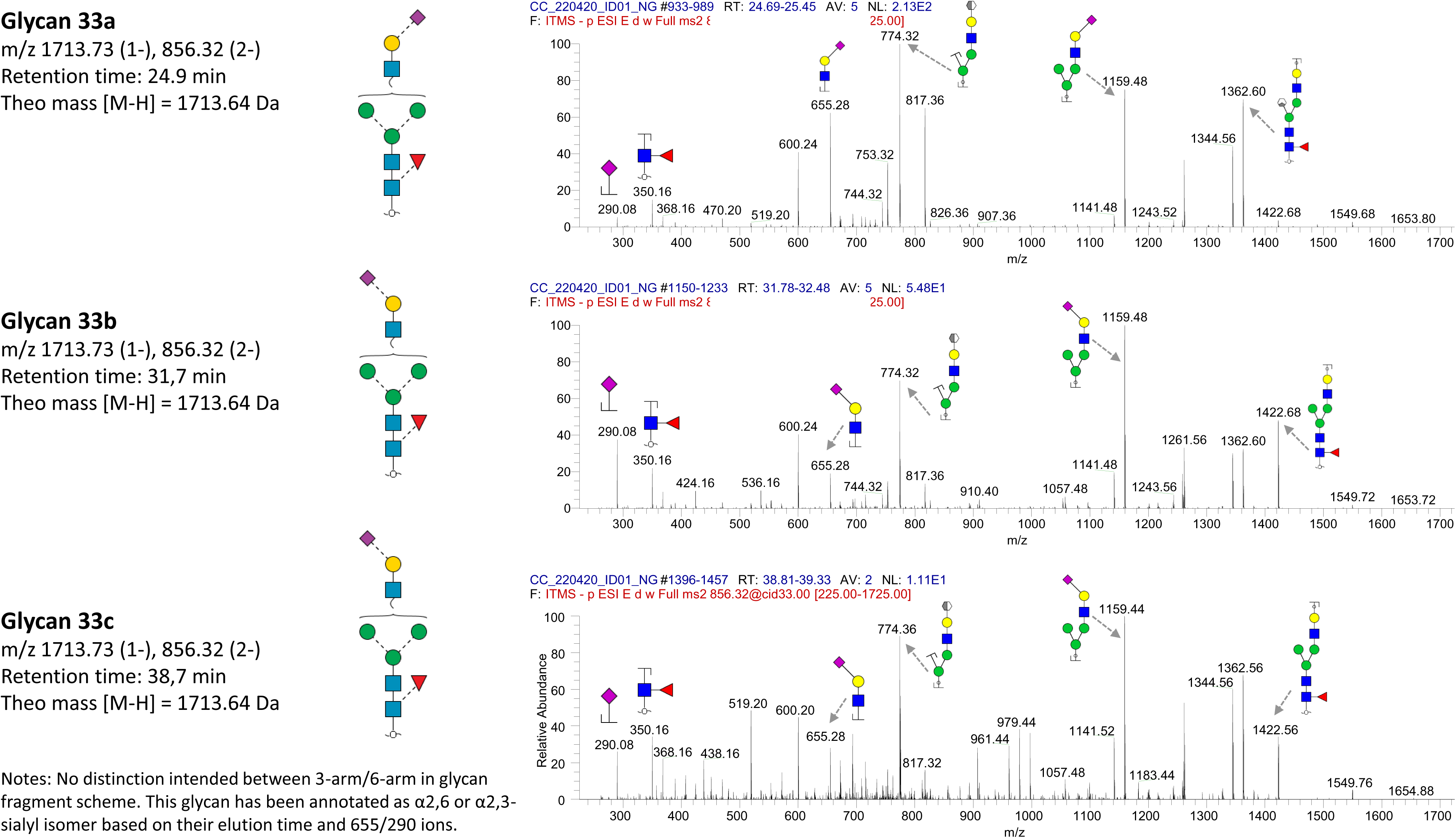
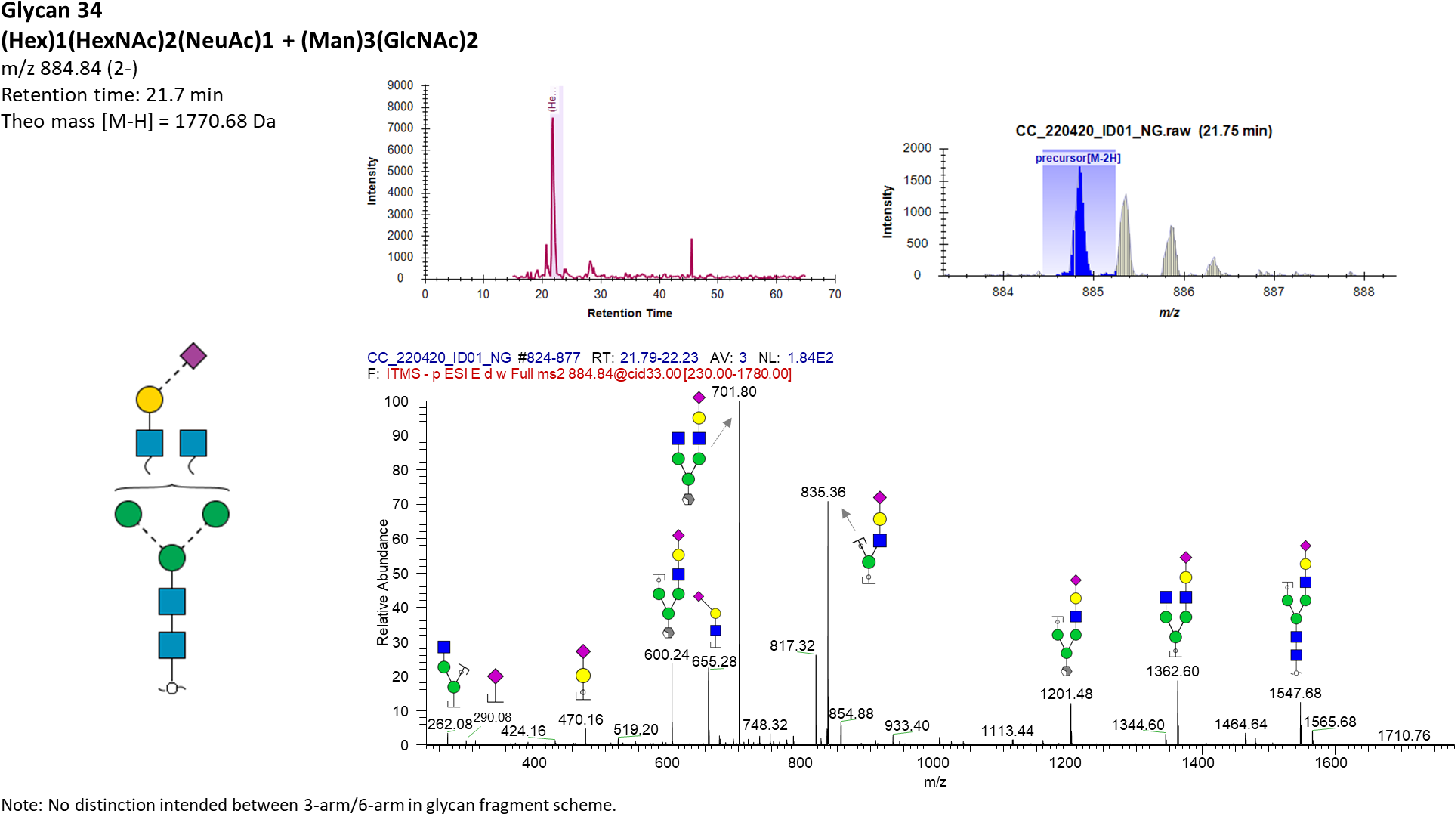
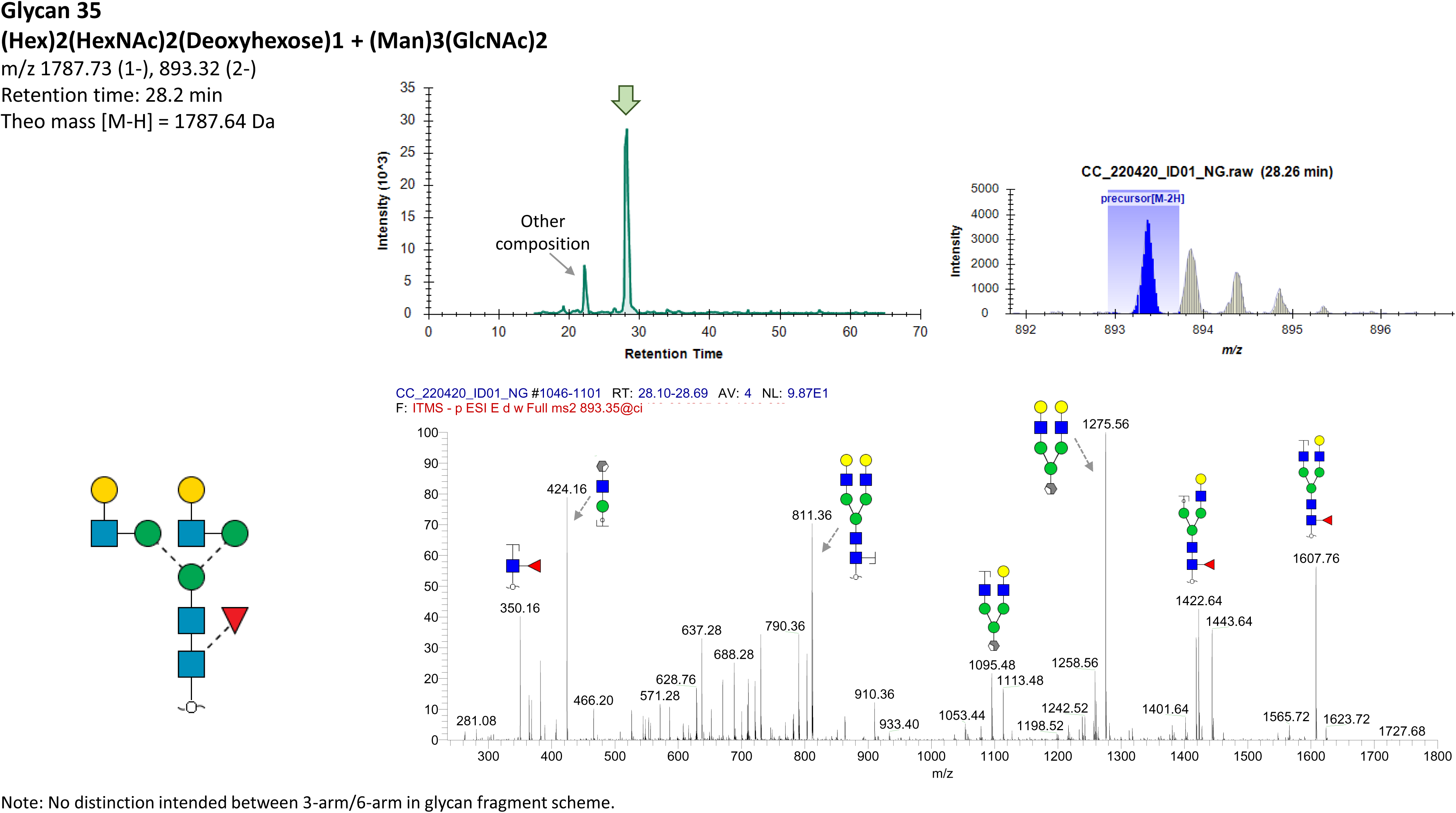
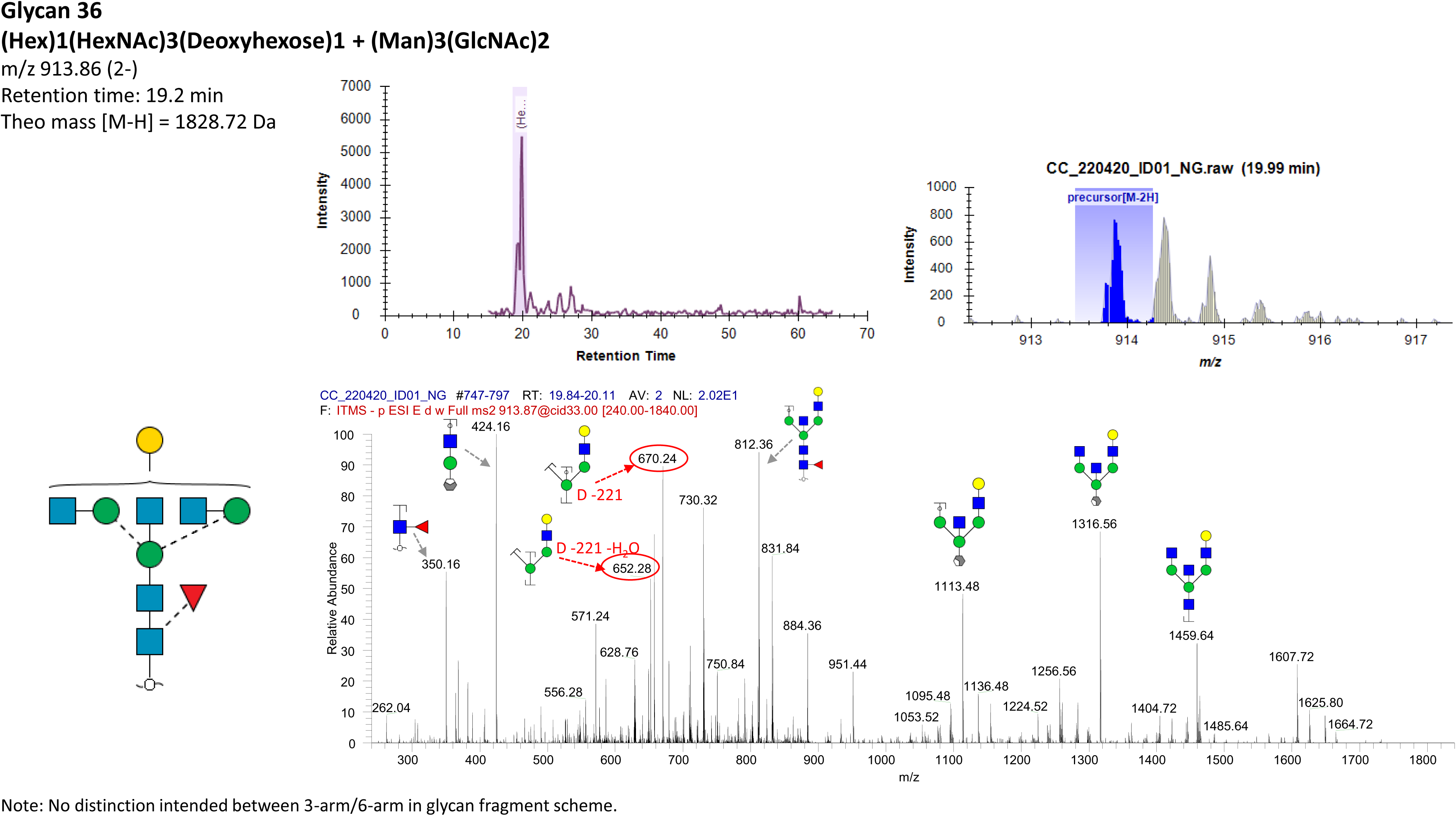
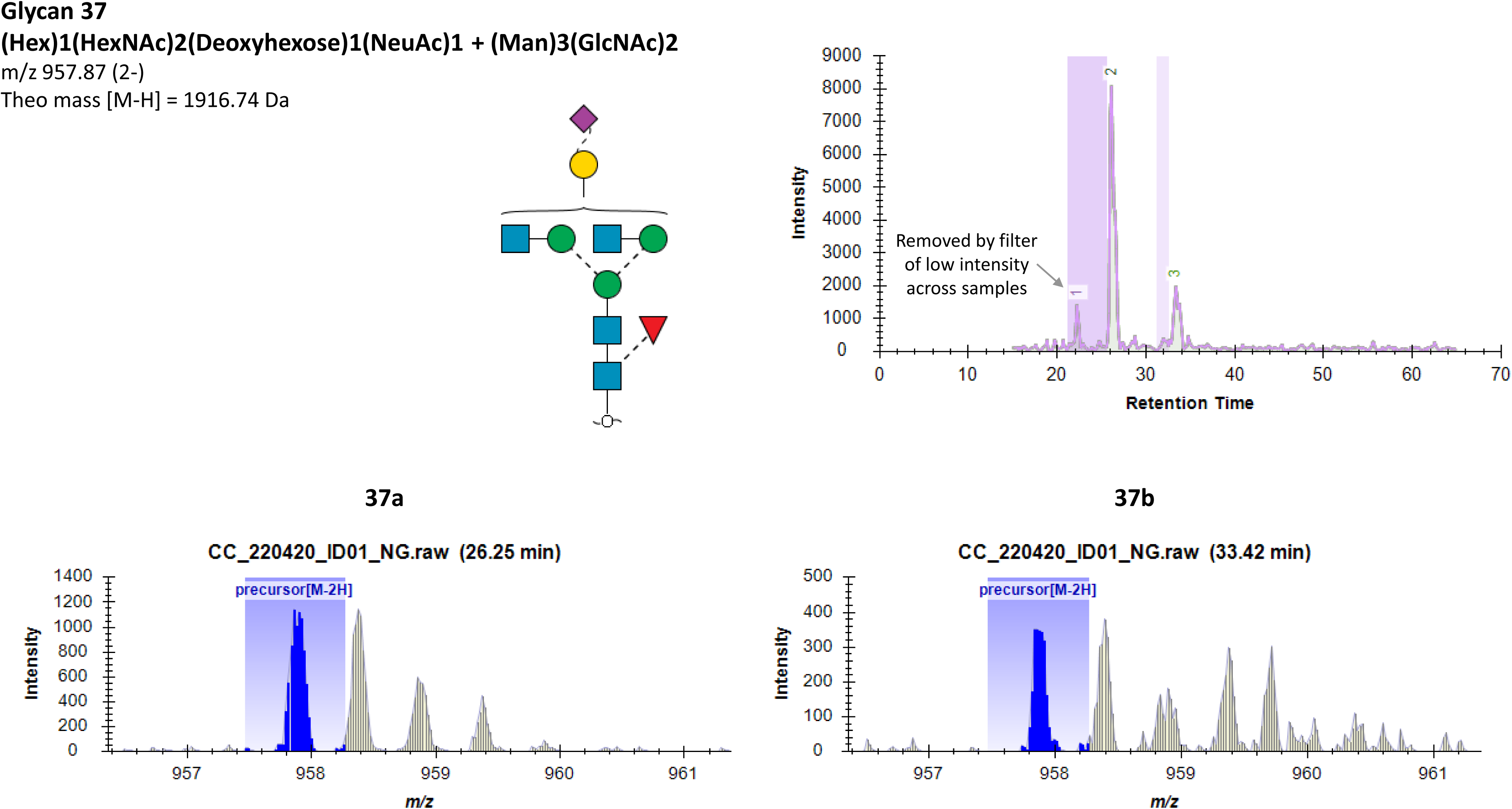
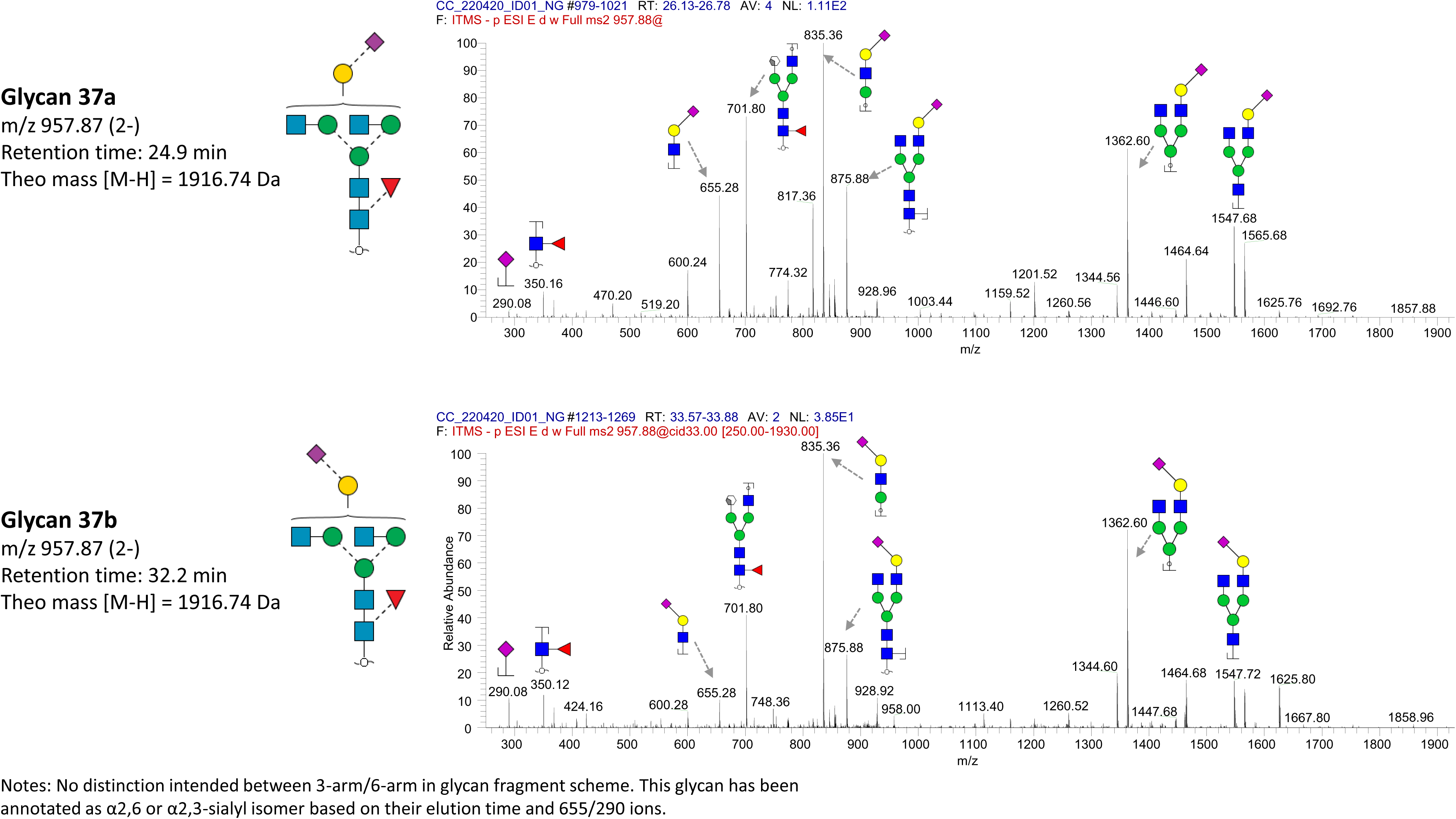
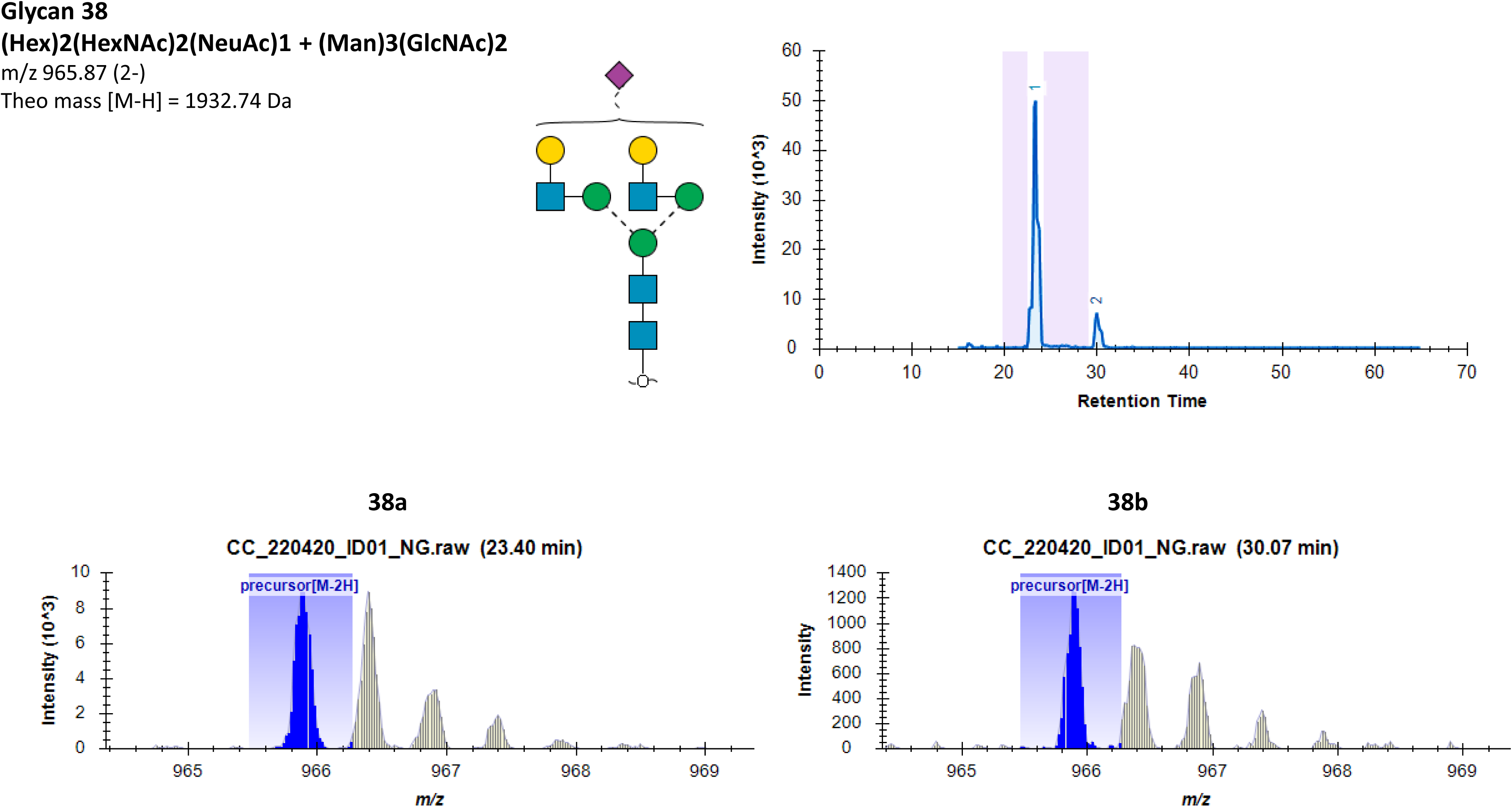
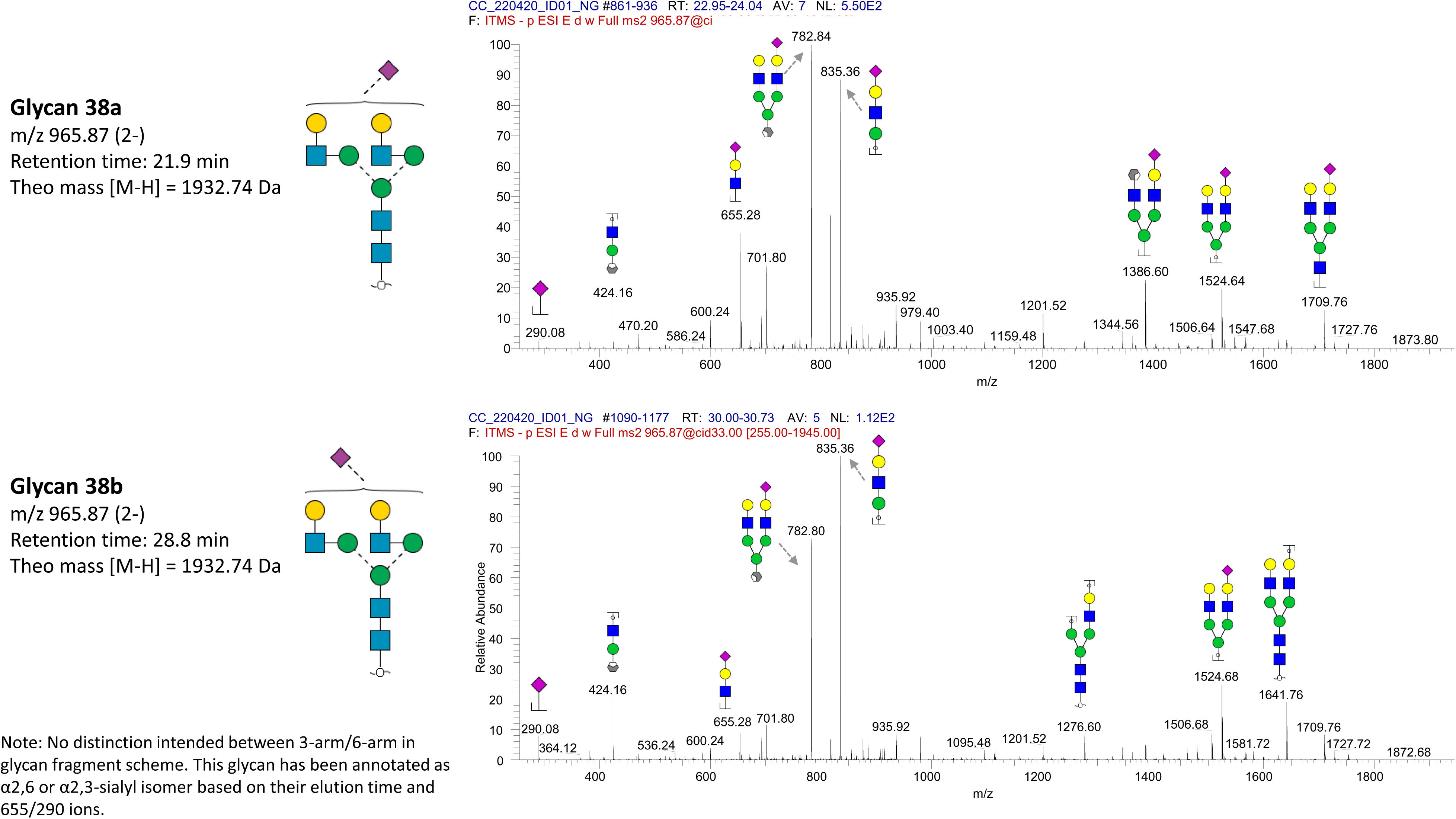
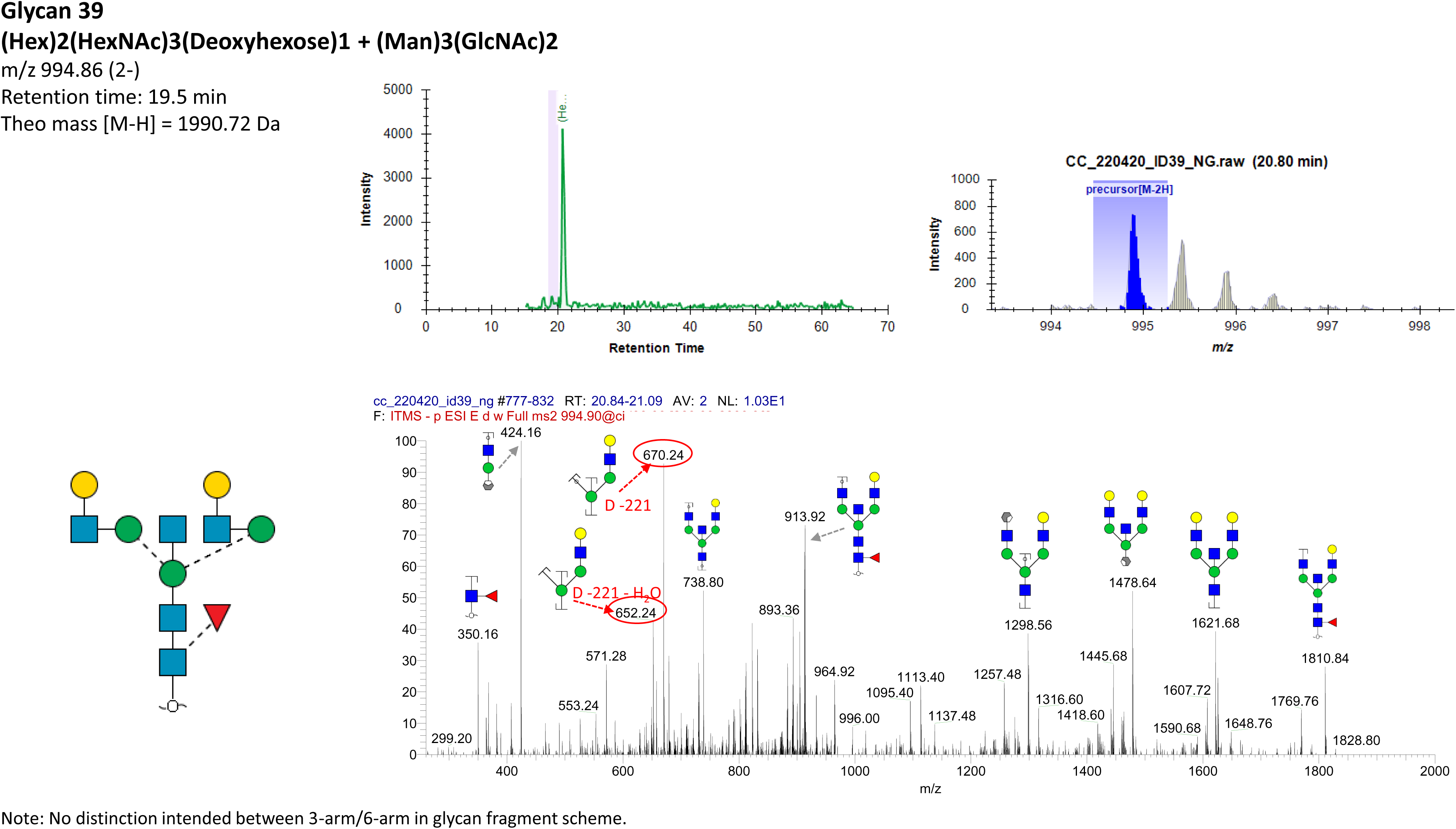
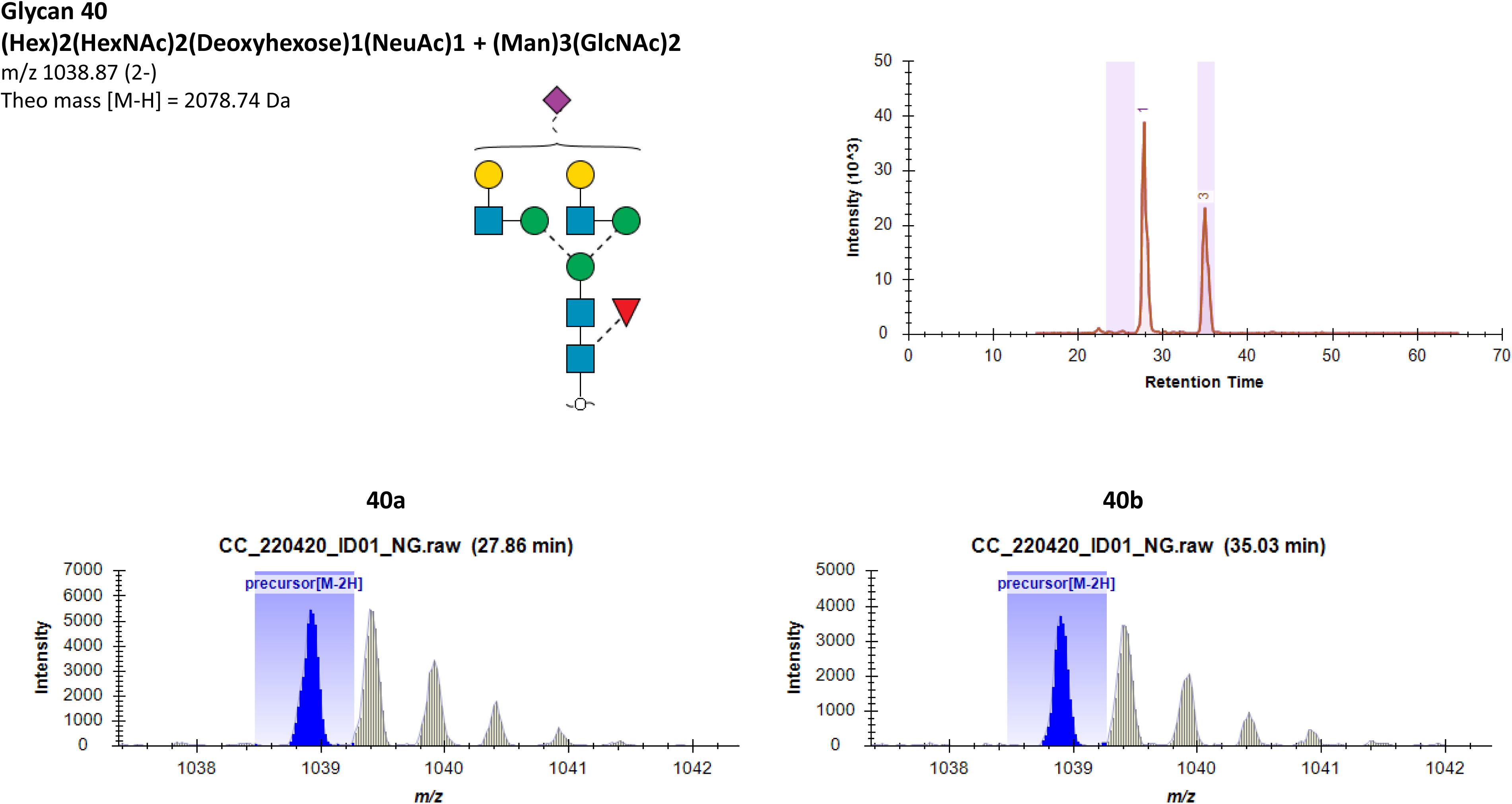
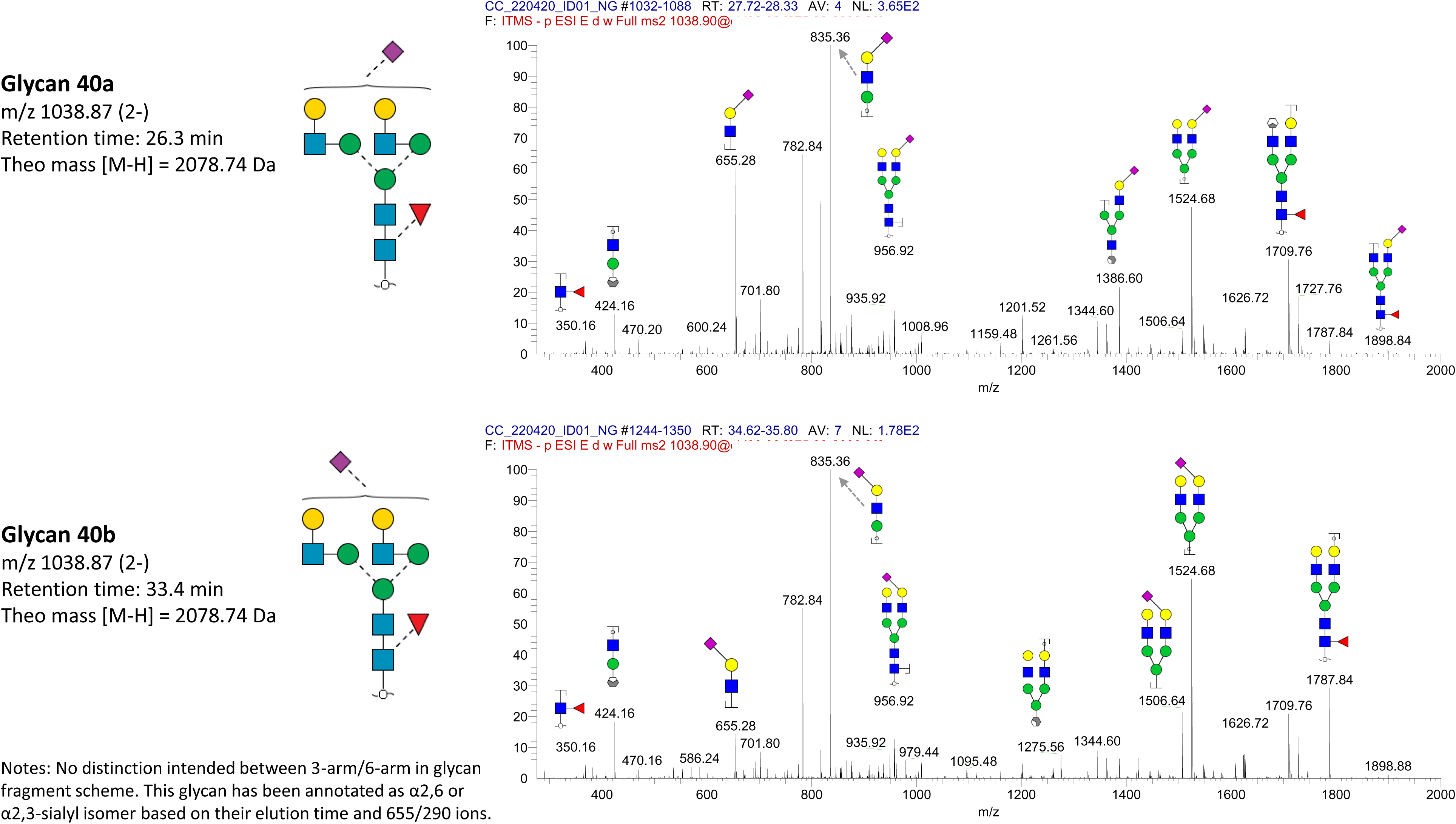
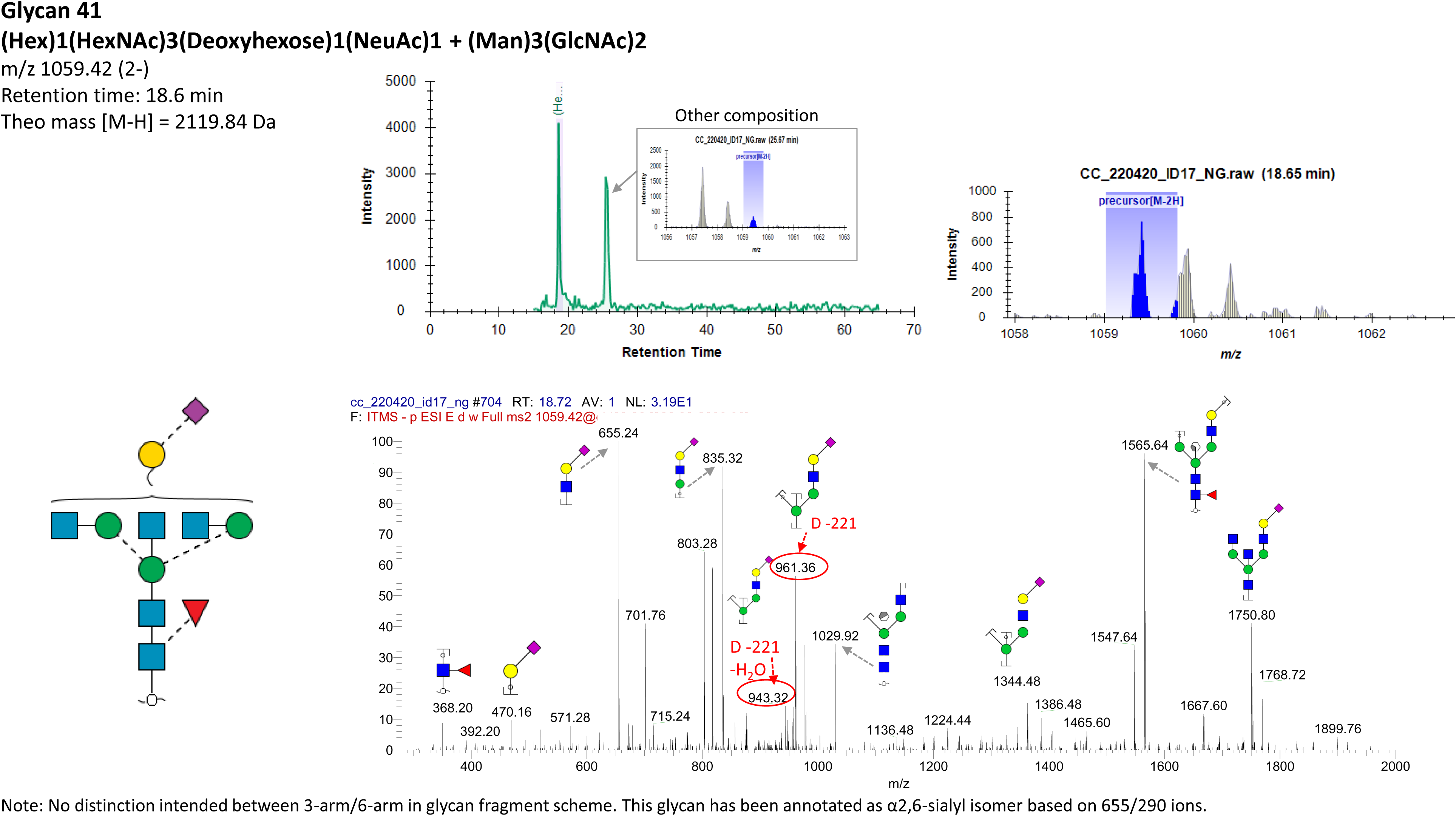
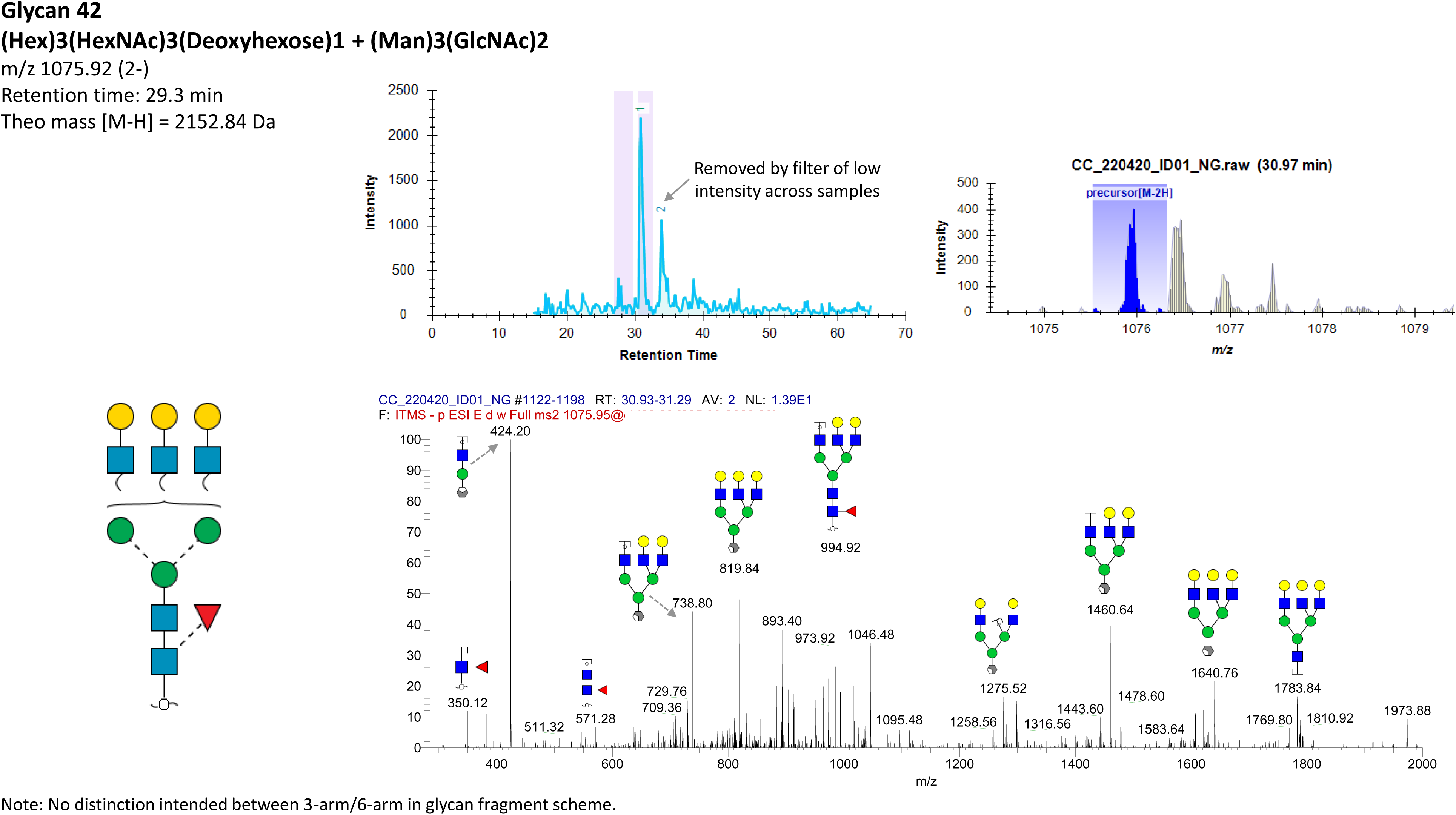
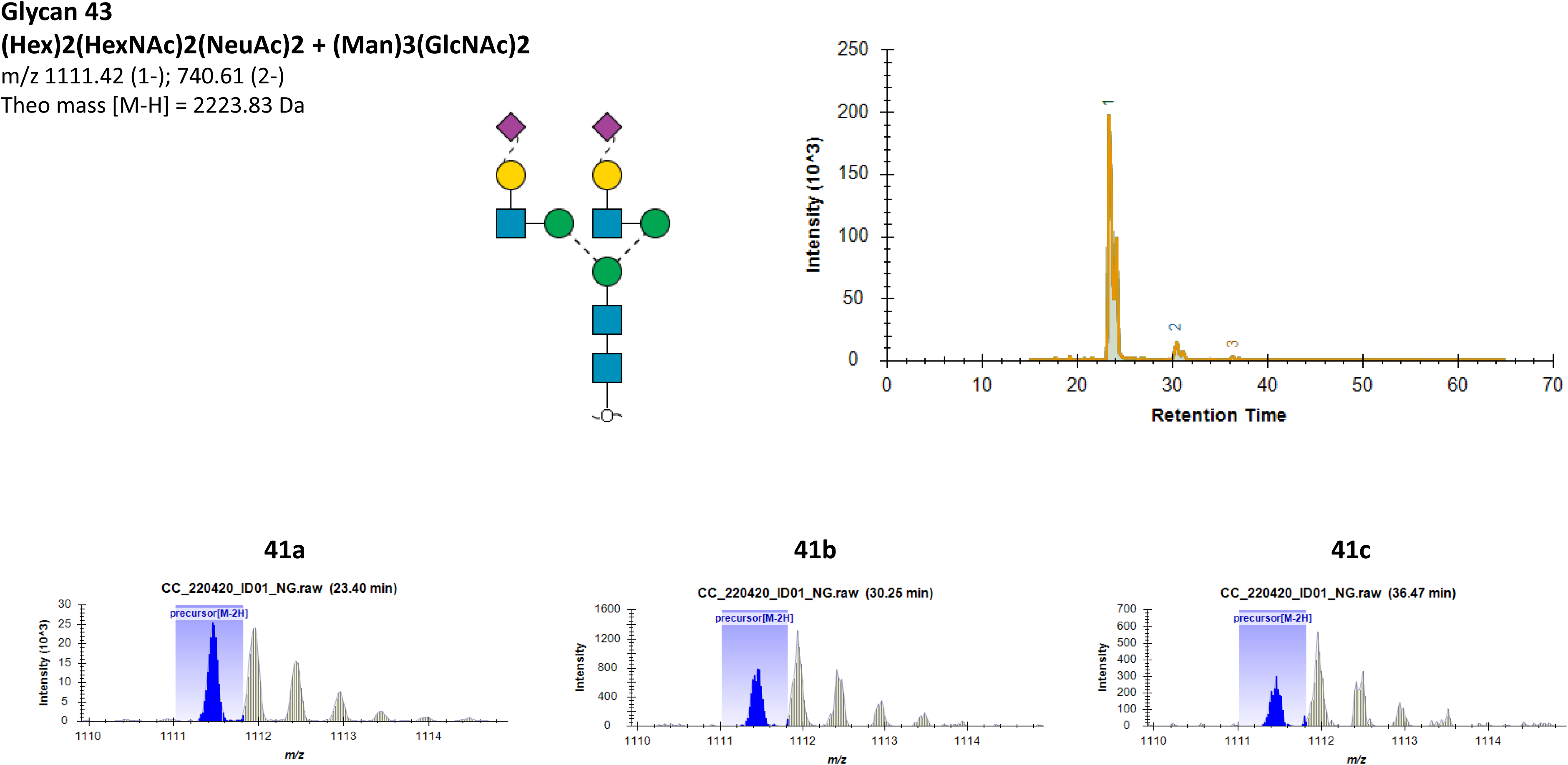
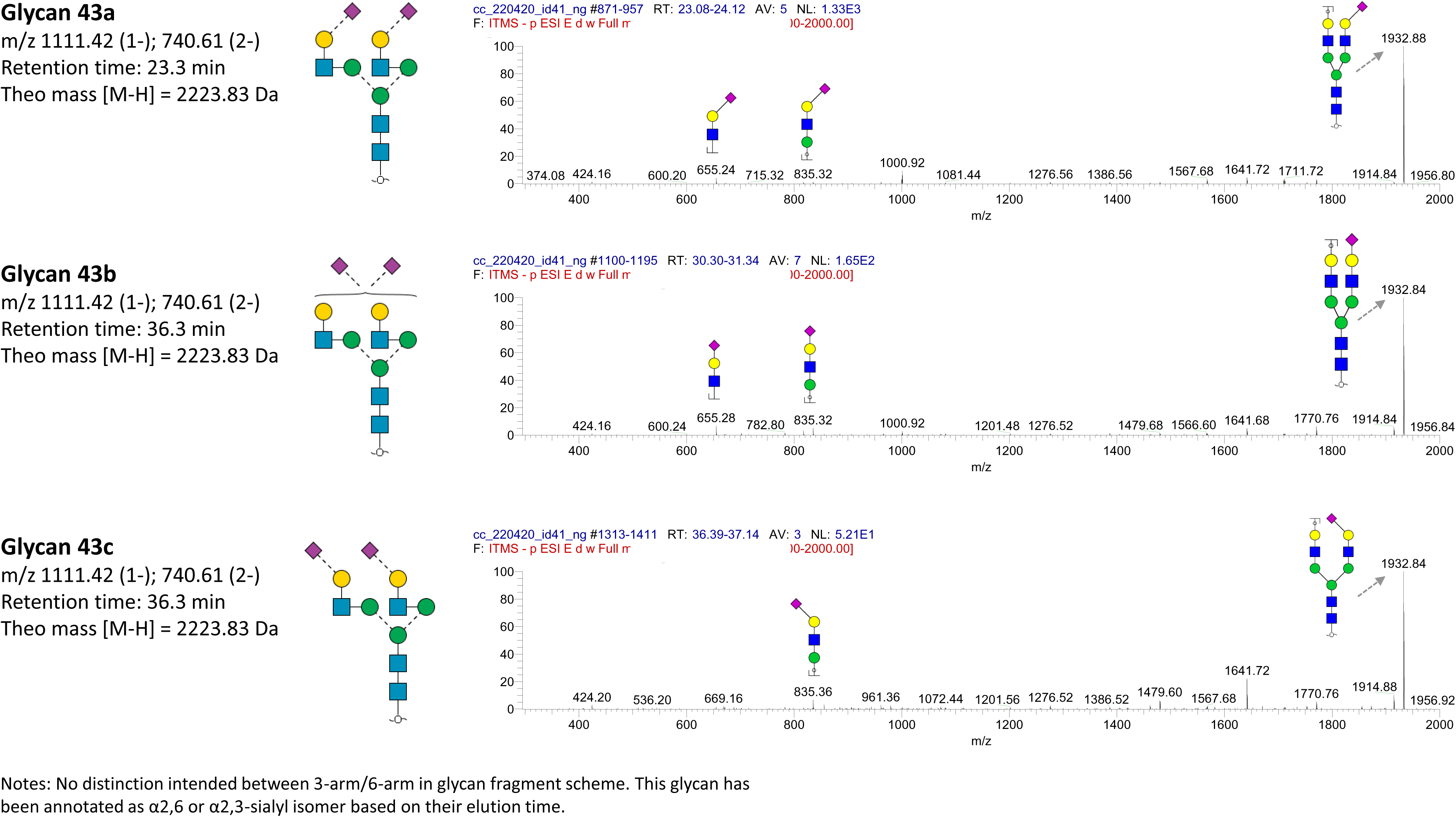
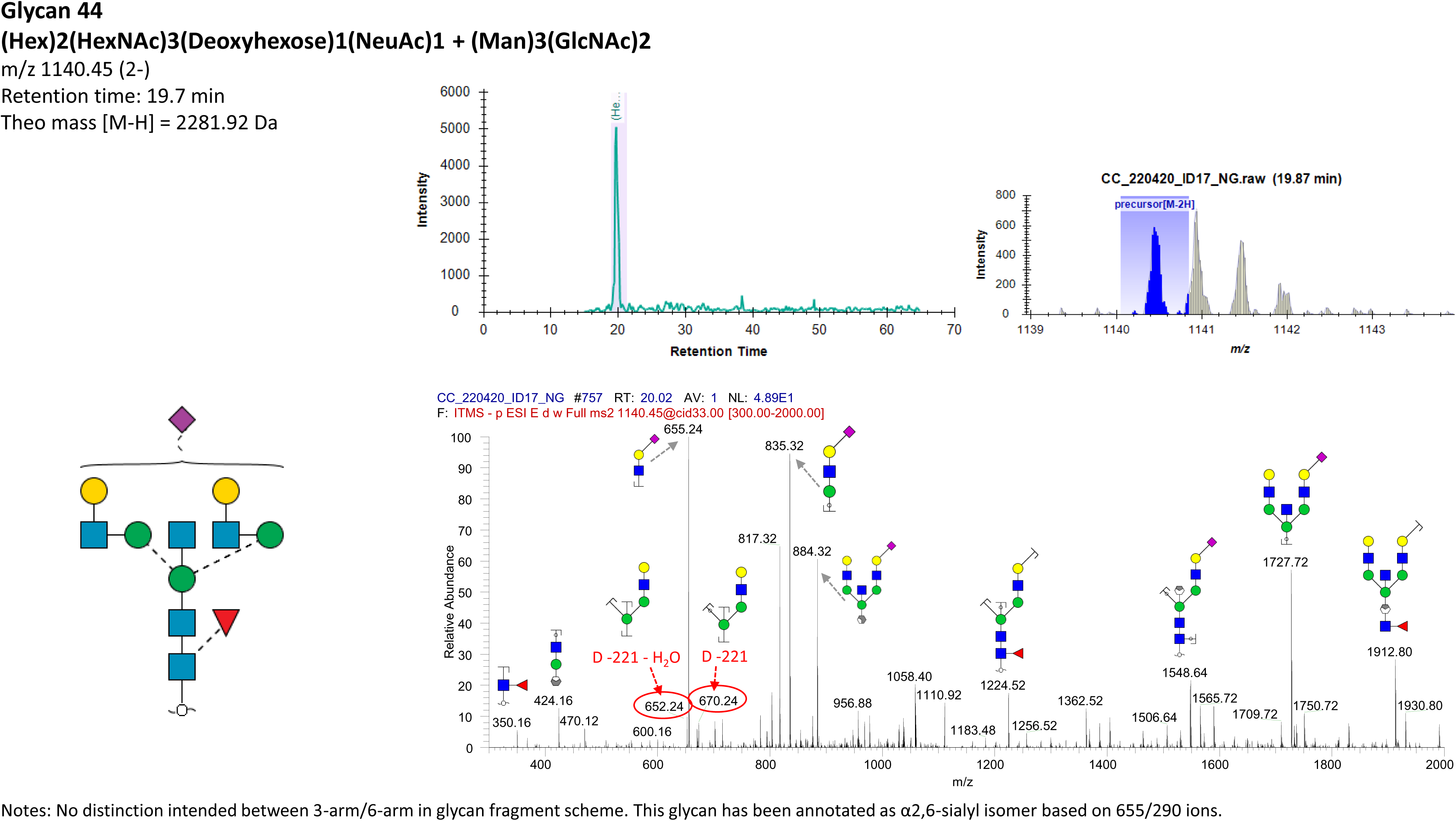
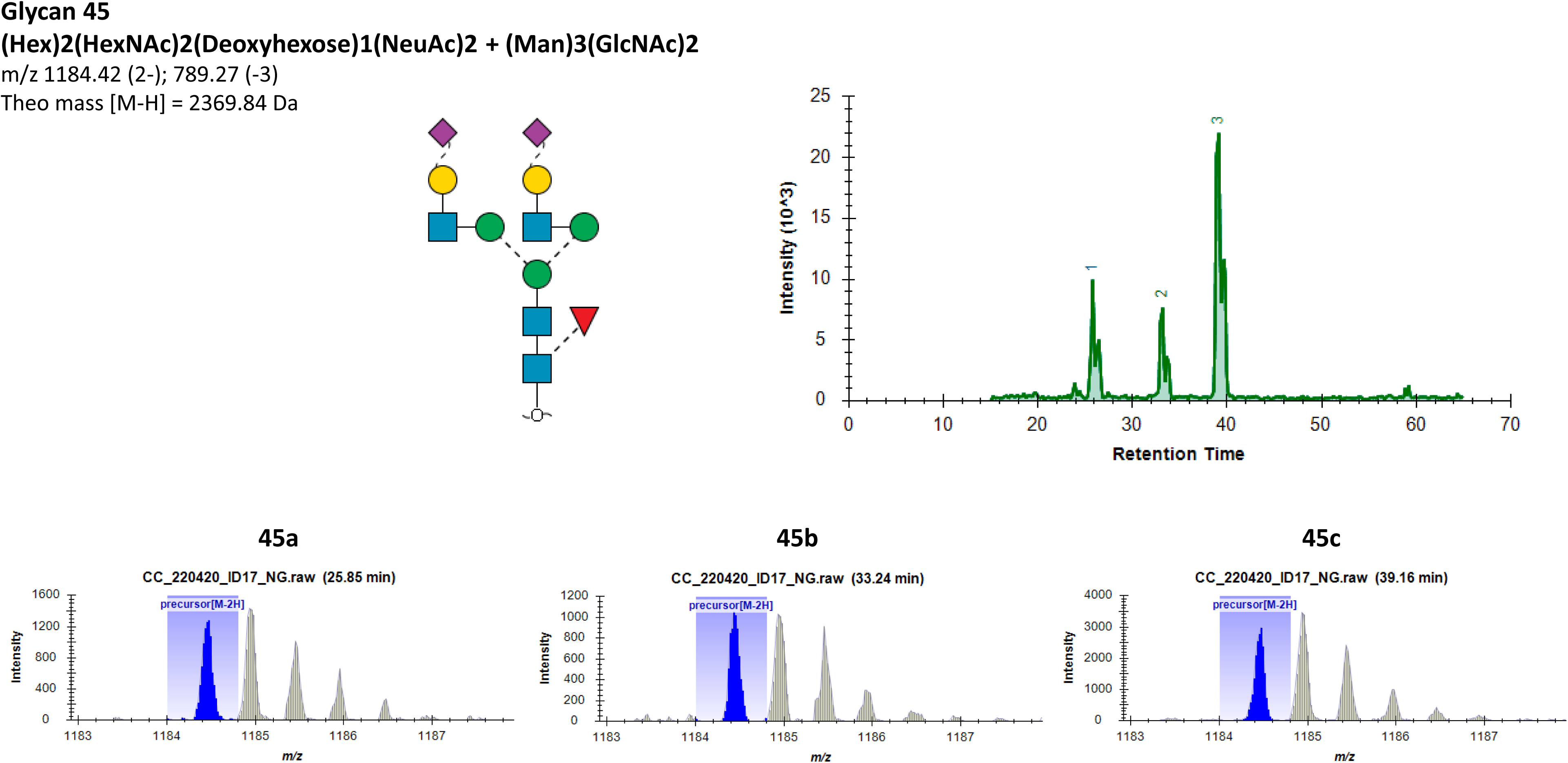
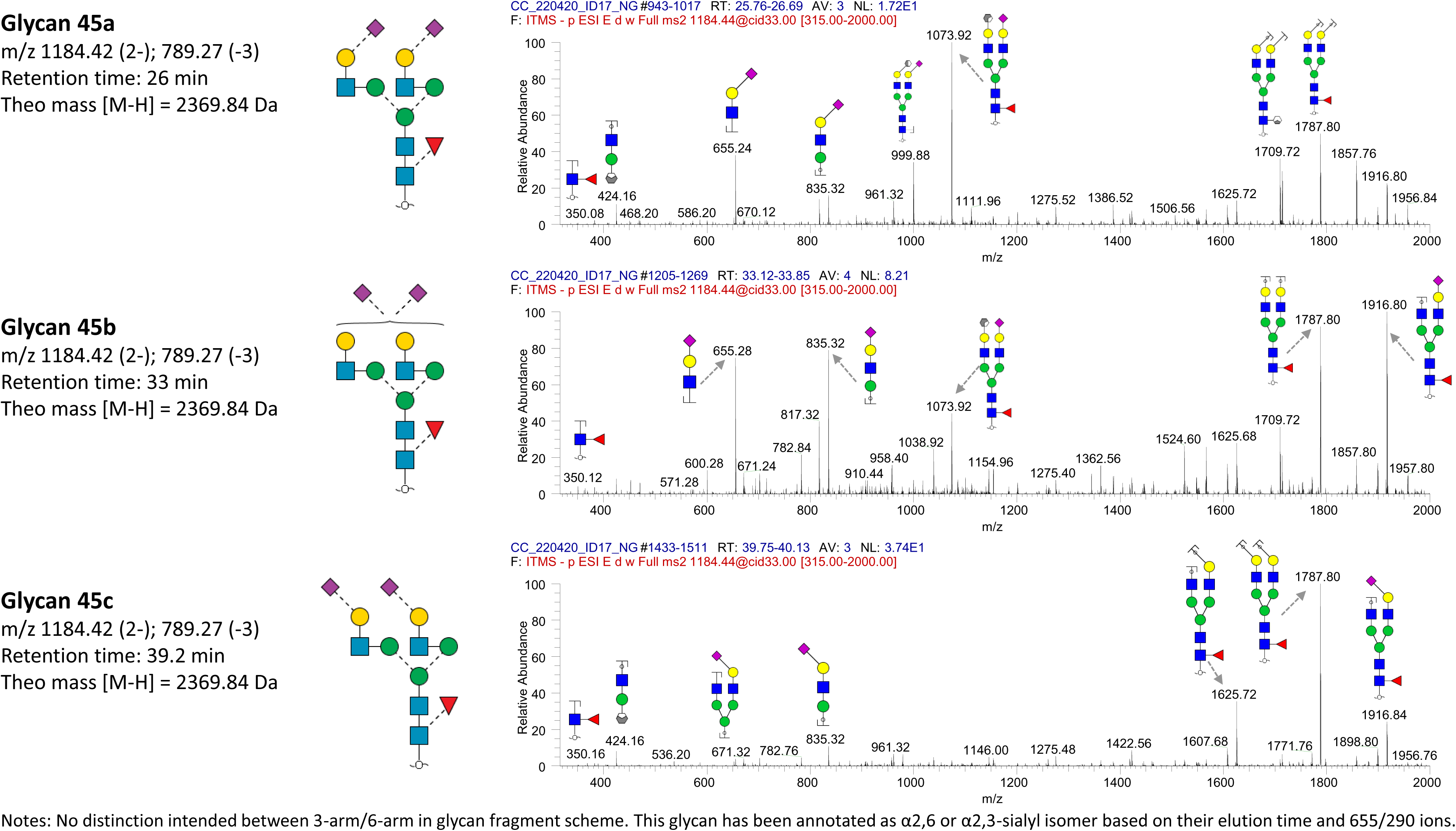
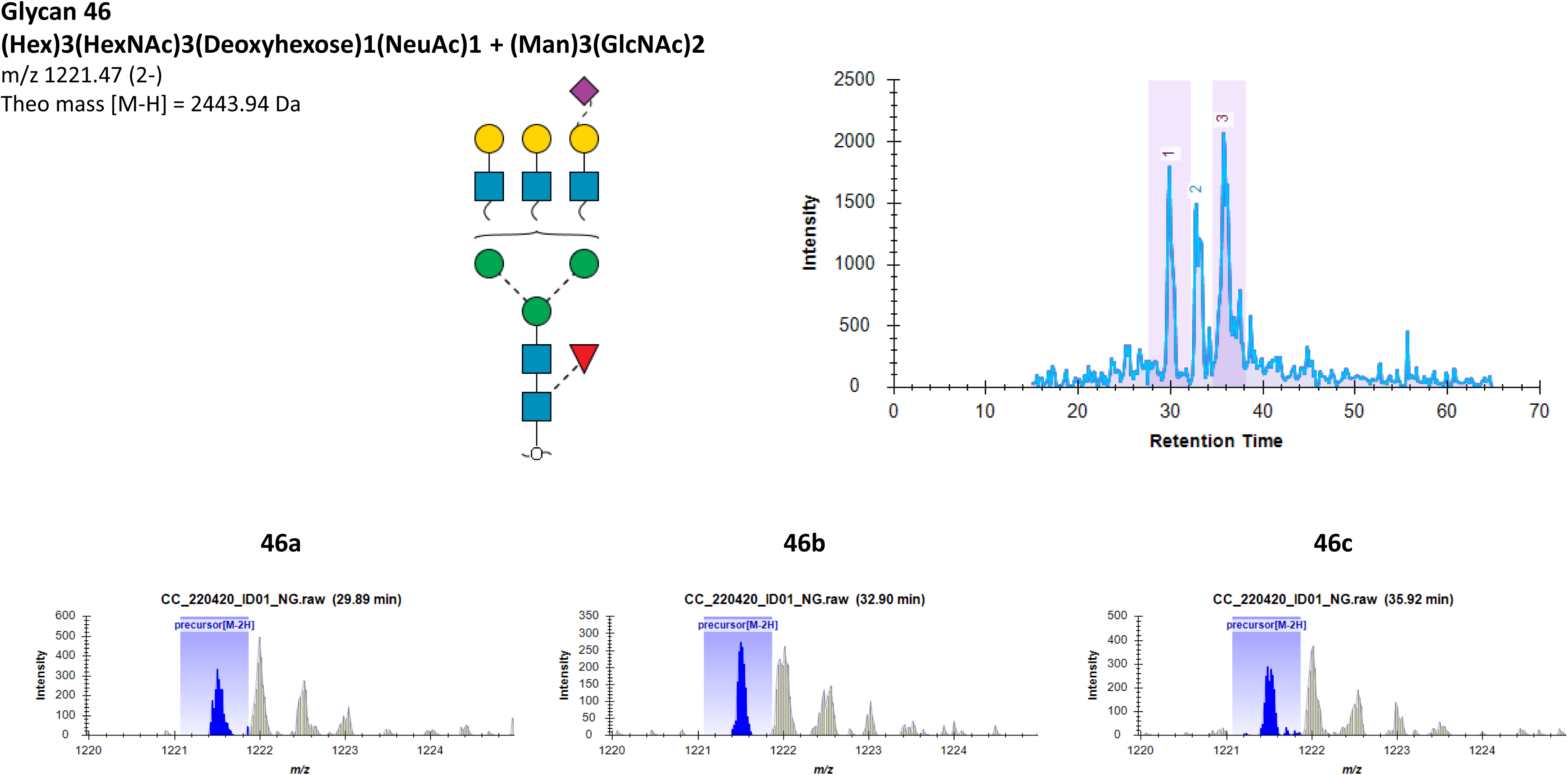
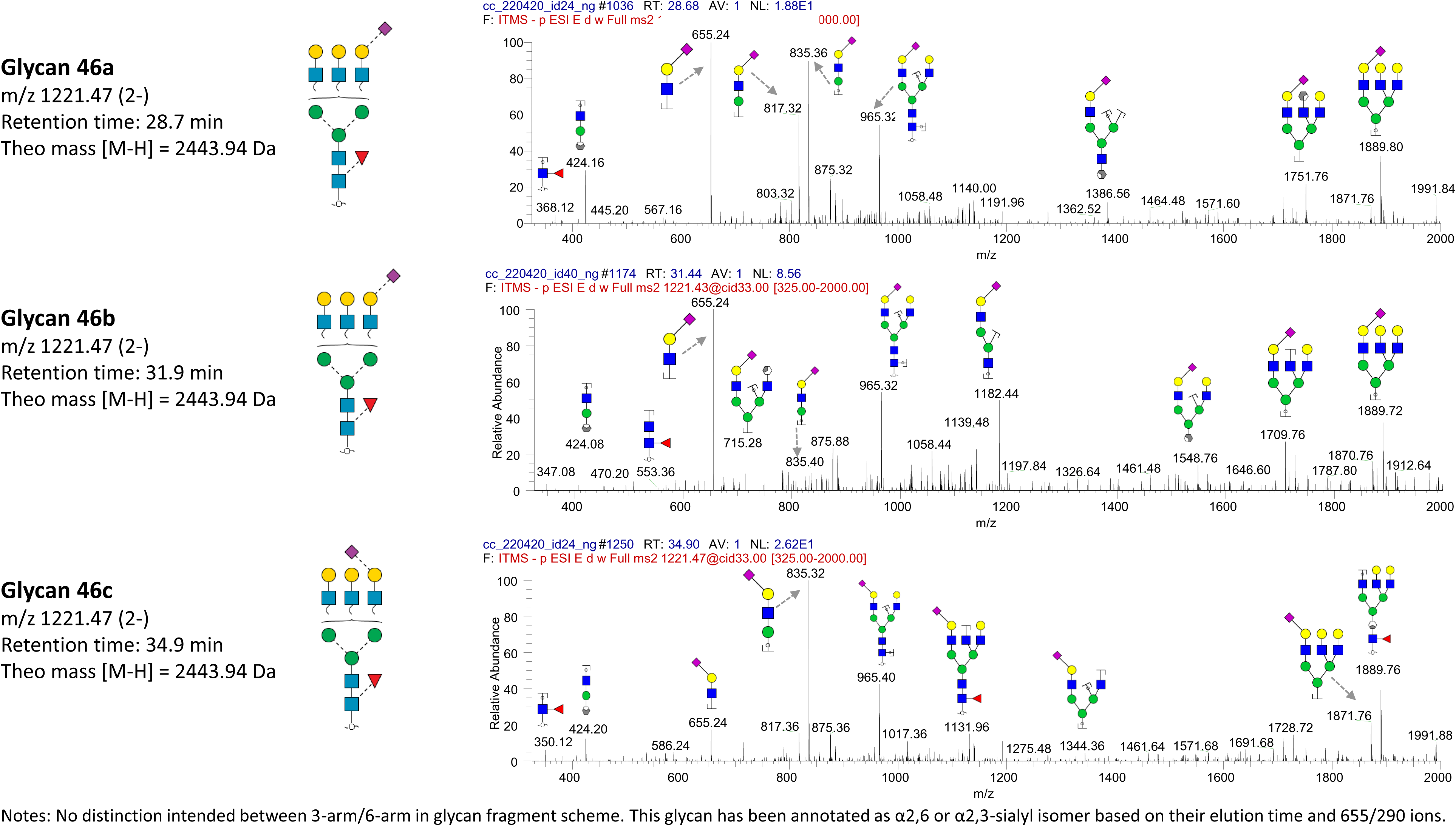
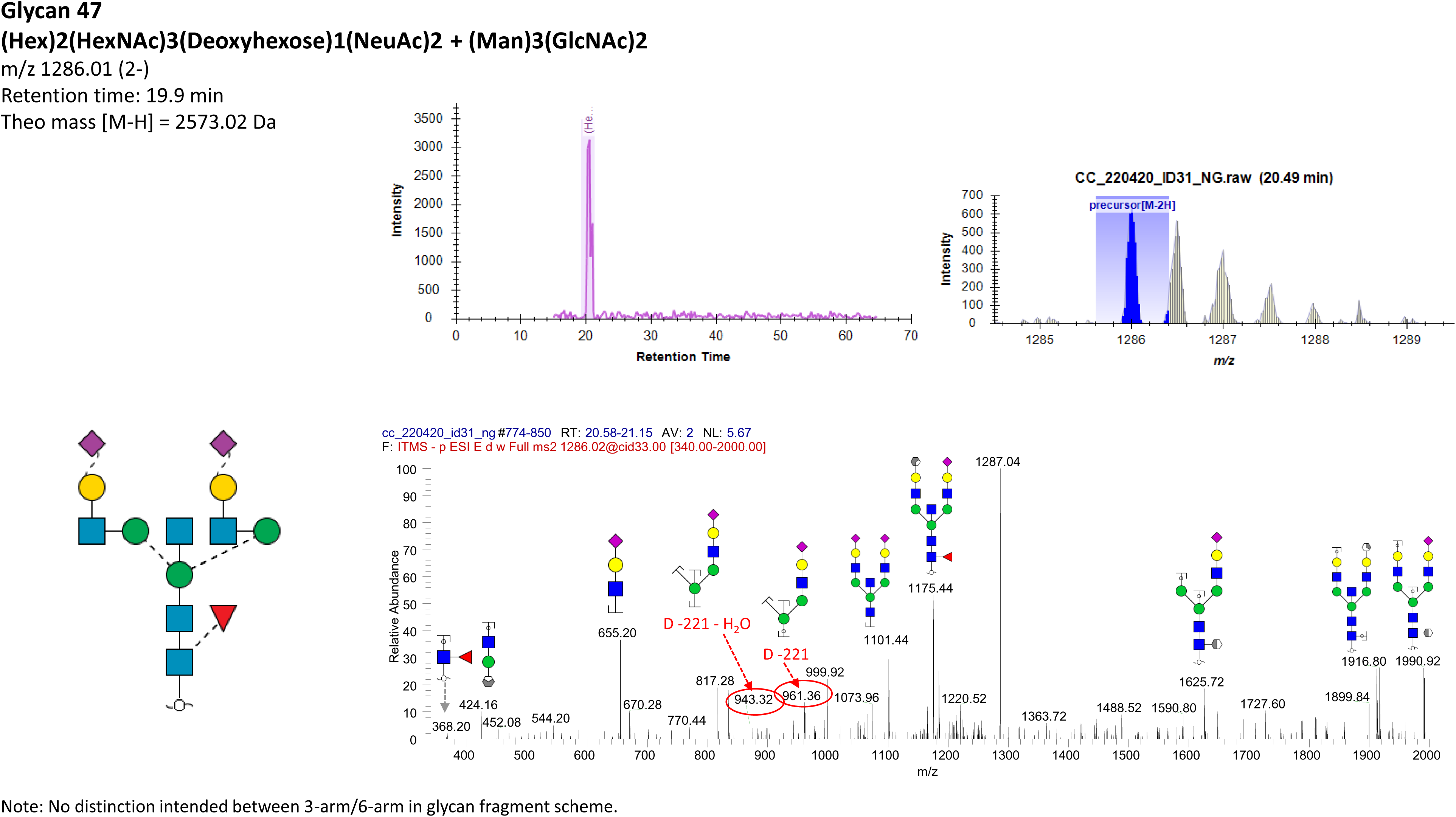
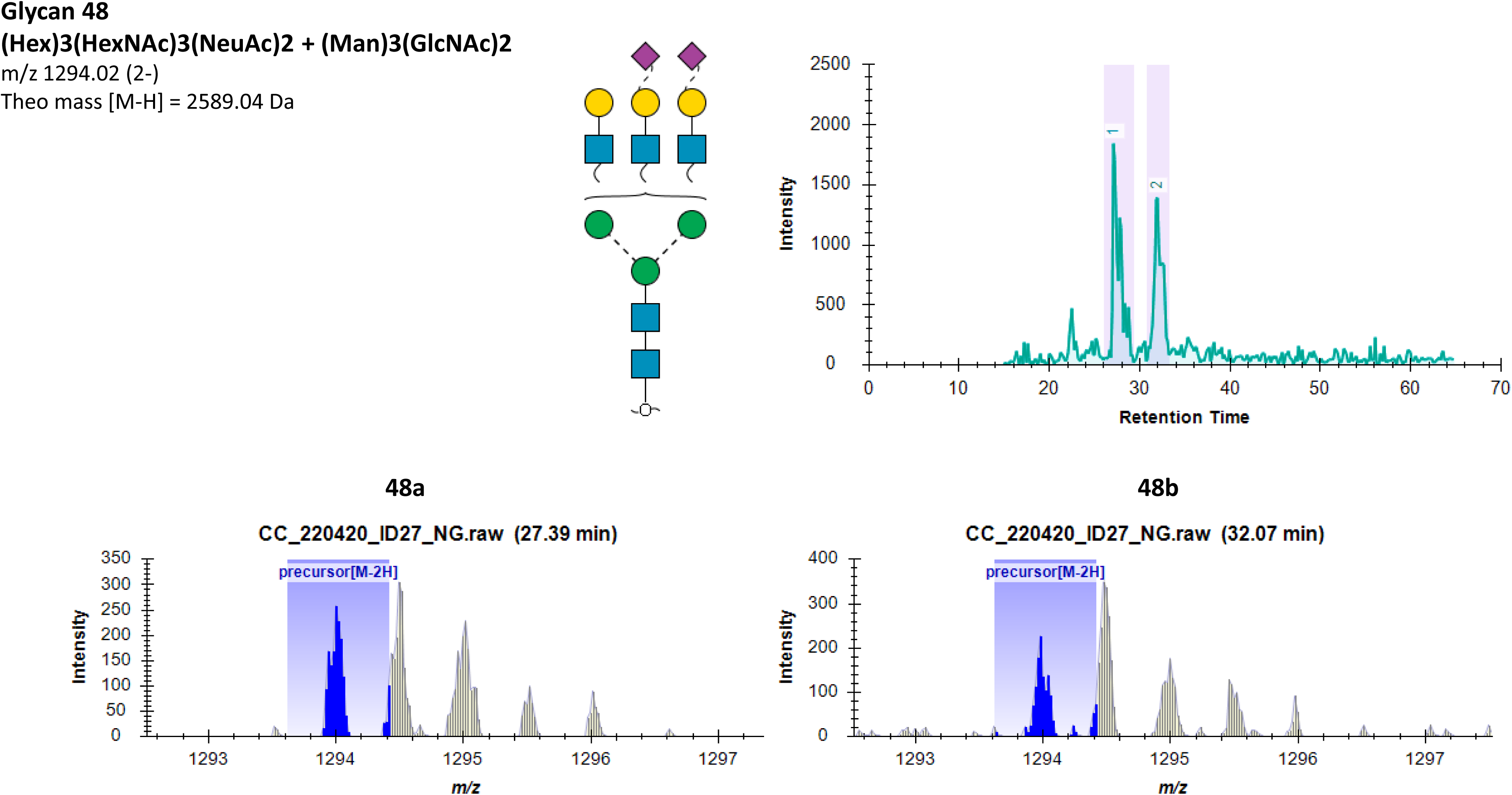
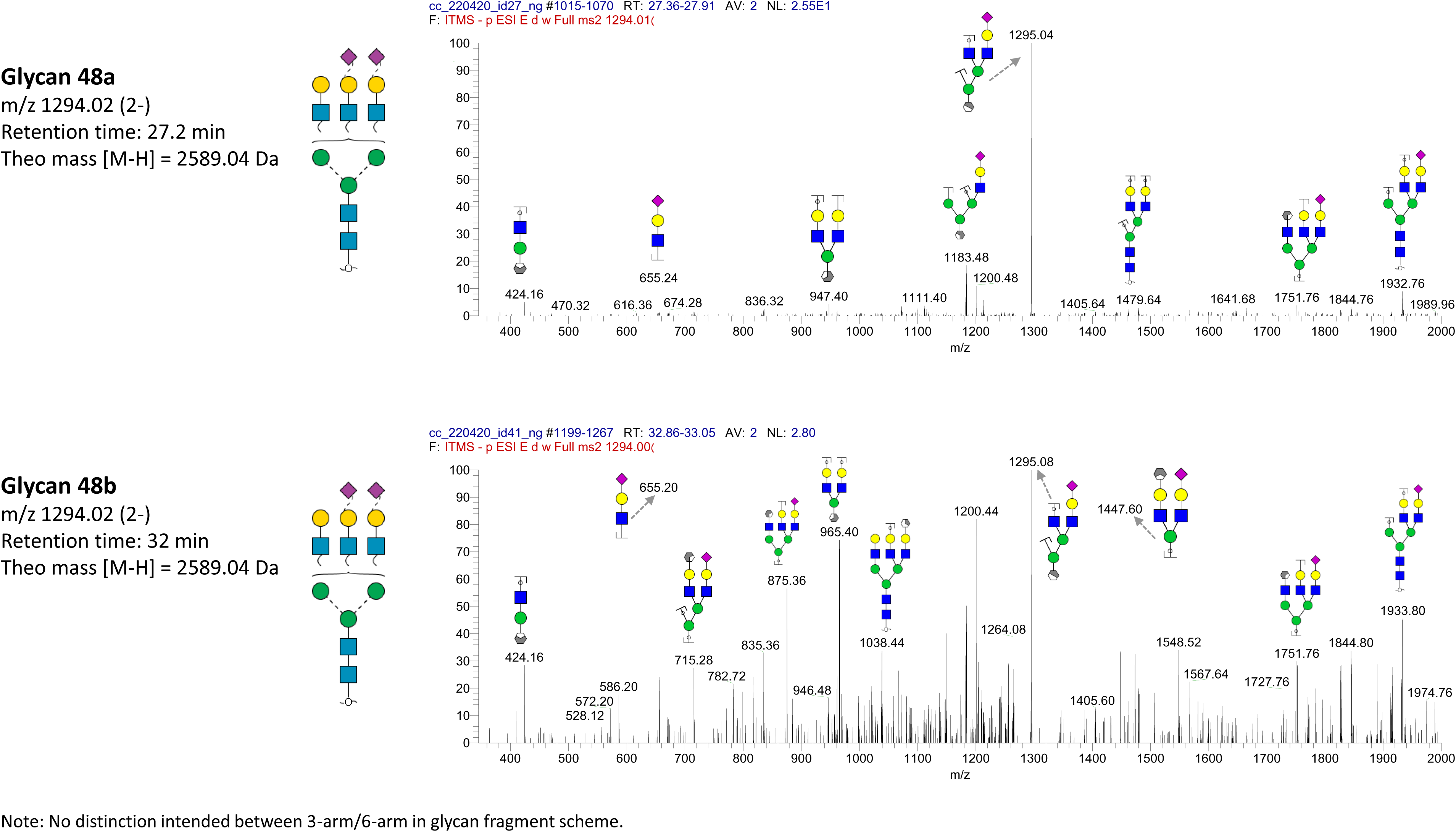
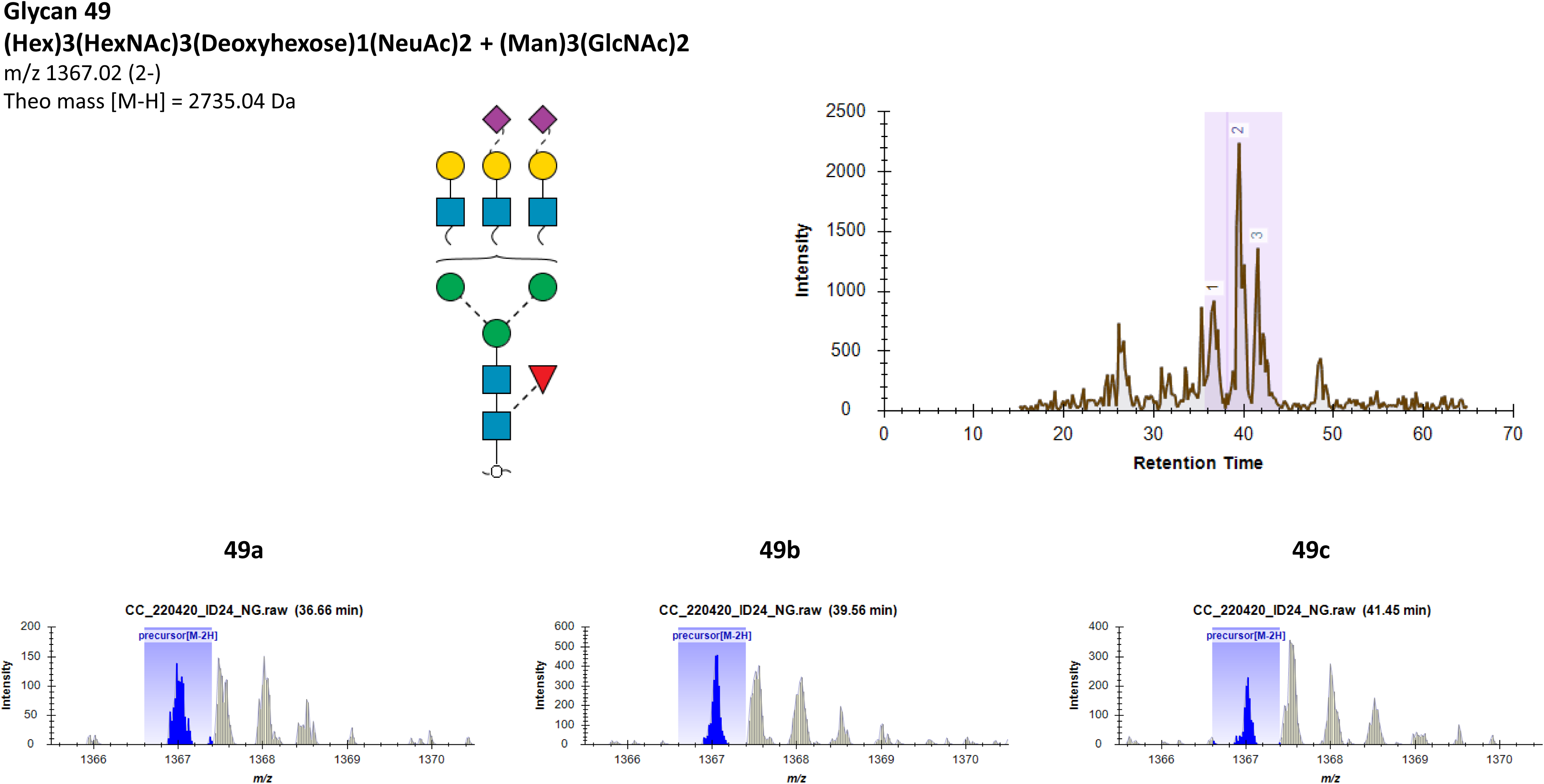
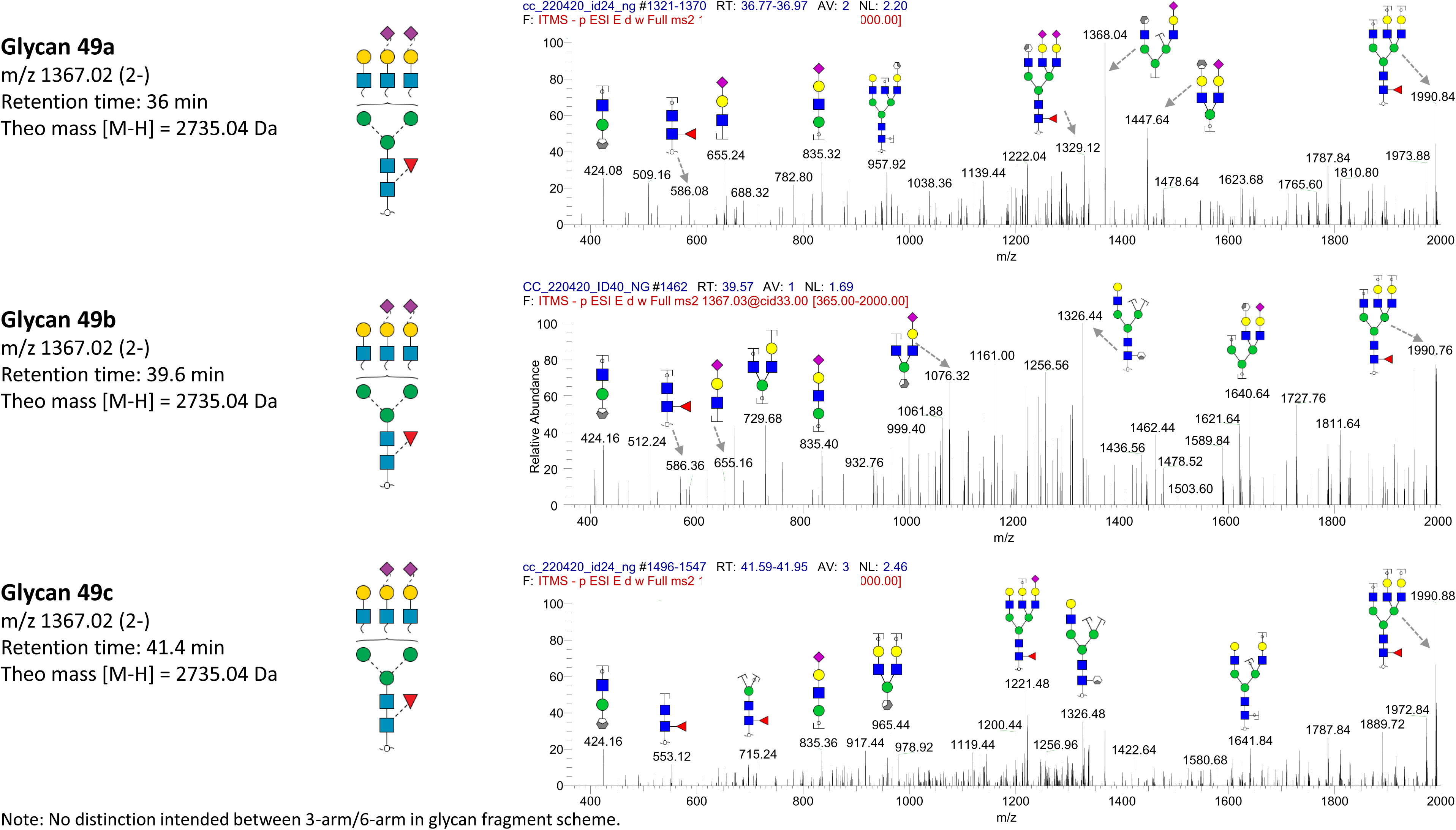
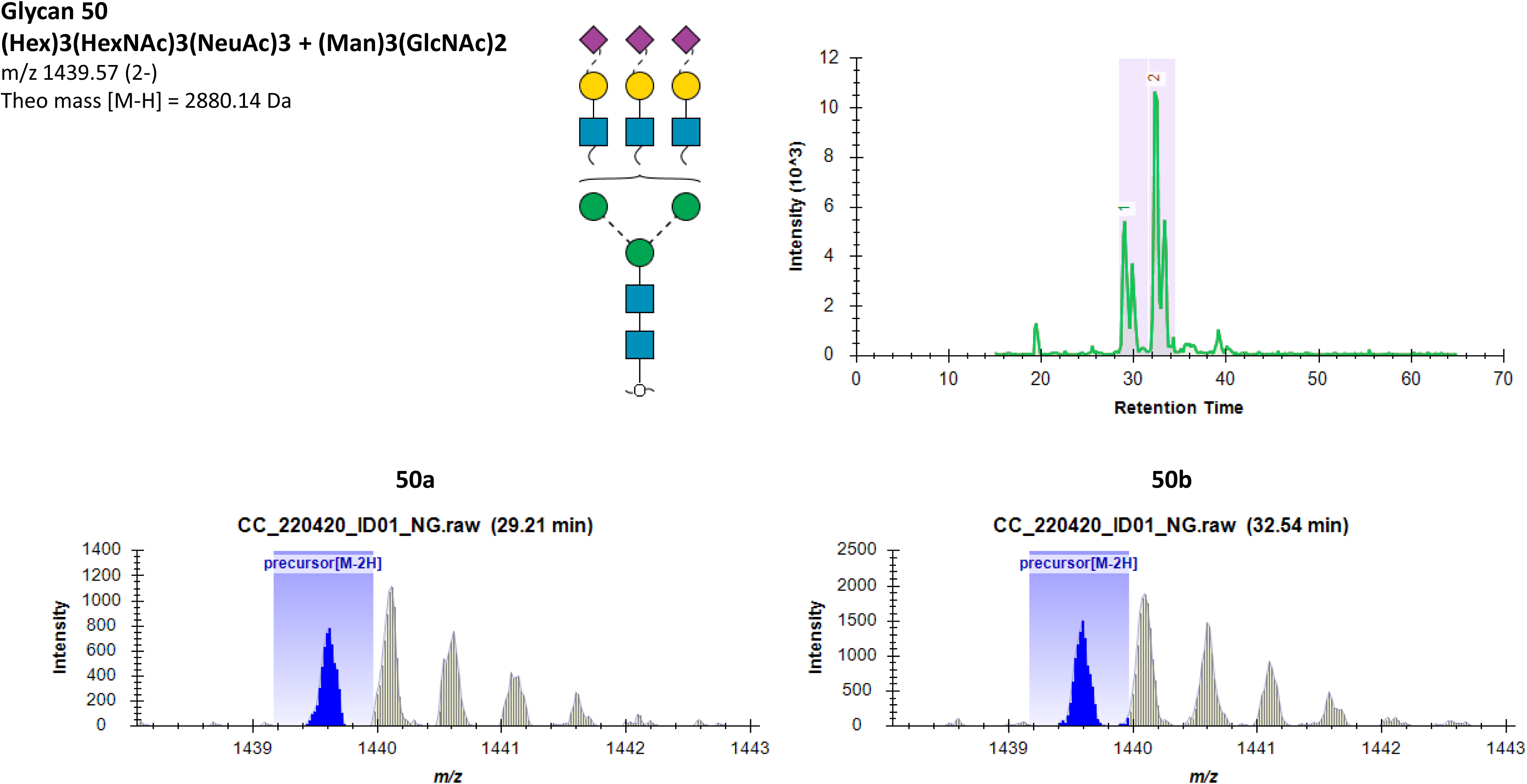
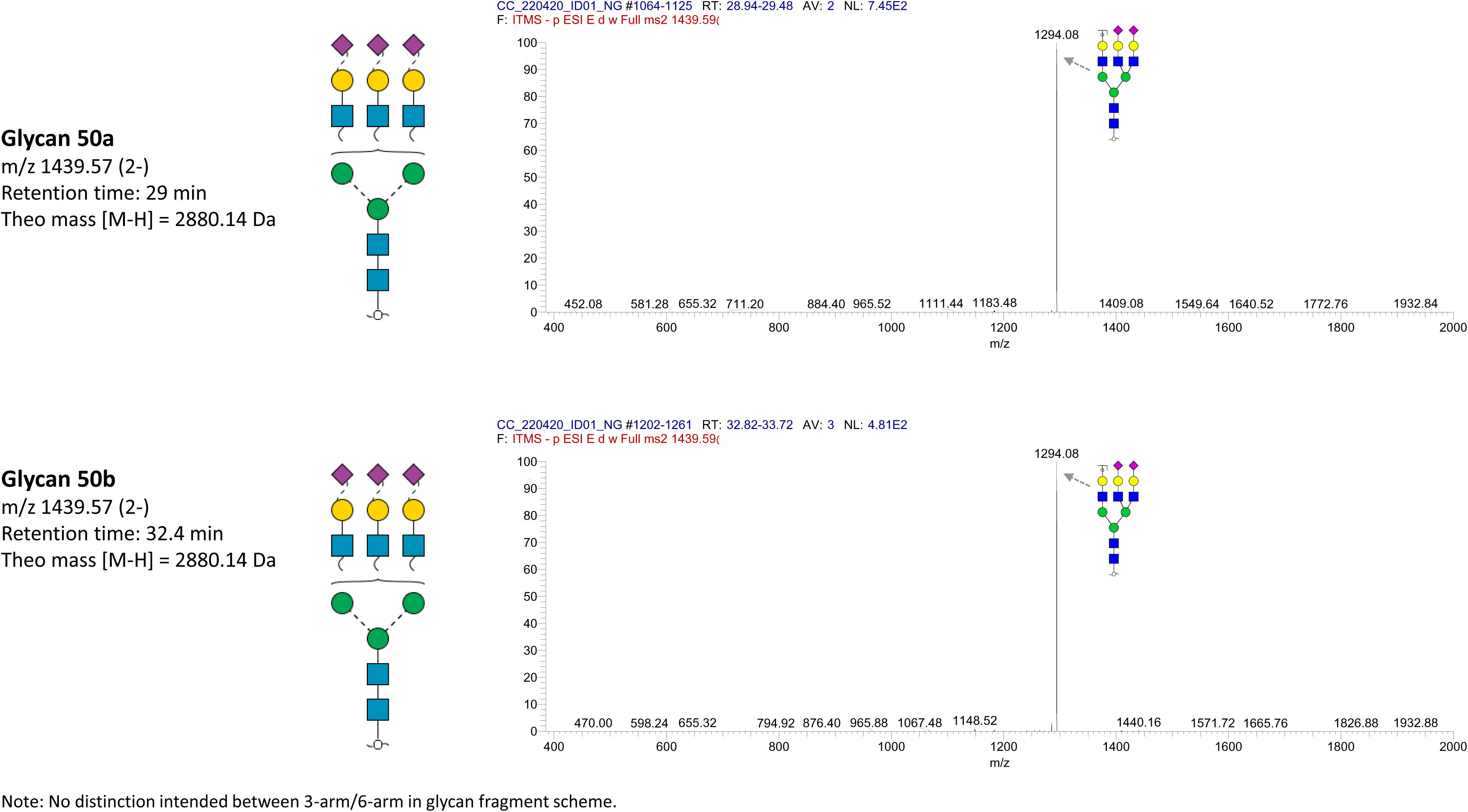
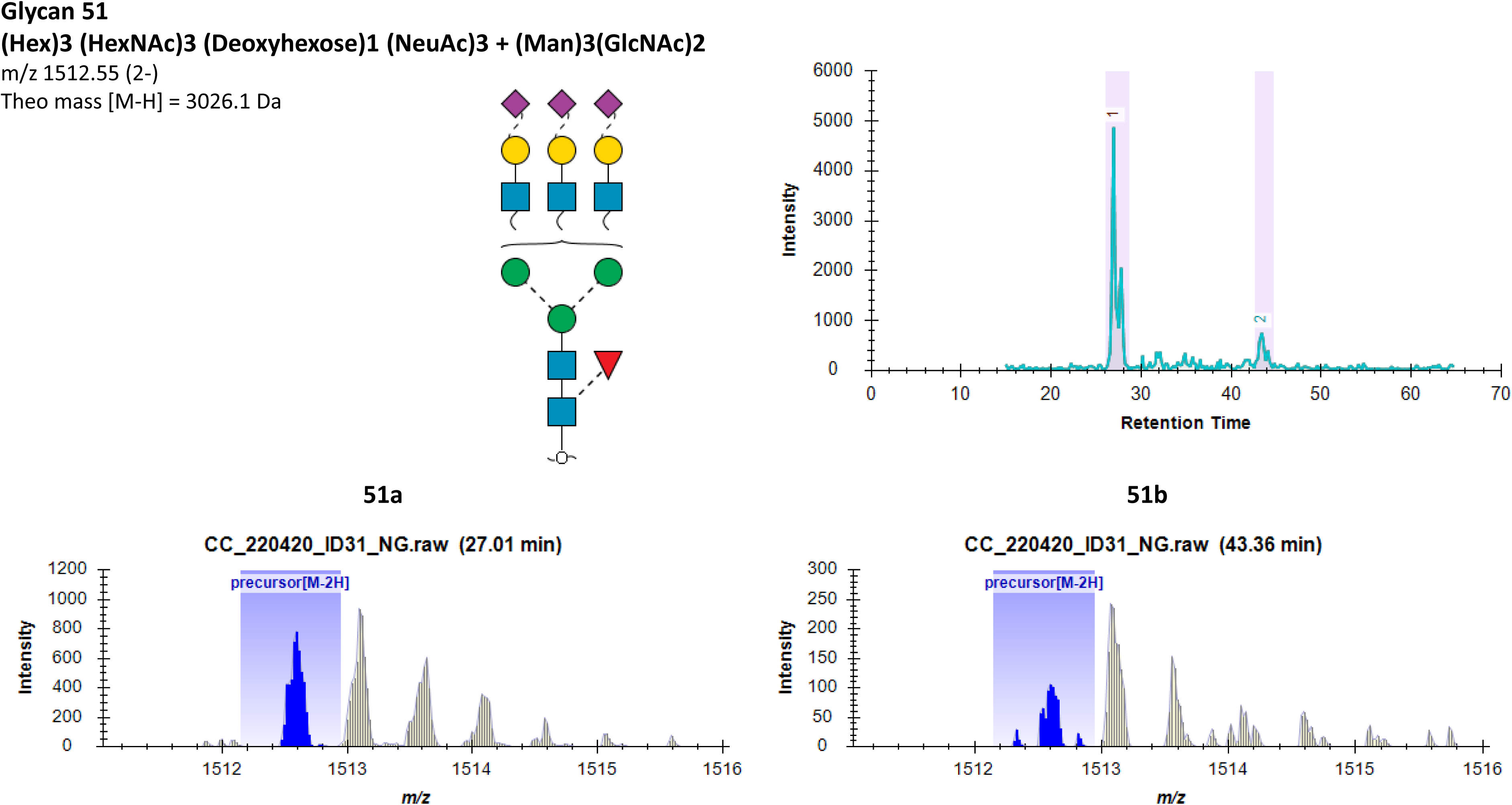
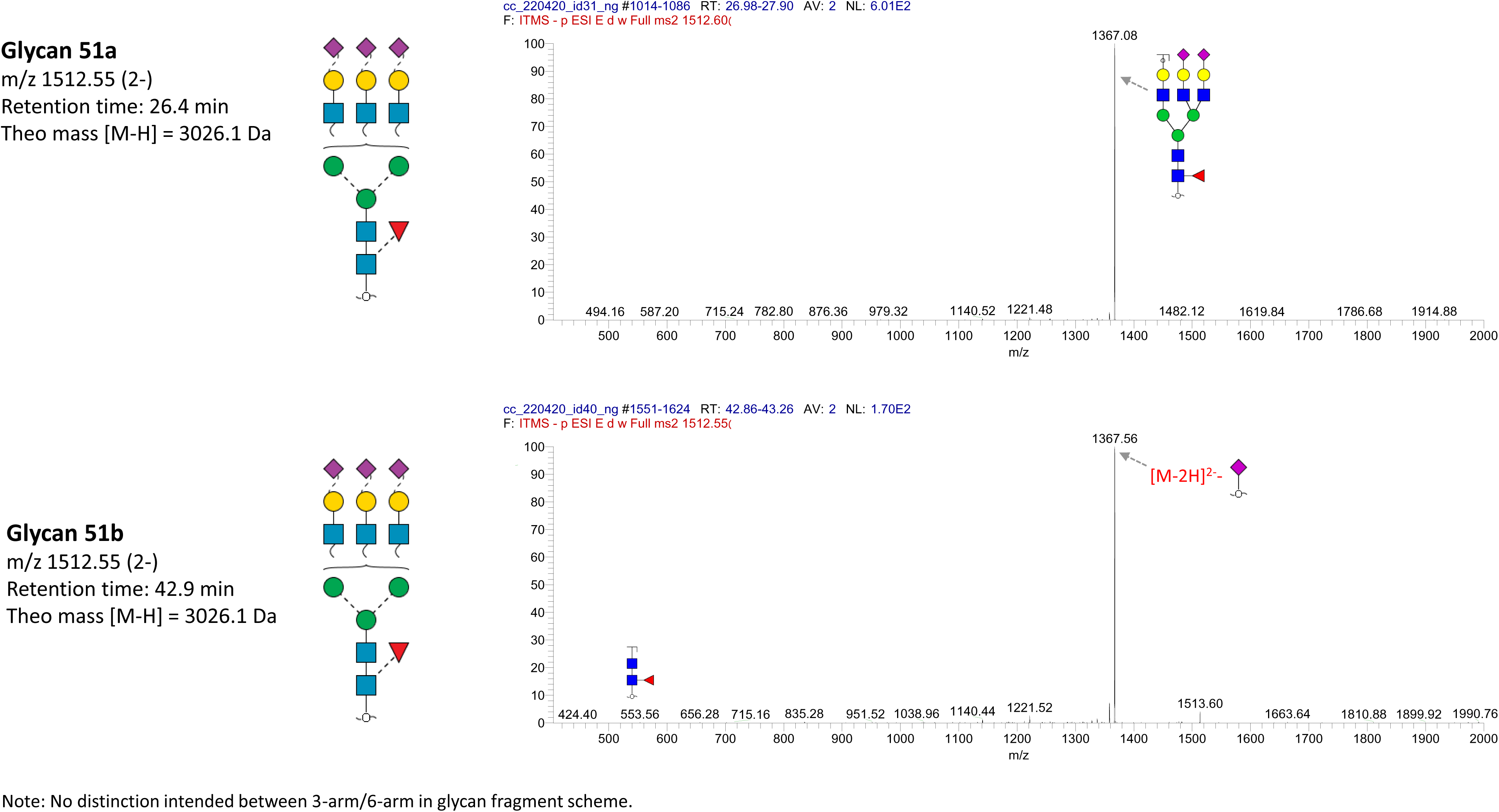
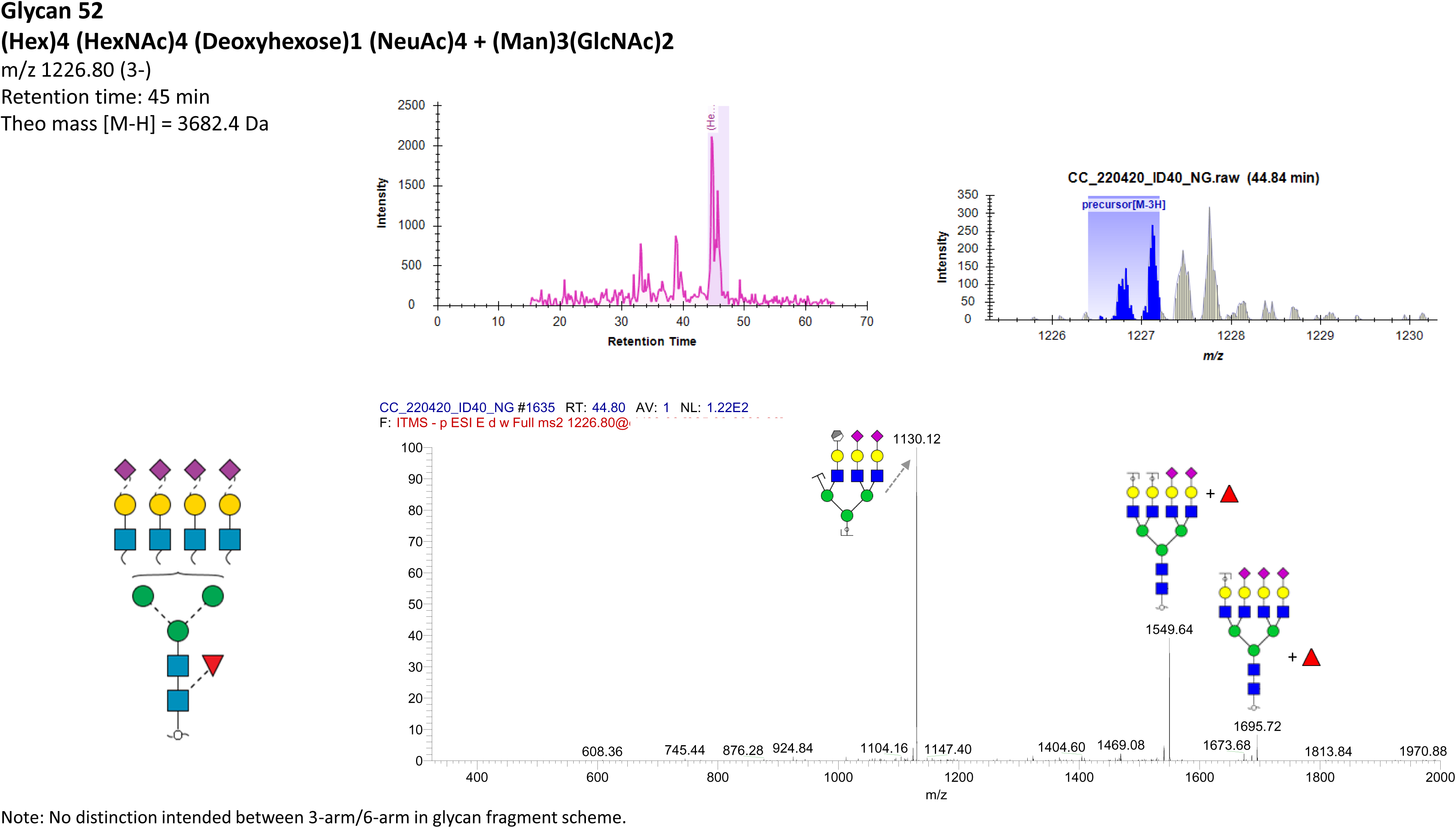

